# Interactions Between SQUAMOSA and SVP MADS-box Proteins Regulate Meristem Transitions During Wheat Spike Development

**DOI:** 10.1101/2020.12.01.405779

**Authors:** Kun Li, Juan M. Debernardi, Chengxia Li, Huiqiong Lin, Chaozhong Zhang, Jorge Dubcovsky

## Abstract

A better understanding of spike development can contribute to improving wheat productivity. MADS-box genes *VRN1* and *FUL2* (*SQUAMOSA*-clade) play critical and redundant roles in wheat spike and spikelet development, where they act as repressors of MADS-box genes of the *SHORT VEGETATIVE PHASE* (*SVP*) clade (*VRT2*, *SVP1* and *SVP3*). Here, we show that wheat *vrt2 svp1* mutants are late flowering, have shorter stems, increased number of spikelets per spike and unusual axillary inflorescences in nodes of the elongating stem. Constitutive expression of *VRT2* resulted in leafy glumes and lemmas, reversion of basal spikelets to spikes, and down-regulation of MADS-genes involved in floral development. Moreover, constitutive expression of *VRT2* enhanced spikelet defects of *ful2*, whereas *vrt2* reduced vegetative characteristics in the spikelets of *vrn1 ful2* mutants heterozygous for *VRN-A1*. These *SVP-SQUAMOSA* genetic interactions were paralleled by physical interactions among their encoded proteins. SVP proteins were able to reduce SQUAMOSA-SEPALLATA interactions in yeast-three-hybrid experiments. We propose that SQUAMOSA-SVP complexes act during the early reproductive phase to promote heading, formation of the terminal spikelet, and stem elongation, but that down-regulation of *SVP* genes is then necessary for the formation of SQUAMOSA-SEPALLATA complexes that are required for normal spikelet and floral development.

## INTRODUCTION

Each year more than 750,000,000 tons of wheat grains are produced around the word providing one fifth of the calories and protein consumed by the human population (FAOSTAT, 2017). These wheat grains are produced in a determinate structure called spike, in which the inflorescence meristem (IM) produces a number of lateral spikelet meristems (SM) that generate sessile lateral spikelets on an axis called rachis, before transitioning itself into a SM to generate a terminal spikelet.

The spikelet is the basic unit of the grass inflorescence (Kellogg, 2001) and, in wheat, it comprises two basal sterile bracts, designated as glumes, followed by an indeterminate number of lateral florets. Each floret has a bract called lemma with an axillary floral meristem (FM) that generates a membranous two-keeled structure called palea, two scales called lodicules that swell to spread the lemma and palea, three stamens and a terminal ovary (Clifford, 1987). In wheat, the SM produces an indeterminate number of FM on an axis called rachilla, with only the most basal florets surviving to set grains (Sakuma et al., 2019). The timing of the transition between the IM to terminal spikelet determines the number of spikelets per spike, and this together with the number of fertile flowers per spikelet determine the maximum number of grains that a spike can produce. Since these are important components of grain yield, a better understanding of their regulatory mechanisms can be useful to engineer more productive wheat plants.

Significant progress has been made to understand the pathways controlling grass inflorescent development, particularly in rice (*Oryza sativa* L.) and maize (*Zea mays* L.), where a complex gene network involving several members of the MADS-box gene family regulates the transitions between different meristem identities (Callens et al., 2018; Wu et al., 2018; Chongloi et al., 2019). MADS-box proteins act as tetrameric complexes and different protein combinations result in the specification of different organ identities, which are well documented in the ABCDE model of flower development in Arabidopsis (*Arabidopsis thaliana* (L.) Heynh.) (Theissen et al., 2016) and rice (Wu et al., 2018). In wheat, there is currently limited knowledge of the role of these genes in spike development. We have recently shown that wheat MADS-box meristem identity genes *VRN1* and *FUL2* from the *SQUAMOSA*-clade are essential for the transition of the IM to a terminal spikelet (Li et al., 2019). In the double *vrn1 ful2* loss-of-function mutants, the IM remains indeterminate and fails to produce a terminal spikelet, whereas the lateral spikelets revert to vegetative structures resembling tillers, some of which have residual flower organs. When a of loss-of-function mutations in *FUL3* (the third member of the *SQUAMOSA*-clade) was combined in a triple *vrn1 ful2 ful3* mutant, the spike lateral meristems generated fully vegetative tillers (Li et al., 2019). These results demonstrate that *VRN1*, *FUL2* and *FUL3* have redundant and essential roles in spikelet development.

In addition to their role in SM identity, these *SQUAMOSA* genes affect wheat heading time and plant height. *VRN1* is a major flowering gene in wheat and its allelic variation determines the need of a long exposure to cold temperatures (vernalization) to flower. Spring wheat varieties carrying the dominant *Vrn1* allele do not have a vernalization requirement, whereas winter wheat varieties the functional but recessive *vrn1* allele require several weeks of vernalization to acquire flowering competence (Yan et al., 2003; Fu et al., 2005; Kippes et al., 2018). The transition from vegetative meristem (VM) to IM is delayed in the *vrn1-*null mutants, further delayed in *vrn1 ful2* and has the greatest delay in *vrn1 ful2 ful3*, which indicates redundant roles of these three genes in the regulation of the initiation of the reproductive phase (Li et al., 2019). Functional redundancy was also observed for plant height, with the triple mutants being shorter than any other mutant combinations (Li et al., 2019).

In this study, we aimed to identify the gene network controlled by *VRN1* and *FUL2* during the early stages of spike development in wheat and, particularly, the genes responsible for the reversion of spikelets to vegetative tillers in the *vrn1 ful2* mutants. By comparing the developing spike transcriptomes of *vrn1 ful2* (spikelets transformed into tillers) and *vrn1* mutants (normal spikes), we identified several up- and down-regulated genes and developmental pathways. Of particular interest was a group of three MADS-box genes of the *SHORT VEGETATIVE PHASE* (SVP) clade, which showed higher expression in the *vrn1 ful2* mutants. These genes include *VEGETATIVE TO REPRODUCTIVE TRANSITION 2* (*VRT2*, synonymous *SVP2*) (Kane et al., 2005), *SVP1* and *SVP3* (Schilling et al., 2020). A previous study in barley (*Hordeum vulgare* L.) showed that constitutive expression of *BM10* and *BM1*, the barley orthologs of *SVP1* and *SVP3*, respectively, results in spikes with reversions to vegetative organs (Trevaskis et al., 2007). A complete list of wheat MADS-box names, synonyms, accession numbers, and orthologs in barley and rice is provided in Supplemental Table 1.

Our interest in the wheat *SVP* genes was also based on the known roles of the Arabidopsis homologs *SVP* and *AGL24*. *SVP* acts as a repressor of the floral transition (Hartmann et al., 2000); whereas *AGAMOUS-LIKE 24* (*AGL24*) acts as a promoter of flowering (Yu et al., 2002; Michaels et al., 2003). In spite of their opposite roles in the VM to IM transition, *SVP* and *AGL24* have overlapping and redundant effects on floral meristem identity specification during the reproductive phase (Gregis et al., 2013), where they contribute to the repression of B-class and C-class flowering genes through direct control of *SEP3* (Liu et al., 2009). The *svp agl24* double mutant shows mild floral defects at basal positions of the inflorescence, but the triple mutant including the *ap1-12* weak mutation in *APETALLA 1* (a homolog of wheat *VRN1/FUL2/FUL3*) showed enhanced floral defects (Gregis et al., 2006). When *ap1-12* was replaced by the strong *ap1-10* mutation, the inflorescences continuously produced IMs, indicating an interaction between *SQUAMOSA* and *SVP* genes to specify the FMs (Gregis et al., 2008). The wheat VRT2 protein has been shown to bind to a CArG box in the *VRN1* promoter (Kane et al., 2007; Dubcovsky et al., 2008; Xie et al., 2019), suggesting also possible interactions between these two groups of MADS-box genes in wheat.

In this study, we explore the genetic interactions between the *SQUAMOSA* and *SVP* genes and their effects on wheat spike development. Using crosses between *vrt2* and *svp1* mutants, transgenic plants constitutively expressing *VRT2* and crosses with *vrn1 ful2* mutants, we show complementary and overlapping roles between genes in the *SVP*- and *SQUAMOSA*-clades in the early reproductive phase, but antagonistic effects later during spikelet and floral development. We show that constitutive expression of *VRT2* down-regulates multiple MADS-box genes of the *SEPALLATA*-clade involved in floral development and promotes vegetative traits in spikelets. Finally, we describe a complex network of protein interactions among wheat proteins from the SQUAMOSA, SVP and SEPALLATA clades that suggests that the down-regulation of *SVP* genes by SQUAMOSA proteins is required to facilitate interactions between SQUAMOSA and SEPALLATA proteins, which are important for normal spike and floral development.

## RESULTS

### Quant-Seq analysis of developing wheat spikes in *vrn1* and *vrn1 ful2* mutants reveals genes regulated by *VRN1* and *FUL2*

During early spike development, the IM in the *vrn1* mutant produces lateral meristems that acquire SM identity and develop into spikelets whereas the IM in the *vrn1 ful2* double mutant produces lateral vegetative meristems that develop into tillers (Figure 1A). To identify the genes and pathways that repress the vegetative program and activate the spikelet identity program, we compared the transcriptomes of developing apices of these two mutants at four developmental stages covering the early steps of spike development: vegetative (VEG), double-ridge (DR), post-double-ridge (PDR) and terminal spikelet (TS) (Figure 1A). The average number of unique reads per sample and other transcriptome statistics are summarized in Supplemental Table 2.

**Figure 1.**
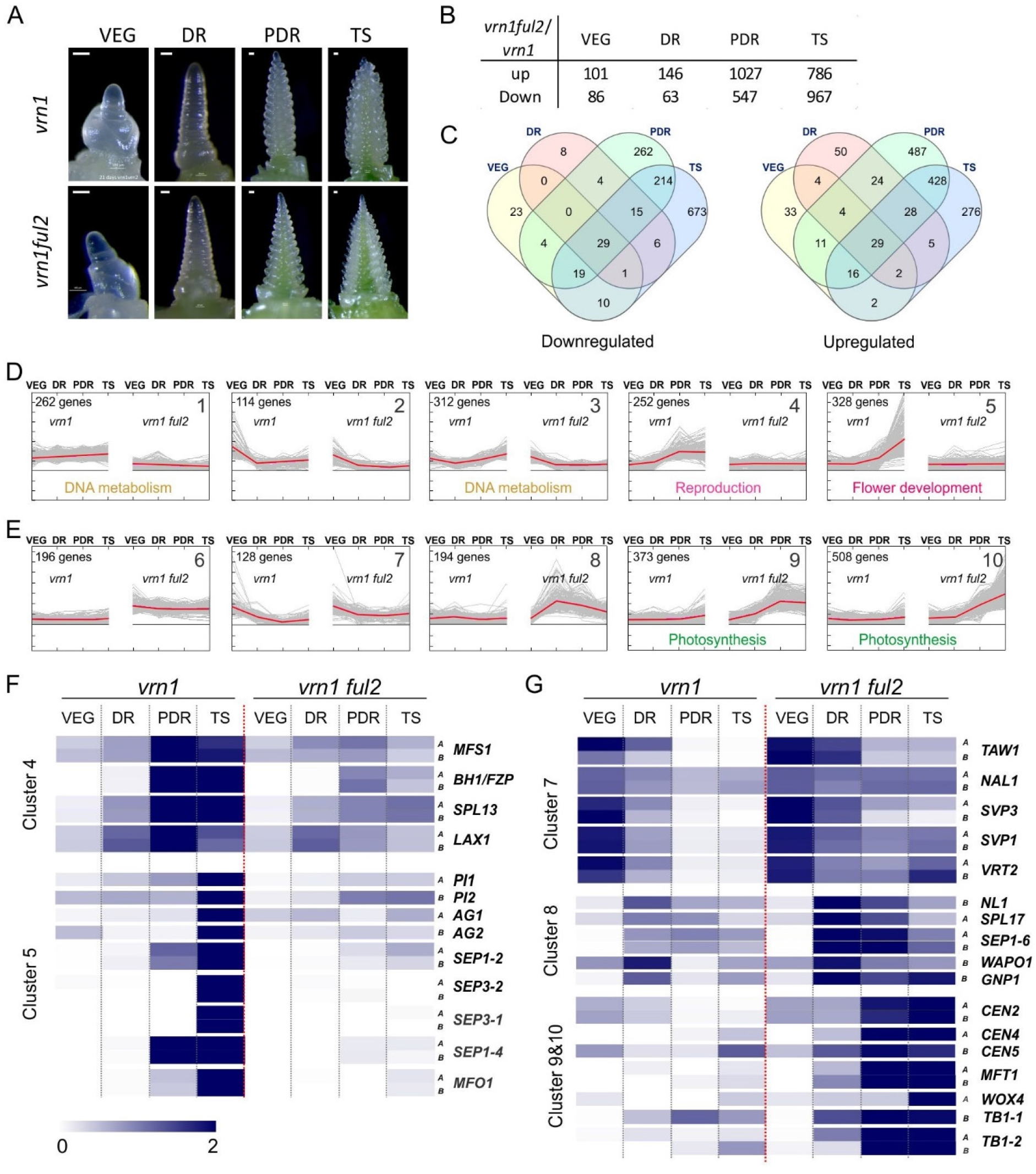
Differentially expressed genes in developing spikes of *vrn1* and *vrn1 ful2* mutants. **(A)** Representative pictures of vegetative (VEG) and reproductive apices (DR = double ridge; PDR = post double ridge; TS = terminal spikelet) from *vrn1* (above) and *vrn1 ful2* (below) collected for Quant-seq analysis. **(B)** Number of DEGs (differentially expressed genes) identified at each stage. **(C)** Venn diagrams showing the non-redundant DEGs in the intersections of the different developmental stages. Down-regulated (left) and up-regulated (right). **(D-E)** Mean normalized expression pattern for clusters of non-redundant DEGs in *vrn1ful2* and *vrn1* samples. The red lines represent the mean of all DEGs in the cluster, while light grey lines show individual DEGs. Significantly enriched GO terms were significant only for cluster 1, 3, 4, 5, 9, 10 and are presented below the curves. **(F)** Heat map showing mean normalized expression of selected genes for clusters 4 and 5. **(G)** Heat map showing the relative expression pattern of selected genes for clusters 7, 8, 9 and 10. Scale 0-2 indicates mean normalized expression. A full list of DEGs is available as Supplemental File 1, and a description of DEGs with known functions in inflorescence development are described in Supplemental Table 3.

In the comparison between developing spikes of *vrn1* and *vrn1 ful2* mutants at the VEG and DR stages, we found 187 Differentially Expressed Genes (DEGs, 86 down-regulated and 101 up-regulated) and 209 DEGs (63 down-regulated and 146 up-regulated), respectively (Figure 1B). These numbers greatly increased in the PDR and TS stages to 1,574 and 1,753 DEGs, respectively (Figure 1B), reflecting drastic differences between the developmental courses of the reproductive spikelets in *vrn1* and the lateral tillers in the spikes of *vrn1 ful2*.

We then performed a cluster analysis of the 1,399 up-regulated and 1,268 down-regulated non-redundant genes (Figure 1C) based on their expression profiles across the four developmental stages. This analysis resulted in 5 clusters for each of the two sets, which included at least 10% of the up- or down-regulated genes (Figure 1D and E). Clusters 4 and 5 included genes that were up-regulated in *vrn1* but not in *vrn1 ful2* at PDR (cluster 4) or TS (cluster 5). A GO analysis of these clusters revealed an enrichment of genes involved in early reproductive development in cluster 4 and flower development in cluster 5, including four genes of the *SEPALLATA* clade (Figure 1D and F). These results agree with the reproductive fate of the *vrn1* spike lateral meristems. Additional genes from these clusters with known roles in inflorescence development are described in Supplemental Table 3.

We observed the opposite profiles in clusters 9 and 10, which included genes up-regulated in *vrn1 ful2* but not in *vrn1* at PDR (cluster 9) or TS (cluster 10) (Figure 1E and G). A GO enrichment analysis of these clusters revealed an enrichment for genes involved in photosynthesis (Figure 1E), which agrees with the vegetative fate of the *vrn1 ful2* spike lateral meristems. This cluster also includes florigen antagonists *CEN2*, *CEN4* and *CEN5* (Figure 1G). Cluster 8 showed a peak at the DR stage and included several genes previously shown to be involved in the regulation of SNS (Figure 1G, Supplemental Table 3).

Since we were particularly interested in negative regulators of spikelet meristem identity, we also analyzed genes from cluster 7, which were highly down-regulated at PDR and TS in *vrn1* but not in *vrn1 ful2* (Figure 1G, Supplemental Table 3). This cluster included three MADS-box genes of the *SVP-*clade, confirming a previously published qRT-PCR result showing significantly lower *VRT2*, *SVP1* and *SVP3* transcript levels in *vrn1* relative to *vrn1 ful2* at the TS stage (Li et al., 2019). Of the three wheat genes in the *SVP*-clade, we prioritized the characterization of *VRT2* and *SVP1* because of their higher expression levels relative to *SVP3* at the PDR and TS stages (Figure 1G), and also because of their closer evolutionary relation relative to *SVP3* (Supplemental Figure 1).

### Identification and combination of loss-of-function mutants for *VRT2* and *SVP1* in tetraploid wheat

We selected truncation mutations for the A and B genome homeologs of *VRT2* and *SVP1*, which are summarized in Figure 2A and B, respectively (for more detail see Materials and Methods). To generate the *VRT2* loss-of-function mutant, designated hereafter as *vrt2*, we combined the premature stop codon mutation Q125* in the A-genome homeolog (*vrt-A2*) with a splice site mutation in the B-genome homeolog (*vrt-B2*) (Figure 2A). This *vrt-B2* mutation results in splice variants with premature stop codons or a large deletion in the middle of the protein (Supplemental Figure 2). To generate the *SVP1* loss-of-function mutant, designated hereafter as *svp1*, we intercrossed an *svp-A1* mutant carrying a splice site mutation that generates splice variants with premature stop codons (Supplemental Figure 2) with a *svp-B1* mutant carrying the premature stop codon Q99* (Figure 2B). We generated PCR markers for each of these four mutations to trace them in the different crosses and backcrosses (Supplemental Table 4).

**Figure 2.**
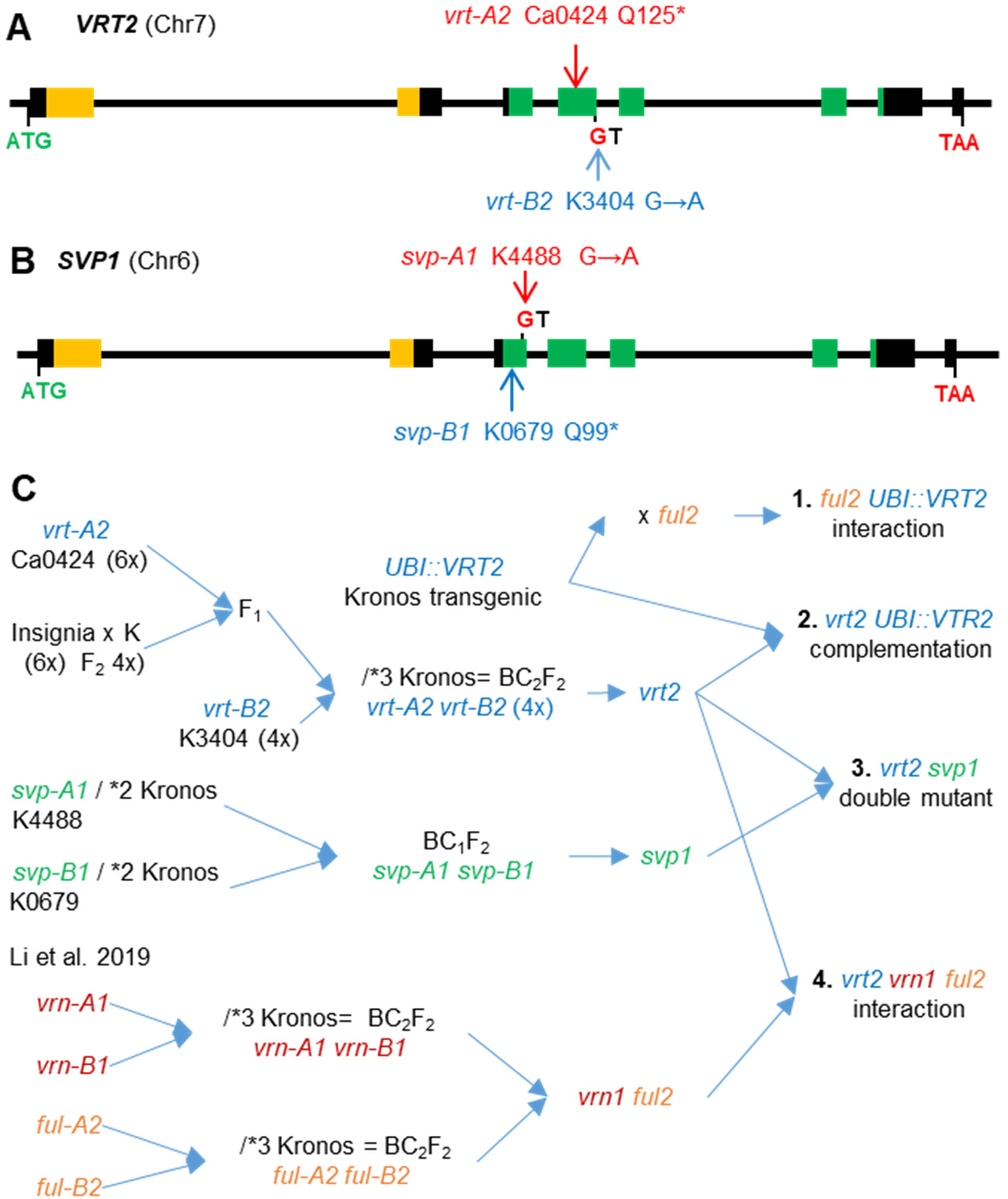
Selected *VRT2* and *SVP1* mutations and their effects on the encoded MADS-box proteins. **(A-B)** Location of the selected mutations in the gene structure diagram: exons are represented by rectangles, with those in orange encoding for the MADS domain and those in green for the conserved K domain. Ca= hexaploid wheat Cadenza and K= tetraploid wheat Kronos. GT= mutated splice site. **(A)** *VRT2.* **(B)** *SVP1*. For both genes, the A genome mutants are indicated above the gene structure diagram and the B genome mutants below. **(C)** Reference map of the crosses used to generate *vrt2* and *svp1* loss-of-function mutants and the higher order mutants used in this study: **1.** Interaction between a transgene with constitutive *VRT2* expression (*UBI::VRT2*) with the *ful2* mutant. **2.** Complementation of *vrt2* with *UBI::VRT2*. **3.** Generation of a *vrt2 svp1* double mutant. **4.** Interactions between *vrt2* and *vrn1 ful2* mutants generated by Li et al. (2019). / *N indicates the number of crosses to Kronos recurrent parent performed to reduce background mutations.

The selected *vrt2* and *svp1* mutants are likely loss-of-function mutants (or severely hypomorphic mutants) because the encoded proteins have truncations that eliminate more than half of the conserved K domain or, for one of the alternative splice forms of *svp-A1*, a protein with a large deletion including parts of the MADS and K domains (Figure 2B and Supplemental Figure 2). Figure 2C presents the crosses and backcrosses used to generate the *vrt2* and *svp1* mutants, the double *vrt2 svp1* mutant and the higher order mutants described in other sections of this study.

### The *vrt2* and *svp1* mutations delay heading time, reduce plant height and increase number of spikelets per spike

Once we obtained single *vrt2* and *svp1* mutants, double *svp1 vrt2* mutants and the WT sister lines, we characterized them under controlled conditions using a 2 × 2 factorial design. The *vrt2 svp1* double mutants flowered later and were shorter than the single mutants or the wild type (WT, Figure 3A) and had spikes with a larger number of spikelets (Figure 3B). The factorial ANOVAs for heading time (Figure 3C), leaf number (Figure 3D), spikelet number per spike (SNS, Figure 3E) and plant height (Figure 3F) revealed highly significant effects (*P* < 0.001) for both *VRT2* and *SVP1* and highly significant interactions (*P* < 0.001, except for the plant height interactions with *P=* 0.0032, Supplemental Table 5). The *vrt2 svp1* double mutant showed stronger effects than the individual mutants for all traits (Figure 3C-F), which indicates overlapping roles of the two genes.

**Figure 3.**
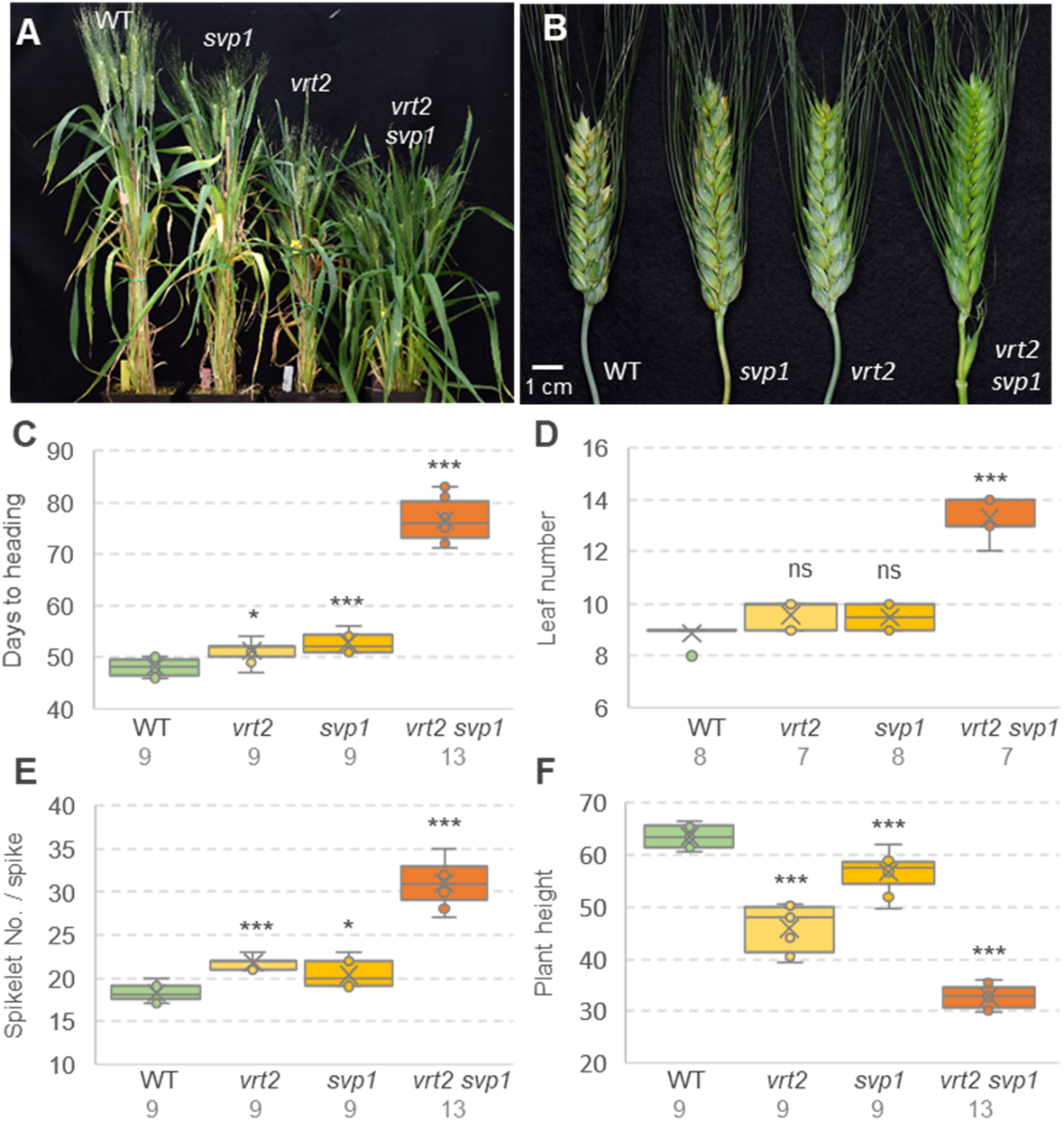
Effects of individual and combined *vrt2* and *svp1* mutants on important agronomic traits. **(A)** Plants 80 days after planting. **(B)** Spikes (note the axillary spikelet in the first node in *vrt2 svp1*). **(C)** Days to heading. **(D)** Leaf number. **(E)** Spikelet number per spike. **(F)** Plant height. **(C-F)** The number of plants analyzed is indicated below the genotypes. ns = not significant, * = *P* < 0.05, *** = *P* < 0.001 for differences with WT using Dunnett tests. Box-plot features are explained in the Statistical analyses section of Material and Methods.

For both *VRT2* and *SVP1*, the mutants combining homozygous mutations in the A and B genomes showed stronger effects than the homozygous mutants for a single homeolog (Supplemental Figure 3), which indicates redundant effects between homeologs for all the analyzed traits. Since the understanding of the effects of individual homeologs is critical for wheat breeding applications, we describe below their individual effects for each trait. The factorial ANOVAs for the homeologs of each gene are described in Supplemental Table 6.

#### Heading time

Both single *vrt2* and *svp1* mutants headed later than WT in two independent experiments (2.9 - 4.8 d, Figure 3A and C, Supplemental Figure 3A and B, and Supplemental Table 7), indicating a small but consistent effect of both genes on heading time. For this trait, the effects of the individual homeologs were not significant in the Dunnett tests (Supplemental Figure 3A and B). However, analysis of the four simple effects in factorial ANOVAs using homeologs as factors and alleles as levels revealed significant differences (*P < 0.05*) when each homeolog was tested in the presence of the mutant allele of the other homeolog, and the result was similar for *VRT2* and *SVP1*. These results indicate redundant roles of the individual homeologs on heading time.

#### Leaf number

The difference between the combined *vrt2 svp1* mutant and the WT was more than 6-fold larger than the individual mutants (4.4 leaves) and was highly significant (*P* < 0.001, Figure 3D), which indicates that part of the strong delay in heading time in *vrt2 svp1* was caused by a delay in the transition of the SAM between the vegetative and reproductive stages. For this trait, we did not study the effect of individual homeologs because the effects of the *vrt2* and *svp1* mutants for both homeologs were already very small (0.6-0.7 leaves, Supplemental Table 7).

#### Spikelet number per spike (SNS)

We observed larger and more significant differences in SNS between *vrt2* and WT (3.6 - 4.9 spikelets) than between *svp1* and WT (2.1 - 2.9 spikelets, Figure 3E, Supplemental Figures 3C and D, and Supplemental Table 7). The differences between the *vrt2 svp1* double mutant and WT (13.0 spikelets, ~70% increase) were larger than the added differences of the individual mutants, supporting the interactions between these two genes. A comparison of the main effects of the individual homeologs showed a stronger effect of *VRT-A2* than for *VRT-B2*, and the opposite for *SVP1* (*SVP-B1* stronger than *SVP-A1*, Supplemental Tables 6 and 7). Given the important role of SNS in grain yield, we also evaluated the four *VRT2* genotypes in the field. This experiment showed similar results to the growth chamber experiments with a stronger effect of *vrt2* relative to *vrt-A2* or *vrt-B2* (Supplemental Figure 3G).

#### Plant height and internode length

The same field experiment revealed large and significant differences in plant height (Supplemental Figure 3H and I, *P* < 0.0001) and a highly significant interaction between *VRT-A2* and *VRT-B2* (*P* < 0.0001, Supplemental Table 6). The single *vrt-A2* and *vrt-B2* mutants were approximately 2 cm shorter than the WT, whereas *vrt2* was 28 cm shorter (31% reduction, Supplemental Figure 3H and I), supporting the significant interaction.

We also performed a more detailed characterization of the effects of *vrt2* and *svp1* on the length of individual stem internodes in three growth chamber experiments. These studies revealed a stronger reduction of peduncle length in *vrt2* than in *svp1* (Supplemental Figure 3E and F), and very short peduncles in the *vrt2 svp1* double mutant (2.0 ± 0.2 cm, Supplemental Table 7). Comparisons between homeologs showed similar peduncle length reductions for *vrt-A2* (9.1 %) and *vrt-B2* (9.2 %), but a larger reduction for *svp-B1* (18.2 %) than for *svp-A1* (2.4 %, Supplemental Figure 3E and F, Supplemental Tables 6 and 7).

The length of the lower internodes was more variable than the peduncle length. For the first internode below the peduncle (henceforth internode −1), we observed a highly significant reduction in length in *vrt2 svp1*, and a small but significant increase in length in the *vrt2* and *svp1* single mutants (Supplemental Figure 3E and F, Supplemental Table 7). The increases in internode −1 were smaller than the decreases in peduncle length (Supplemental Table7), resulting in an overall reduction in plant height for the single mutants (Supplemental Figure 3E and F). The next two basal internodes (−2 and −3) showed smaller differences among genotypes, but were all reduced in *vrt2 svp1* (Supplemental Table 7).

In summary, these results indicate that *VRT2* and *SVP1* have overlapping functions during early reproductive development in wheat, both accelerating the transition from vegetative meristem to IM, and from IM to terminal spikelet, and both promoting elongation of the peduncle.

### The *vrt2 svp1* mutant has axillary spikelets or spikes at the nodes of the elongating stems

A surprising characteristic of the *vrt2 svp1* mutant was the presence of axillary spikelets and spikes in the nodes below the peduncle and the other elongating internodes (Figure 4A). Although axillary inflorescences are common in some species from other grass families including the Bambusoideae, Andropogoneae and Panicoideae (Stapleton, 1997; Vegetti, 1999), wheat does not have axillary buds in the nodes of the elongating stem (Figure 4 B-D) (Williams and Langer, 1975).

**Figure 4.**
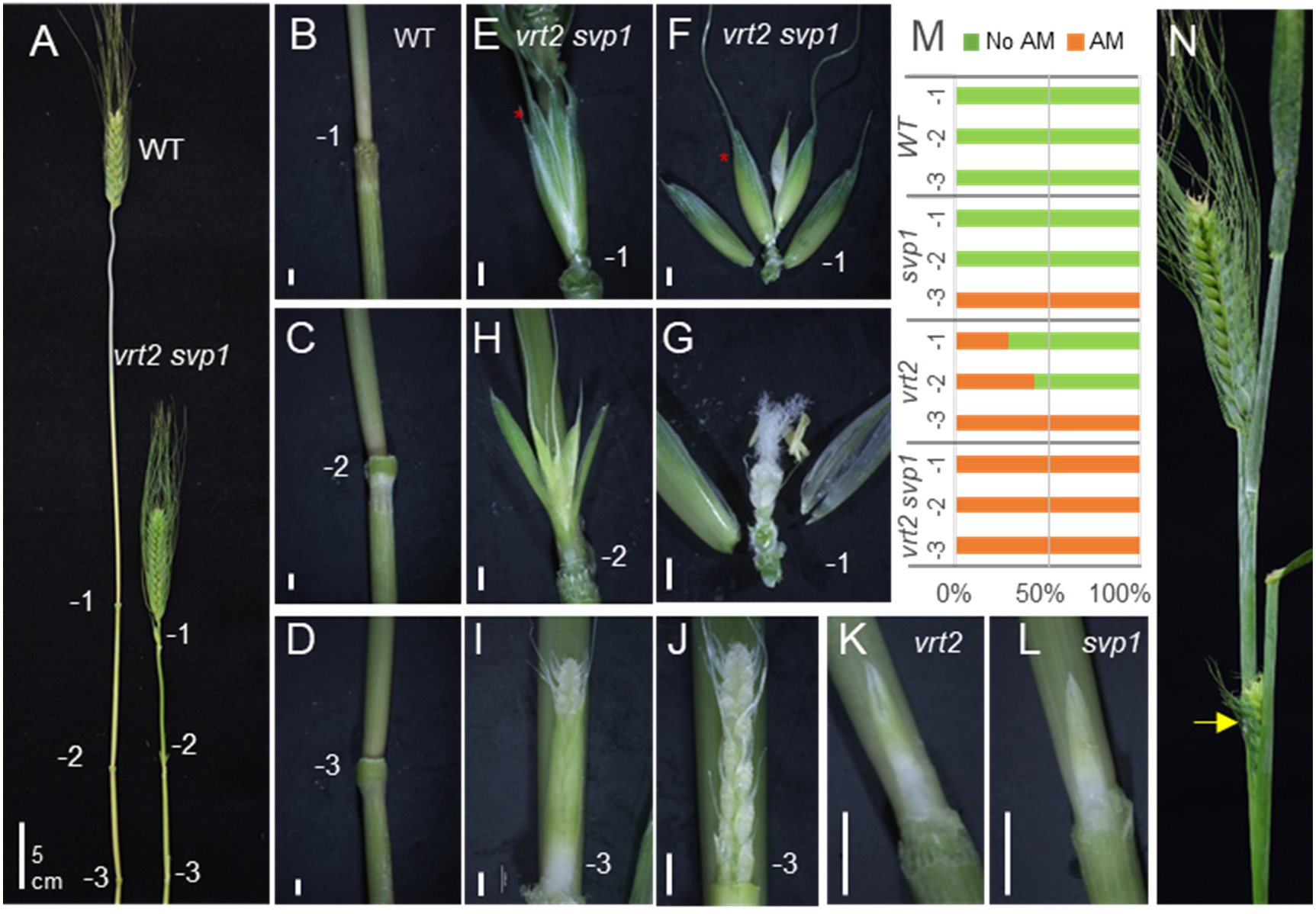
Axillary inflorescences in *vrt2, svp1*, and *vrt2 svp1*. **(A)** Comparison of internodes in WT and *vrt2 svp1* (−1 is the node below the peduncle, and −3 is the most basal node). **(B-D)** Detail of the three nodes in WT. **(E-J)** Nodes in *vrt2 svp1.* **(E)** Spikelet in internode −1. **(F)** Dissection of the spikelet showing glumes and three florets. **(G)** Dissection of floret one (red star in F) showing normal floral organs. **(H)** Spikelet in internode −2. **(I)** Axillary spike in internode −3 surrounded by a bract. **(J)** Same axillary spike without the bract showing lateral spikelets. **(K)** Axillary spike in *vrt2* surrounded by a bract. **(L)** Axillary spike in *svp1* surrounded by a bract. **(M)** Proportion of plants with axillary meristems in each of the three internodes below the spike (n = 7). Green = no axillary meristem (AM), Orange= Developed axillary meristem (spike or spikelet). **(N)** The yellow arrow points to an axillary spike emerging from its subtending leaf in *vrt2 svp1*. Bars in B to L are 2 mm (5 cm in A).

We observed a gradient in the development of the axillary buds, with those in node −1 developing into a single spikelet (Figure 4E), those in node −2 into single or sometimes two spikelets, and those in the more basal node −3 developing into more complex spikes (Figure 4I, J and N). The single axillary spikelets in node −1 showed normal glumes, lemmas, paleas, lodicules and floral organs (Figure 4F and G). The axillary spikes in the lower nodes were subtended and wrapped by a bract. As the axillary spike developed, it emerged from this bract, similarly to a regular spike emerging from the sheath of a flag leaf. However, this bract did not have a lamina (Figure 4I). When we removed this bract we observed a normal developing spike (Figure 4J), which was delayed in its development relative to the regular spikes at the same time point (Figure 4A). These unusual axillary spikelets and spikes were also observed in some nodes of *vrt2* (Figure 4K) and *svp1* (Figure 4L) individual mutants, but they were less frequent (Figure 4M) and usually less developed than in the *vrt2 svp1* double mutant. At later stages, some axillary spikes emerged from their subtending leaves in *vrt2 svp1* (node −3, Figure 4N arrow).

### Constitutive expression of *VRT2* alters spike development

To further characterize the *SVP*-like genes, we generated transgenic plants constitutively expressing *VRT-A*^*m*^*2* (cloned from *T. monococcum* A^m^ genome) under the maize *UBIQUITIN* promoter (hereafter referred to as *UBI::VRT2*). We characterized three independent transgenic events (T#2, T#4 and T#8), which displayed different levels of spike defects (Figure 5), which partially correlated with *VRT2* transcript levels in the developing spike at the TS stage. Transgenic plants T#2 and T#4 showed higher transcript levels than T#8, and all three had transcript levels significantly higher (*P* < 0.0001) than the non-transgenic sister lines (WT, Figure 6A).

**Figure 5.**
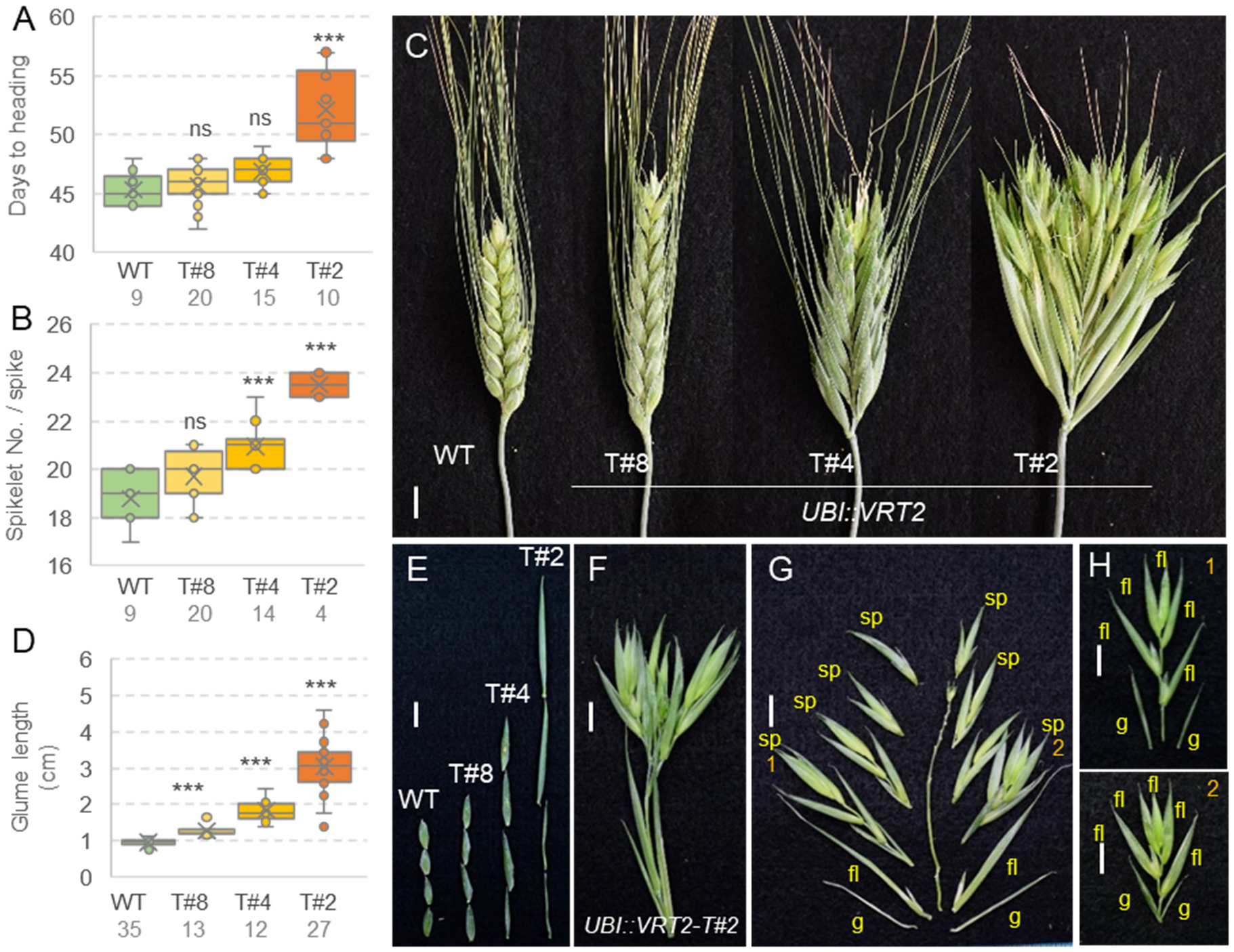
Phenotypic characterization of Kronos lines constitutively expressing *VRT2*. Three independent events with weak (T#8), intermediate (T#4) and strong phenotypes (T#2). **(A)** Days to heading. (B) Spikelet No. per spike. **(C)** Spike phenotype. **(D)** Glume length. **(A, B, D)** ns = not significant, ** = *P* < 0.01, *** = *P* < 0.001 in Dunnett tests versus WT control. Box-plot features are explained in the Statistical analyses section of Material and Methods. **(E)** Aligned glumes showing difference in length. **(F)** Basal “spikelet” from *UBI::VRT2* event T#2. **(G)** Dissection of the basal “spikelet” shows a determinate branch with multiple spikelets. **(H** and **I)** Detail of spikelets 1 and 2 in panel (G), each with glumes, florets and an elongated rachilla. Bars= 1cm. g = glume, fl= floret and sp= spikelet.

**Figure 6.**
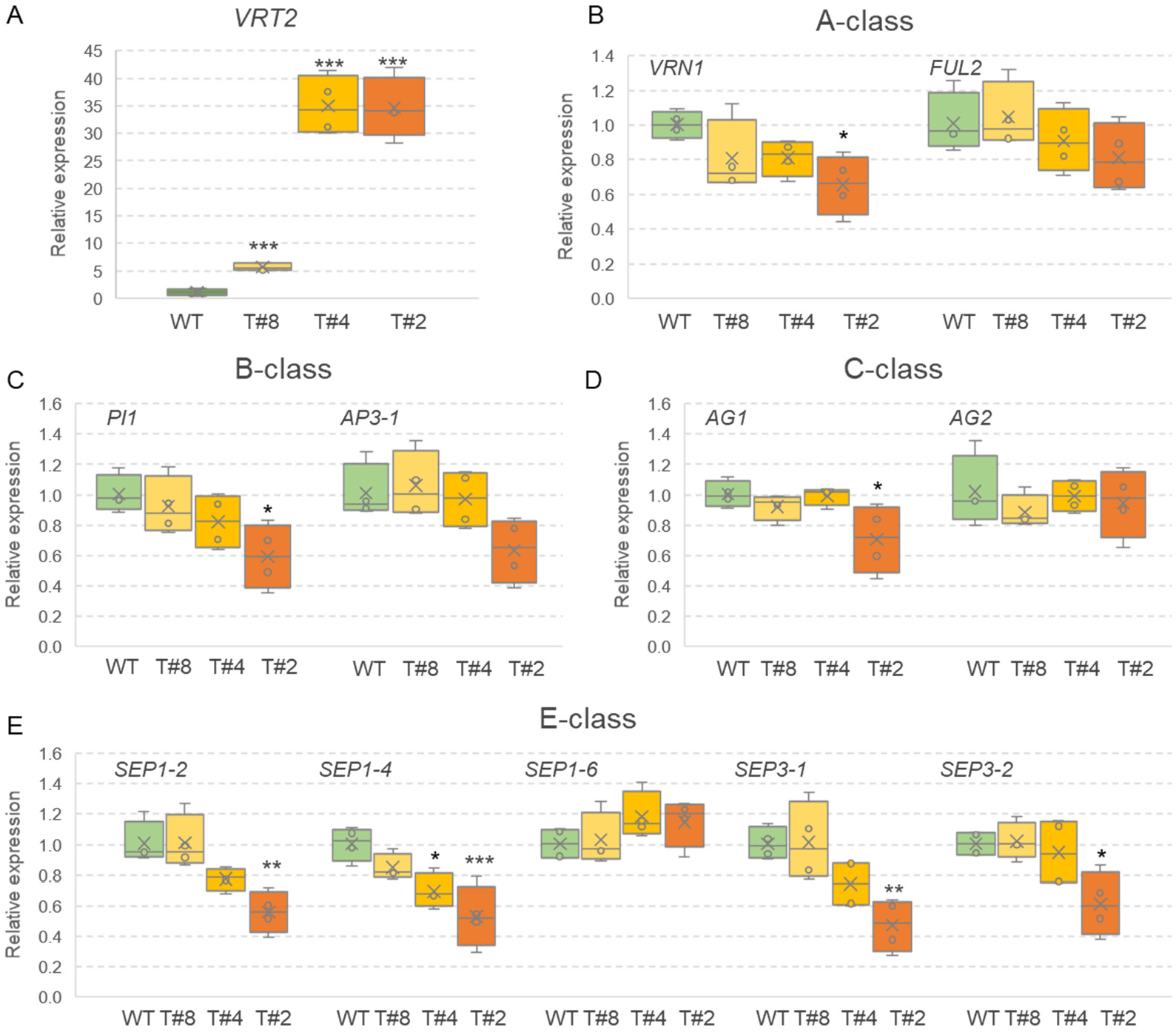
Relative expression of wheat flowering genes in developing spikes at the TS stage of *UBI::VRT2* transgenic lines T#8, T#4 and T#2 and sister lines without the transgene (WT). **(A)** *VRT2* (transgenic plus endogenous transcripts). **(B)** A-class MADS-box genes *VRN1* and *FUL2*. **(C)** B-class MADS-box genes *PI1* (~ *OsMADS4*) and *AP3* (~ *OsMADS16*). **(C)** C-class MADS-box genes *AG1* (~ *OsMADS58*) and *AG2* (~ *OsMADS3*). **(D)** E-class MADS-box genes *SEP1-2* (~ *OsMADS1*), *SEP1-4* (~ *OsMADS5*), *SEP1-6* (~ *OsMADS34*), *SEP3-1* (~ *OsMADS7*) and *SEP3-2* (~ *OsMADS8*). Graphs are based on 4 biological replicates (each replicate is a pool of 6-8 developing spikes at the TS stage). * = *P* < 0.05, ** = *P* < 0.01, *** *= P* < 0.001 in Dunnett tests versus the WT control. Expression was determined by qRT-PCR using *ACTIN* as endogenous controls and normalization relative to the WT (WT= 1). Box-plot features are explained in the Statistical analyses section of Material and Methods.

Transgenic T#2 headed 6.8 d later than WT and T#4 1.5 d later than WT, but only the T#2 difference was statistically significant (Figure 5A, *P* < 0.0001). Both lines also showed significant differences relative to the WT in SNS (*P* < 0.0001), with T#2 showing a larger increase (4.7 spikelets) than T#4 (2.2 spikelets) (Figure 5B). All transgenic plants showed changes in spike or spikelet morphology (Figure 5C), ranging from a slight increase in glume length in T#8 (0.34 cm, *P* < 0.01, Figure 5D and E), followed by longer glumes in T#4 (1.2 cm), and ending with T#2, which exhibited severe morphological alterations. These included very long glumes (2.6 cm, Figure 5D and E) and lemmas, and replacement of basal spikelets by branches with multiple spikelets (which also have elongated glumes, lemmas and rachilla, Figure 5F-H).

We also explored the ability of the weakest *UBI::VRT2* transgenic line (T#8) to complement the changes observed in *vrt2* in the F_2_ progeny of a cross between T#8 and *vrt2* (Figure 2C). We selected lines homozygous for *vrt2* or WT, and within each of them lines with the transgene (T) or without it (NT). Within the non-transgenic plants, *vrt2* mutants headed 3.2 d later than WT, but those differences disappeared in the presence of the transgene, indicating full complementation (Supplemental Figure 4A). It is interesting to point out that the weak transgenic line T#8 accelerated heading time both in the *vrt2* and *Vrt2* (WT) backgrounds, whereas the strong transgenic line T#2 headed one week later than the WT (Figure 4A). These results indicate that strong ectopic expression of *VRT2* can revert its normal role as a promoter of heading time.

The differences in peduncle length between the WT and *vrt2* mutant in the non-transgenic line (19.6 cm) were significantly reduced in the *UBI::VRT2* transgenic line (11.4 cm, Supplemental Figure 4B), indicating partial complementation. However, there was no complementation for the differences in SNS, with similar increases in SNS in the *vrt2* mutant relative to the WT in the transgenic and non-transgenic backgrounds (Supplemental Figure 4C). This result is not surprising since the weak T#8 transgene resulted in non-significant effects on SNS (Figure 5B).

To understand better the effect of *VRT2* on the regulation of spikelet development, we used qRT-PCR to compare the transcript levels of several MADS-box flowering regulators between the three *UBI::VRT2* transgenic and the non-transgenic sister line at the TS stage (Figure 6). The T#2 transgenic line, which displayed the strongest phenotypic effects and high transcript levels of *VRT2*, showed significant reductions in the transcript levels of wheat A-class gene *VRN1*, B-class gene *PI1*, C-class gene *AG1*, and E-class genes *SEP1-2*, *SEP1-4*, *SEP3-1* and *SEP3-2* (Figure 6B-E).

The T#4 transgenic line (intermediate spike phenotypes) showed reductions in the transcript levels of the same genes down-regulated in T#2, but the differences were significant only for *SEP1-4* (Figure 6E). The T#8 transgenic line, with the lowest increase in *VRT2* transcripts and the mildest phenotypic effects, showed no significant differences in transcript levels for any of the floral genes, but a small reductions in *VRN1*, *PI1*, and *SEP1-4* paralleled the significant reductions in the stronger transgenic plants (Figure 6A, C and E). In summary, the magnitude of the spike phenotypic changes in the different transgenics correlated well with the levels of downregulation of the transcripts of multiple flowering regulatory genes.

### The *ful2* mutant enhances spike defects in weak *UBI::VRT2* transgenic plants

Strong constitutive expression of *VRT2* results in spikelets with leafy-like glumes and lemmas similar to those observed in partial *vrn1 ful2* mutants carrying one functional copy of *VRN-A1* in heterozygous state and no functional *ful2*, which was previously designated as *<u>Vrn1</u> ful2* (Li et al., 2019). Based on these results, we hypothesized that *VRT2* and *FUL2* may have opposite effects on spikelet development. To test if the combination of *ful2* and *UBI::VRT2* would enhance the defects of the individual lines, we crossed the weak *UBI::VRT2* T#8 with *ful2*.

For stem length, *ful2* showed a significant reduction in length (*P* < 0.001) but there was no significant effect for *UBI::VRT2* or for the *FUL2* × *UBI::VRT2* interaction in the factorial ANOVA (Figure 7A and Supplemental Table 8). By contrast, the ANOVAs for SNS, glume length and lemma length showed significant effects for both *FUL2* and *UBI::VRT2* (*P* < 0.001) and a highly significant interaction between them (*P ≤* 0.001, Supplemental Table 8). In all three traits the *UBI::VRT2* – *ful2* showed values that were higher than the addition of the individual effects (Figure 7B-D). Plants combining the *UBI::VRT2* T#8 and *ful2* showed long leafy-like glumes and lemmas in young spikes, similar to those of the T#4 and T#2 transgenic lines with stronger phenotypes (Figure 7E). In more mature spikes, most of the spikelets of the *UBI::VRT2* – *ful2* plants showed a large increase in floret number (Figure 7F), a phenotype reported previously in partial *<u>Vrn1</u> ful2* mutant plants (Li et al., 2019). Dissection of the basal spikelets showed that some florets were replaced by spikelets and that the rachilla ended in a terminal spikelet, resembling a determinate branch (Figure 7G-I, 7 out of 8 plants analyzed showed branching). Taken together these results suggest that *VRT2* and *FUL2* have antagonistic effects on spikelet development.

**Figure 7.**
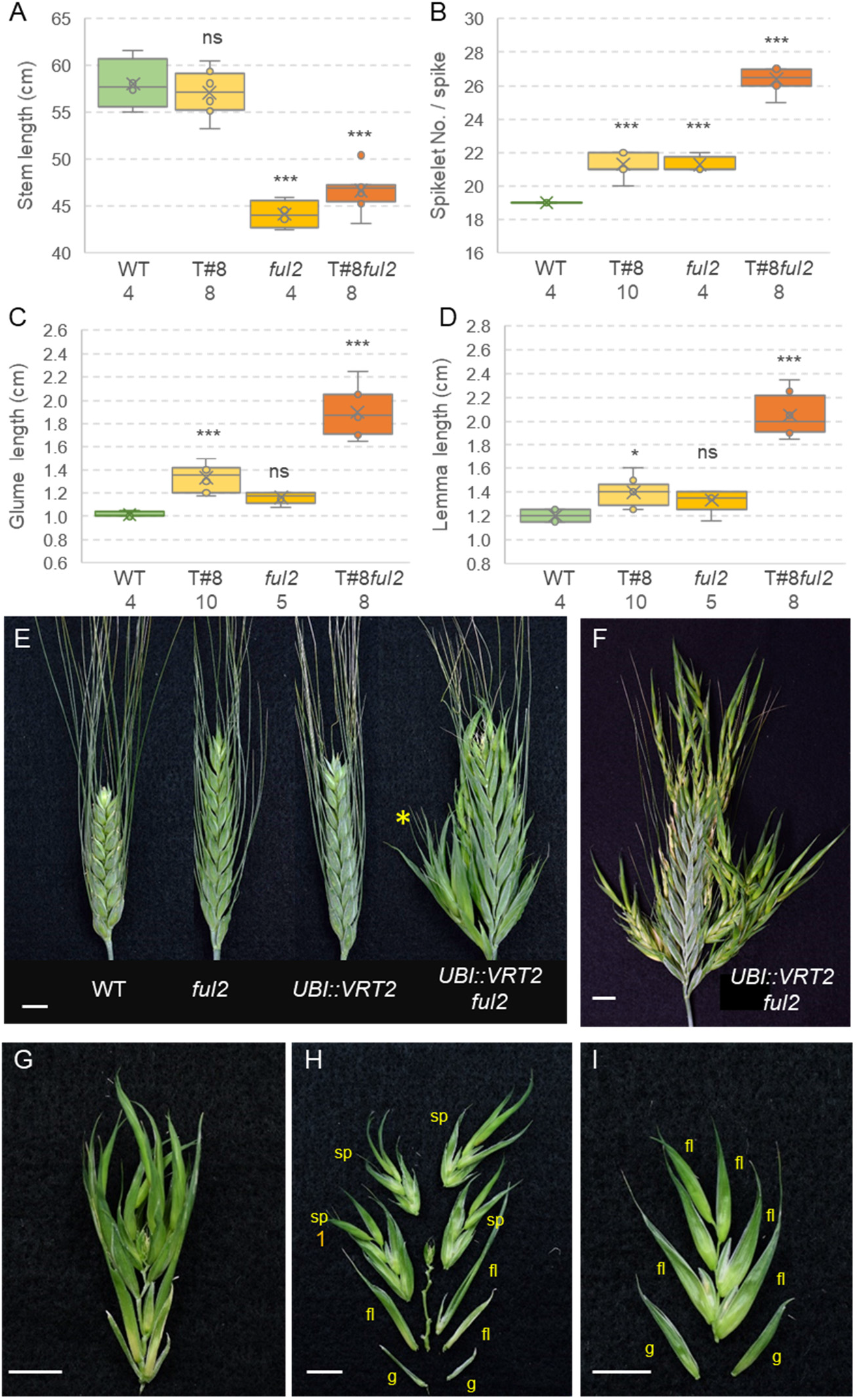
Effect of combined *ful2* mutation and *UBI::VRT2* T#8 transgene on stem length and spike and spikelet development. **(A)** Stem length (without spikes). **(B)** Spikelet number per spike (SNS). **(C)** Glume length in cm. **(D)** Lemma length in cm. WT= homozygous *Ful2* and not transgenic. T#8: weak constitutive transgenic *UBI::VRT2* line T#8. *ful2* = loss-of-function mutant for *ful-A2* and *ful-B2* and not transgenic. T#8*ful2 =* homozygous *ful2* and T#8 transgenic present. N= number of plants analyzed is indicated below genotype. ns = not significant, * = *P* < 0.05, ** = *P* < 0.01, *** = *P* < 0.001 in Dunnett tests. Box-plot features are explained in the Statistical analyses section of Material and Methods. **(E)** Young spikes of WT Kronos, *ful2*, *UBI::VRT2* T#8, and combined *UBI::VRT2* T#8 –*ful2*. **(F)** A more mature spike of *UBI::VRT2* T#8 – *ful2* **(G)** Basal spikelet transformed in a branch (yellow star in A) **(H)** Dissection of the basal spikelet converted in branch showing lateral spikelets **(I)** Dissection of the spikelet marked with number 1 in H. Bars = 1 cm. g = glume, fl= floret and sp= spikelet.

### The *vrt2* mutant reduces spikelet developmental defects in the partial *<u>Vrn1</u> ful2* mutant

Since *VRT2* constitutive expression exacerbated the spikelet defects observed in the *ful2* mutant, we hypothesized that a *vrt2* mutation could reduce some of the vegetative characteristics of the *vrn1 ful2* spikelets. To test this hypothesis, we intercrossed *vrt2* and *<u>Vrn1</u> ful2* and selected a triple homozygous *vrt2 vrn1 ful2* mutant and a partial mutant with one functional copy of *Vrn-A1* (*vrt2 <u>Vrn1</u> ful2*). All mutants were developed in a *vrn2-*null background to avoid late flowering.

The *vrn1 ful2* mutant produced spikes in most shoots, in which the lateral spikelets were replaced by vegetative tillers (henceforth, spike-of-tillers), a phenotype already described in a previous study (Li et al., 2019). Compared to *vrn1 ful2*, the triple *vrt2 vrn1 ful2* mutants were shorter and showed emerging spikes-of-tillers in only 16 % of shoots (Supplemental Figure 5A, B and E).

Dissections of some of the shoots without emerging spikes revealed underdeveloped spikes with very short peduncles and internodes (Supplemental Figure 5F), most of which eventually died within the sheaths (Supplemental Figure 5G). The few emerging spikes, showed lateral spikelets replaced by vegetative tillers similar to the double *vrn1 ful2* mutant (Supplemental Figure 5C and D). Taken together, these results suggest that *VRT2* and *VRN1/FUL2* interact to regulate internode elongation and inflorescence development, and that *VRN1* and *FUL2* may be essential to initiate spikelet development (*vrt2* alone is not sufficient to revert this phenotype).

Since the *vrn1 ful2* and *vrt2 vrn1 ful2* have no spikelets, we had to use a genotype with at least one functional *SQUAMOSA* genes to study the interactions of these genes on spikelet development. For these experiments, we compared the double *<u>Vrn1</u> ful2* partial mutant (one functional copy of *Vrn-A1*) with the triple *vrt2 Vrn1 ful2* mutant (Figure 8 and Supplemental Figure 6). The *vrt2 <u>Vrn1</u> ful2* mutant headed 3.5 d later (Supplemental Figure 6A), had a 2.2 cm shorter stem (Supplemental Figure 6B) and produced 4.7 more spikelets per spike (Supplemental Figure 6C) than the *<u>Vrn1</u> ful2* mutant. Interestingly, we observed that while *<u>Vrn1</u> ful2* spikes had long and leafy glumes and lemmas (Figure 8A and B), *vrt2 <u>Vrn1</u> ful2* had significantly shorter glumes and lemmas (27-28% shorter, Figure 8C and D, Supplemental Figure 6D and E). In older spikes, spikelets from *<u>Vrn1</u> ful2* showed a very large number of florets (Figure 8E), but that number was greatly reduced in *vrt2 <u>Vrn1</u> ful2*, resulting in a more normal spike (Figure 8F).

**Figure 8.**
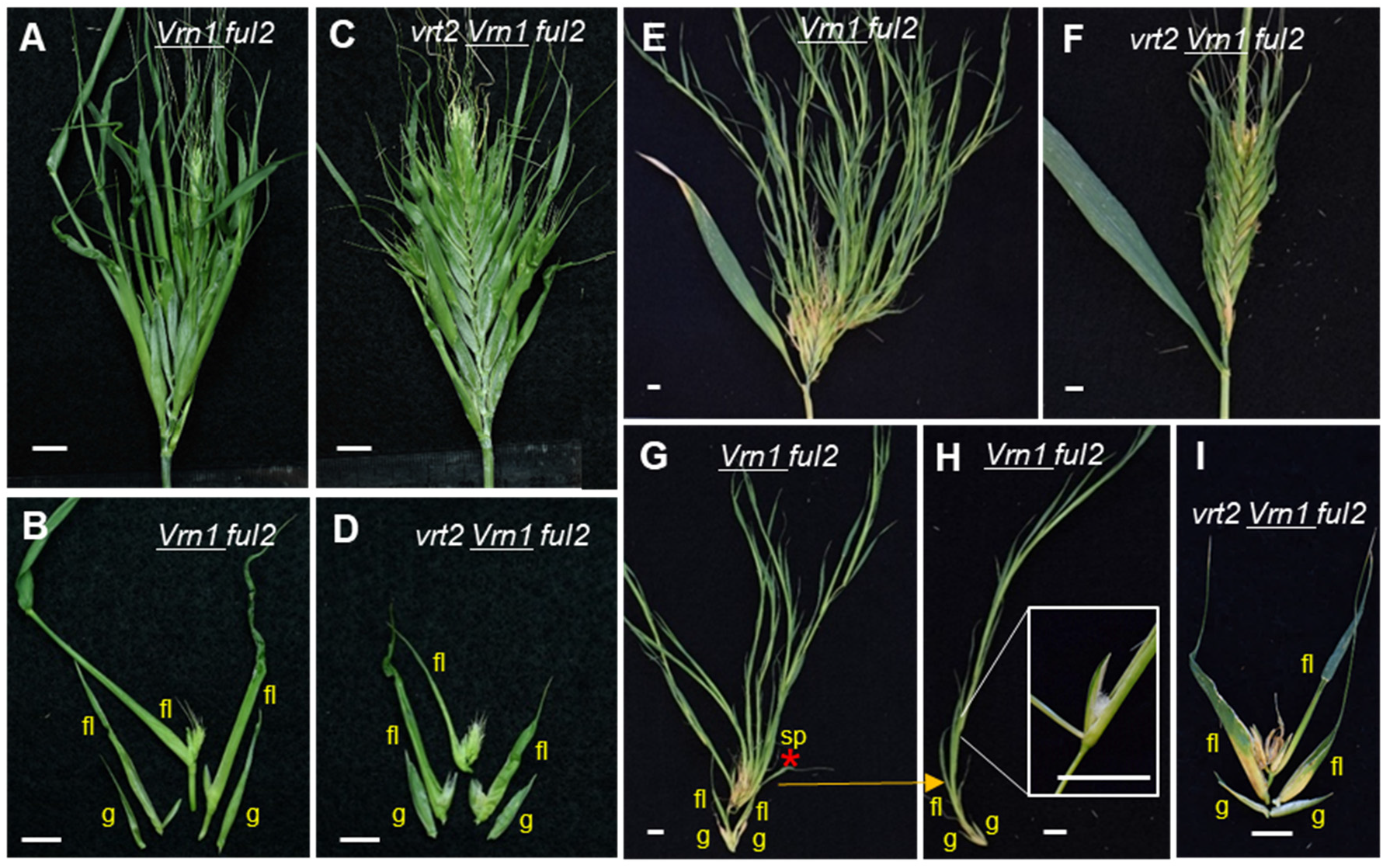
Effect of combined *<u>Vrn1</u> ful2* and *vrt2* mutations on spike and spikelet development. **(A, B, E, G-H)** *<u>Vrn1</u> ful2.* **(C, D, F, I)** *vrt2 <u>Vrn1</u> ful2.* **(A** and **C)** Young spikes. **(B** and **D)** Dissection of basal spikelets. **(E-F)** Older spikes. **(G-I)** Dissection of older spikelets. **(H)** Detail of the third “floret” in G (red asterisk) that reverted to a spikelet with its own glumes. The inset in **(H)** shows a floret of this spikelet. **(I)** Spikelet of the same age as in **G** from *vrt2 <u>Vrn1</u> ful2.* g = glume, fl = floret and sp= spikelet. Bars = 1 cm.

Dissection of the spikelets from both genotypes showed that 80.4% of the *<u>Vrn1</u> ful2* spikelets (n = 56) had ‘branches’ as a result of the regression of some of their florets into spikelets (Figure 8G and H, Supplemental Figure 6F). By contrast, only 30.1% of the *vrt2 <u>Vrn1</u> ful2* spikelets (n = 51) showed regression of florets to spikelets, with most of the spikelets showing a normal number of florets and an appearance more similar to a normal spikelet, although still with some leafy characteristics in the lemmas (Figure 8I). These results indicate that *VRT2* is partially responsible of the spikelet defects of the *<u>Vrn1</u> ful2* mutants.

### Wheat SQUAMOSA proteins interact with SVP and SEPALLATA MADS-box proteins

Characterization of protein interactions using yeast two-hybrid (Y2H) assays is a first step to identify proteins that possess the capacity and specificity to interact with each other. MADS-box proteins are known to interact with each other and form complexes with critical functions in floral development, so we decided to characterize the pairwise interactions among wheat proteins from the SQUAMOSA, SVP and SEPALLATA clades. We first confirmed that none of these wheat proteins caused autoactivation in Y2H assays (Supplemental Figure 7).

We then tested interactions within and between the proteins of the SQUAMOSA and SVP clades. Individual members of the SVP-clade did not interact with each other, and only SVP1 was able to form homodimers (Figure 9 and Supplemental Figure 8). Among the SQUAMOSA proteins, FUL2 and FUL3 showed strong and weak homodimerization, respectively (Supplemental Figure 8). FUL2 interacted with both VRN1 and FUL3, whereas the latter two did not interact with each other (Supplemental Figure 8). Pairwise interactions between proteins from the two clades revealed that VRT2 and SVP1 can interact with all three wheat SQUAMOSA proteins, whereas SVP3 can interact only with FUL2. The interactions of SVP1 with all three SQUAMOSA proteins were of similar strength but the VRT2 interaction were weakest with VRN1 and strongest with FUL2 (Supplemental Figure 9).

**Figure 9.**
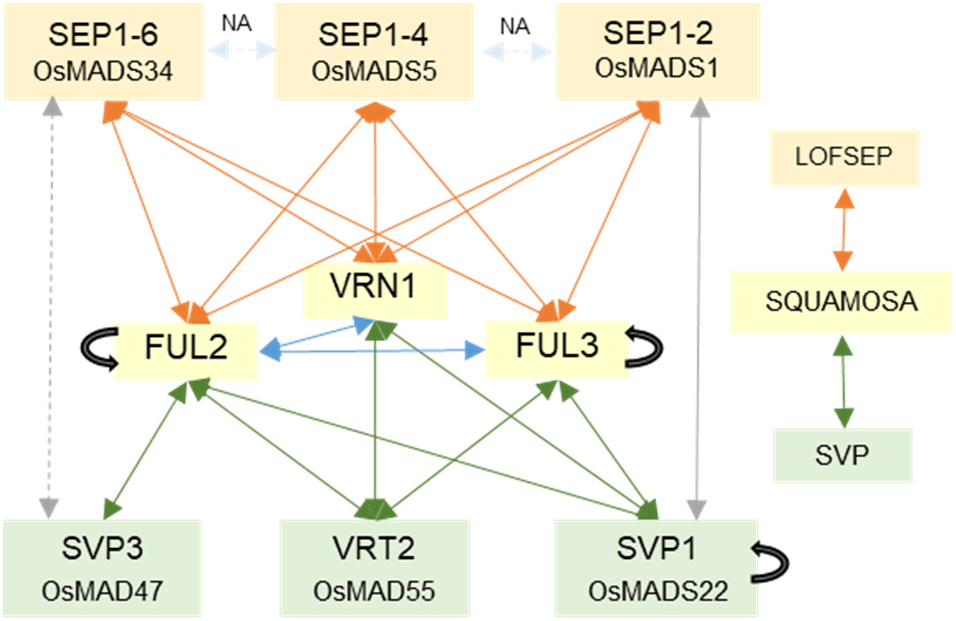
Yeast-two-hybrid (Y2H) interactions between wheat SQUAMOSA, SVP and SEPALLATA proteins. Wheat MADS-box proteins of the SQUAMOSA-clade are indicated by yellow boxes (VRN1, FUL2 and FUL3), proteins of the SVP-clade by green boxes (VRT2, SVP1 and SVP3) and proteins of the LOFSEP-clade by orange boxes (SEP1-2, SEP1-4, and SEP1-6). Positive interactions between SQUAMOSA and LOFSEP-clade proteins are shown with orange arrows, between SQUAMOSA- and SVP-clade proteins with green arrows, and between SVP- and LOFSEP-clade proteins in grey (the weak interaction between SVP3 and SEP1-6 is indicated by a dotted line). Black curved arrows indicate positive homodimerization. Interactions among SEPALLATA proteins were not analyzed.

In rice, SQUAMOSA proteins also interact with MADS-box proteins of the SEPALLATA subclade LOFSEP (Wu et al., 2018), which includes OsMADS1 (ortholog of wheat SEP1-2), OsMADS5 (ortholog of wheat SEP1-4), and OsMADS34 (ortholog of wheat SEP1-6). *LOFSEP* genes are essential for spikelet and floret organ identity in rice, with *osmads1 osmads5 osmads34* triple mutants producing spikelets with homeotic transformations of the sterile lemma and lemma/palea into leafy structures (Wu et al., 2018). Since this phenotype is similar to the one we observed in *UBI::VRT2* lines and *vrn1 ful2* mutants, we characterized the Y2H interactions between the proteins from the wheat LOFSEP-, SQUAMOSA- and SVP-clades.

We observed positive Y2H physical interactions for all nine possible pairwise combinations of the three wheat LOFSEP proteins with the three SQUAMOSA proteins (Figure 9). The interactions for all three LOFSEP proteins were strongest with FUL2, intermediate with FUL3 and weakest with VRN1 (Supplemental Figure 9). By contrast, there were fewer positive Y2H interactions between proteins of the LOFSEP and SVP clades. Among the nine possible pairwise combinations, we only detected a strong interaction between SEP1-2 and SVP1 and a weak interaction between SEP1-6 and SVP3. In summary, wheat proteins from the SQUAMOSA-clade interact in yeast with most proteins from the SVP and LOFSEP clades, whereas the latter two show limited interaction with each other (Figure 9).

We also used BiFC to validate the positive Y2H interactions in wheat protoplasts (Supplemental Table 9). We observed fluorescence signals in the nucleus and sometimes in the cytoplasm for nine of the 15 tested interactions (Supplemental Table 9 and Supplemental Figure 10A-I), whereas the six interactions of SEP1-4 and SEP1-6 with the SQUAMOSA proteins showed no nuclear fluorescent signals (Supplemental Figure 10J-O). Some of the positive and negative interactions showed fluorescent protein aggregates outside the nucleus (Supplemental Figure 10G and H, see food note). We did not detect fluorescent nuclear signals or aggregates for the negative controls using the YFP-C (C-terminal part of YFP) paired with individual proteins of all three clades fused to YFP-N (N-terminal part of YFP) (Supplemental Figure 10P-W). The negative interactions between the three SQUAMOSA proteins with SEP1-4 and SEP1-6 (Supplemental Table S6) serve as additional negative controls using proteins of the same families.

### Wheat SVP and LOFSEP proteins compete for interactions with SQUAMOSA in yeast

Since both SVP and LOFSEP proteins interact with SQUAMOSA proteins, we then tested if the presence of the SVP proteins could interfere with the interaction between the SQUAMOSA and LOFSEP proteins using Y3H assays. In these experiments, LOSEP proteins were fused to the GAL4 DNA binding domain, VRT2 was controlled by methionine levels in the medium (expressed in the absence of methionine, and repressed by inclusion of methionine), and SQUAMOSA proteins were fused to the GAL4 activation domain. Protein interaction strength was determined by quantitatively measuring α-galactosidase activity (henceforth α-gal assays) in the presence or absence of methionine.

The α-gal assays confirmed that all three LOFSEP proteins had much stronger interactions with FUL2 than with FUL3 or VRN1 (Figure 10). SEP1-4 and SEP1-6 showed interactions of similar strength with FUL3 and VRN1, but SEP1-2 interaction with FUL3 was stronger than with VRN1 (Figure 10). The expression of VRT2 as the competing protein in Y3H assays significantly reduced the α-gal activity of the three strong FUL2 - LOFSEP interactions (12.0% in SEP1-2, 20.6 % in SEP1-4 and 22.0 % in SEP1-6, *P* < 0.01). Among the weak interactions, the presence of VRT2 only had a significant effect on the FUL3 - SEP1-2 interaction (93.8 % reduction, *P* < 0.0001, Figure 10). Taken together, these results indicate that the presence of wheat VRT2 can interfere with some of the wheat SQUAMOSA - LOFSEP interactions in yeast.

**Figure 10.**
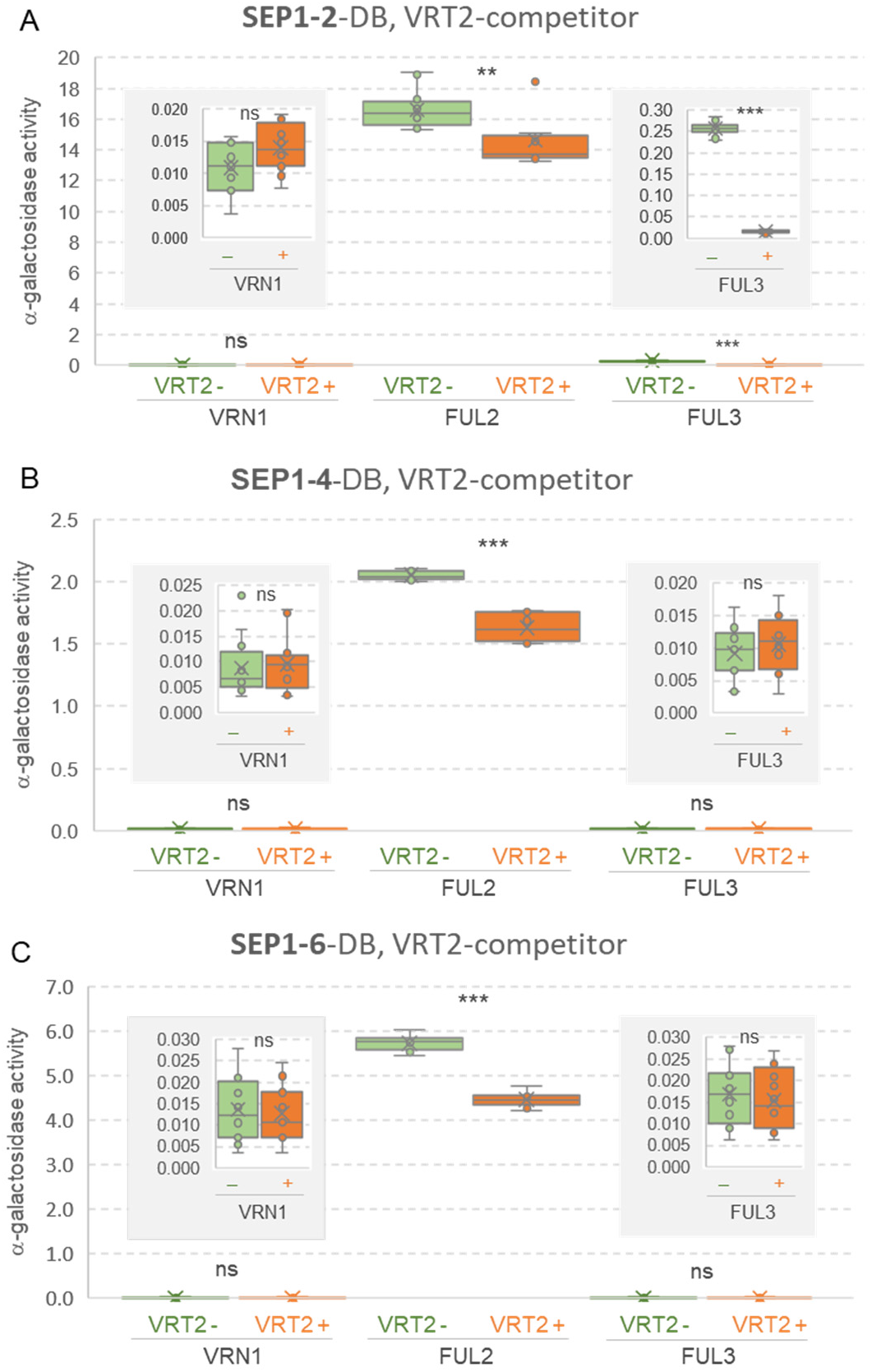
Wheat VRT2 competes with LOFSEP proteins for interactions with SQUAMOSA proteins. Yeast three-hybrid assays were used to test the effect of VRT2 as a competitor, when **(A)** SEP1-2 (~OsMADS1), **(B)** SEP1-4 (~OsMADS5), and **(C)** SEP1-6 (OsMADS34) were expressed as the DNA-binding domain fusions, and SQUAMOSA proteins VRN1/FUL2/FUL3 were expressed as activation domain fusions. The α-gal activity of the protein interactions in the absence of the competitor is shown in green box-plots and in the presence of the competitor in orange box-plots. Relative α-gal activity values for each interaction are the average of 12 replicates. ns= not significant, ** = *P* < 0.01 and *** *P* < 0.001). The insets show the α-gal activity for weaker interactions using different scales. Box-plot features are explained in the Statistical analyses section of Material and Methods.

## DISCUSSION

### Roles of *SQUAMOSA* genes in the transcriptome of developing spikes and spikelets in wheat

By comparing the transcriptomes of developing spikes from *vrn1 ful2* (where spikelets are replaced by tillers) and *vrn1* (normal spikelets) at four developmental stage, we identified downstream genes and pathways controlling spike and spikelet development (Figure 1A). An important observation from this analysis was the dramatic increase in the number of DEG after the double-ridge state (PDR). This observation suggests that this developmental stage is critical for the establishment of the different developmental fates of the lateral spike meristems in the *vrn1* and *vrn1 ful2* mutants. The lower number of DEGs in the vegetative and DR stages of spike development correlates with the similar morphology of the early developing spikes of *vrn1* and *vrn1 ful2* mutants up to DR stage. This last result indicates that both mutants have the ability to transition from the vegetative SAM to the IM (Li et al., 2019). The dramatic morphological differences in the spike lateral meristems appear at PDR and TS, which correlates with the increase in DEGs (Figure 1).

The large morphological differences at the later stages of spike development are reflected in the DEGs in clusters 4, 5, 9 and 10. Genes in cluster 4 are up-regulated at PDR in *vrn1* but not in *vrn1 ful2* (Figure 1D) and are enriched in genes involved in the regulation of the early stages of spike development. Genes in this cluster include several known regulators of axillary branches and spikelet meristem development and determinacy, which are described in more detail in Supplemental Table 3. Cluster 5 DEGs include multiple class-B, class-C and class-E (except *SEP1-6*) MADS-box floral genes that are up-regulated in *vrn1* at TS, reflecting the progress of the lateral meristems into spikelets and florets (Figure 1F). By contrast, transcript levels of the genes promoting floral development remain at low levels in clusters 9 and 10, whereas florigen antagonists *CEN2*, *CEN4* and *CEN5* are upregulated (Figure 1G). DEGs in these two clusters are also enriched in genes with photosynthetic functions reflecting the vegetative fate of the *vrn1 ful2* lateral meristems (Figure 1E).

Genes in cluster 8 peak at the DR stage and include several genes previously shown to control the number of spikelets per inflorescence (Supplemental Table 3). This cluster also includes *SEP1-6*, which is the earliest expressed of the *LOFSEP* genes during reproductive development in both wheat (Debernardi et al., 2020) and rice (Wu et al., 2018), and is the only wheat gene from the *LOFSEP*-clade up-regulated in the double *vrn1 ful2* mutant (Figure 1G).

Finally, cluster 7 includes genes down-regulated in PDR and TS in *vrn1* but not in *vrn1 ful2*, which is the expected profile for genes with potential roles in repressing SM identity. This cluster includes the three related MADS-box genes from the SVP clade (*VRT2, SVP1* and *SVP3*) (Figure 1G) that constitute the focus of this study, and genes known to extend the activity of the IM (resulting in increased spikelet number) or to delay the transition of meristems to SM identity (Supplemental Table 3).

In summary, the complete table of DEGs presented in Supplemental File 1 represents a valuable genomics resource for researchers interested in genes and gene networks that act downstream of *VRN1* and *FUL2* and that play important roles in the early stages of spike and spikelet development. Supplemental Table 3 highlights a subset of these genes, which have already been found to play important roles in inflorescence and flower development in grasses.

### Wheat genes from the *SVP* clade promote developmental transitions of vegetative and inflorescence apical meristems

#### Vegetative SAM transition to IM

In Arabidopsis, *SVP* acts as a flowering repressor and the related *AGL24* acts as a flowering promoter, showing the flexibility of MADS-box genes from the SVP-clade to work as flowering promoters or repressors. Our results show that changes in *VRT2* expression levels or in its spatio-temporal expression profiles can revert the function of this gene from a flowering promoter to a flowering repressor.

Both individual *vrt2* and *svp1* mutants headed approximately three days later than the WT, and the delay in *vrt2* was fully complemented by a weak constitutively expressed *VRT2* (*UBI::VRT2* T#8). The *vrt2 svp1* double mutant showed a more significant delay in heading time (28 d) and had 4.4 more leaves than the WT, which indicates that a delay in the transition of the SAM from the vegetative to the reproductive stage contributed to the delay in heading time. The wheat mutant results suggest that *VRT2* and *SVP1* normally function redundantly to accelerate the transition of the SAM from the vegetative to the reproductive stage.

Surprisingly, when *VRT2* spatio-temporal expression pattern and transcript levels were altered by a strong constitutive promoter, it delayed heading time almost one week. A similar delay (10 d) was observed in barley for *BM1* (~*SVP3*) but not for *BM10* (~*SVP1*) when both genes were constitutively expressed under the *ZmUBIQUITIN* promoter (Trevaskis et al., 2007). The effect of the constitutive expression of *VRT2* on heading time is likely modulated by genetic background and environment, since constitutive expression of *UBI::VRT2* in winter wheat accelerated flowering 5-6 d in unvernalized plants but not in fully vernalized plants (Xie et al., 2019).

A possible interpretation of the contrasting *VRT2* roles in mutants and transgenic plants is that changes in VRT2 protein abundance and distribution in the *UBI::VRT2* transgenic plants may affect the composition and stability of different MADS-box protein complexes resulting in multiple pleiotropic effects. The altered balance of these multiple effects across a complex and heavily interconnected regulatory network can determine different outcomes depending on the time, place and level of *VRT2* expression. This hypothesis is based on VRT2’s interactions with all SQUAMOSA proteins (Figure 9), its ability to compete with other MADS-box for interactions with SQUAMOSA proteins (Figure 10) and its transcriptional effects on multiple floral regulatory genes (Figure 6). A similar hypothesis may explain the contrasting effects of *VRT2* on SNS in mutants and *UBI::VRT2* transgenic plants described below.

#### IM transition to terminal spikelet

Individual *vrt2* and *svp1* mutants showed 4-5 more spikelets per spike than the WT (Figure 3B and E) and that difference increased to 13 in *vrt2 svp1*. An increase in SNS was also observed in individual *vrn1* and *ful2* mutants relative to the WT (Li et al., 2019), and in *vrt2 <u>Vrn1</u> ful2* triple mutant relative to *<u>Vrn1</u> ful2* or *vrt2* separately (Supplemental Figure 6C and Figure 3E). Taken together, these results suggest an overlapping role of *SQUAMOSA* and *SVP* genes in promoting or accelerating the transition of the IM into a terminal spikelet. However, constitutive expression of *VRT2* resulted in a delay of this transition, which was reflected in significant increases in SNS (Figure 5B). These results suggest that, as in the regulation of heading time, ectopic expression or drastic increases in *VRT2* transcript levels can reverse its normal role as a promoter of the transition from IM to terminal spikelet into a repressor of this transition. Even in this reversed role, *UBI::VRT2* showed a highly significant interaction with *FUL2* in the determination of SNS (Figure 7B, Supplemental Table 8), supporting a strong connection between *SQUAMOSA* and *SVP* genes in the regulation of the transition from IM to terminal spikelet in wheat.

In Arabidopsis, a complex involving SQUAMOSA proteins AP1 or CAL and any of SVP, AGL24, SOC1 or SEP4 proteins has been shown to control inflorescence branching by repressing expression of *TFL1*, a mechanism that seems to be conserved in rice (Liu et al., 2013). Expression levels of *TFL1* orthologs in rice, barley and wheat are positively correlated with the number of spikelets per inflorescences in these three species (Nakagawa et al., 2002; Wang et al., 2017; Bi et al., 2019). Our Quant-Seq results show that the transcript levels of the wheat *TFL1* homologs *CEN2*, *CEN4* and *CEN5* are higher in *vrn1 ful2* than in *vrn1* at the PDR and TS stages (Figure 1G). This result suggest a potential contribution of the wheat *CEN* genes to the increased SNS in the *vrn1* and *ful2* mutants and to the indeterminate IM in the *vrn1 ful2* double mutant.

In addition to the effect on SNS, *SQUAMOSA* genes have been shown to be critical for normal spike development in wheat (Li et al., 2019). In this study, we show that most spikes of the triple *vrt2 vrn1 ful2* mutant are unable to emerge from the shoots and remain undeveloped and eventually die within the sheaths (Supplemental Figure 5F and G). Similarly, in the Arabidopsis triple mutant *ap1 svp agl24* the IM fails to produce floral meristems (Gregis et al., 2008). Taken together, these results suggest that genes from both the *SQUAMOSA* and *SVP* clades are necessary for normal inflorescence development, and that this mechanism may be conserved in monocot and dicot species.

### *SVP* and *SQUAMOSA* genes contribute to peduncle elongation and plant height

In addition to their positive interactions in accelerating the transitions of apical meristems, wheat *SQUAMOSA*- and *SVP*-genes contribute to stem elongation. Mutants for all three wheat *SQUAMOSA* genes have shorter stems, with the triple mutants being shorter than any other mutant combinations (Li et al., 2019). Significant reductions in plant height have been also reported for the mutants of the *SQUAMOSA* orthologs in rice (*osmads14* and *osmads15*), suggesting a conserved function in grasses (Wu et al., 2017).

A conserved role in promoting stem elongation has been reported also for genes in the *SVP* clade in several grass species. Increases in plant height were observed in transgenic plants constitutively expressing *BM1* in barley (Trevaskis et al., 2007) or *OsMADS55* in rice (Lee et al., 2008), and in a natural *VRT-A2* high-expression allele from *T. turgidum* subsp. *polonicum* (Adamski et al., 2020). The complementary reduction in plant height was observed in non-functional *vrt2* and *svp1* mutants in tetraploid wheat (Figure 3F) and in RNAi transgenic rice plants with reduced transcript levels of *OsMADS55* (~*VRT2*) and *OsMADS47* (~*SVP3*) (Lee et al., 2008).

In the complementation experiment, we detected a significant increase in peduncle length for *UBI::VRT2* in the *vrt2* background (140%, *P* < 0.0001), but no significant difference was detected in the WT background (Supplemental Figure 4B). Similarly, a slight increase in stem length (2.5 cm, *t*-test *P* = 0.055) was observed for the weak *UBI::VRT2* transgene T#8 relative to the control in the *ful2* mutant background but not in the presence of the wild type *Ful2* allele (Figure 7A). These results suggest that even though the positive effect of *SVP* genes on peduncle elongation is conserved across grass species, constitutive expression or expression above certain threshold levels may be no longer effective. Since SQUAMOSA genes also promote stem elongation, we speculate that interactions between these two groups of grass MADS-box genes, possibly through SQUAMOSA-SVP protein complexes, may explain the drastic reduction in stem elongation in the *vrt2 vrn1 ful2* triple mutant.

### *SVP* genes repress axillary meristems in elongating stems

An unexpected phenotype of the *vrt2 svp1* mutant was the development of axillary spikelets or spikes in the nodes of the elongating stem (Figure 4E-J and N). We also observed some axillary spikes in the single *vrt2* and *svp1* mutants (Figure 4K and L), but they were less frequent and less developed than in *vrt2 svp1* (Figure 4M). These results indicate that both *VRT2* and *SVP1* function redundantly as repressors of axillary meristems in the nodes of the elongating stem.

The *vrt2 svp1* axillary spikes are located in the same position as the ears in a maize plant, or the axillary inflorescences or “paracladia” in species from the Bambusoideae, Panicoideae and Andropogoneae families (Stapleton, 1997; Vegetti, 1999). The phylogenetic distribution of this trait, together with the *vrt2 svp1* results presented in this study suggest that axillary inflorescence formation is likely an ancestral trait in grasses that is likely repressed in many modern grass species.

Andropogoneae species can develop large axillary inflorescences or small ones consisting of pairs of spikelets or single spikelets, a variation similar to the one we observed in the *vrt2 svp1* mutant from the nodes closest to the spike to the basal nodes of the elongating stem (Figure 4F). These results, together with those from the companion paper (Adamski et al., 2020), provide good examples of changes in conserved developmental genes that generate different inflorescence architectures and spikelet morphologies that parallel morphological differences observed during grass evolution.

### Failure to repress *SVP* genes results in spikelets with vegetative characteristics, branches and increased number of florets

Ectopic expression of *SVP* genes has been associated with vegetative characteristic in spikelets in barley plants constitutively expressing *BM1* (~*SVP3*) or *BM10* (*~SVP1*) (Trevaskis et al., 2007), rice plants overexpressing *OsMADS22* (~*SVP1*) (Sentoku et al., 2005), maize plants carrying the dominant *Tunicate1* pod corn mutation (Han et al., 2012; Wingen et al., 2012), and our tetraploid wheat lines constitutively expressing *VRT2* (Figure 5C). Interestingly, *UBI::VRT2* T#4 plants have a spikelet phenotype that resembles *T. turgidum* subsp. *polonicum*. The companion paper shows that the long and leafy glumes of *T. turgidum* subsp. *polonicum* are the result of natural variation in a regulatory region of *VRT-A2* that increases transcript levels and alters the tissues where this gene is normally expressed (Adamski et al., 2020). The role of *VRT2* in these phenotypes is supported in our study by the reduced vegetative characteristics of *vrt2 <u>Vrn1</u> ful2* mutant compared with *<u>Vrn1</u> ful2* (Figure 8) and the enhanced vegetative characteristics associated with the combined *ful2 UBI::VRT2* alleles relative to their individual effects (Figure 7). The highly significant interaction between *FUL2* and *UBI::VRT2* in glume and lemma length (Supplemental Table 8) indicates a genetic interaction between these two genes in the determination of vegetative characteristics in the spikelet.

Although mutations in multiple *SQUAMOSA* genes or constitutive expression of *SVP* genes can both lead to spikelets with vegetative characteristics, the *SQUAMOSA* genes seem to play a more essential role in SM identity. Whereas the wheat *vrt2 svp1* double mutant had normal and fertile spikelets, the *vrn1 ful2* mutants formed tillers instead of spikelets and the addition of *vrt2* was insufficient to restore spikelet identity in the *vrt2 vrn1 ful2* triple mutant (Supplemental Figure 5D).

We currently do not know if the *SVP-*clade genes actively induce vegetative characteristics or if these are an indirect effect of the repression of floral organs and the regression to a “default” vegetative development program. We have shown here that constitutive expression of *VRT2* results in the down-regulation of MADS-box A-, B-, C- and most E-class flowering genes, a function conserved in Arabidopsis (Gregis et al., 2009; Liu et al., 2009). Moreover, continuous expression of *VRT2* can compete with the formation of SQUAMOSA-LOFSEP complexes (Figure 10), which are important for normal floral development. Since *SEPALLATA* triple mutant *osmads1 osmads5* and *osmads34* in rice has leaf-like lemmas and paleas (Wu et al., 2018), we speculate that the downregulation of these floral genes or their reduced activity by protein competition may contribute to the observed vegetative characteristics in *UBI::VRT2* and *<u>Vrn1</u> ful2* plants.

In addition to the vegetative characteristics of paleas and lemmas, high levels of constitutive expression of *VRT2* in T#2 transgenic plants resulted in the transformation of some basal spikelets into branches. These branches had lateral and terminal spikelets and resembled small spikes (Figure 5F). This phenotype was also observed in plants with the combined *ful2 UBI::VRT2* T#8 alleles but not in those with the individual T#8 or *ful2* alleles (Figure 7). The antagonistic interaction between the *SQUAMOSA* and *SVP* genes on spikelet development was also evident in the reduction in the proportion of spikelets with branches from 80% in *<u>Vrn1</u> ful2* to 32% in *vrt2 <u>Vrn1</u> ful2* (Supplemental Figure 6F). Together, these results suggest that dynamic changes in the relative abundance of *SQUAMOSA* and *SVP* genes are critical for the normal progression of the lateral floret meristems within the spikelet.

The *SQUAMOSA* and *SVP* genes also showed opposite effects in the regulation of the number of florets per spikelet. The large number of florets observed in *<u>Vrn1</u> ful2* (Figure 8E) was reduced dramatically in the *vrt2 <u>Vrn1</u> ful2* (Figure 8F and I); whereas that number increased in the *ful2 UBI::VRT2* plants relative to the individual mutant or transgenic plants (Figure 7). While *VRN1* and *FUL2* negatively control the number of florets per spikelet, ectopic expression of *VRT2* prolonged the activity of the SM and the production of lateral FM. This *SVP* function seems to be conserved in other grass species because overexpression of *OsMADS22* increased the number of florets per spikelet from one to two in some of the rice transgenic plants (Sentoku et al., 2005).

Large increases in the number of florets per spike have been reported in wheat loss of function mutants of the *Q* gene (=*AP2L5*) and in plants with constitutive expression of *miRNA172*, which down-regulates *Q* and other *AP2L* genes (Debernardi et al., 2017). Moreover, plants combining *ful2* and *UBI::miRNA172* showed a large increase in the number of florets per spike (Li et al., 2019), similar to the combined *ful2 UBI::VRT2* in this study. Since Q was identified as a VRT2 interactor in a Y2H screening (Kane et al., 2005), it would be interesting to investigate the genetic interactions between genes in the SVP-clade and those down-regulated by miR172 (e.g. Q) on the regulation of floret number per spikelet.

### Dynamic changes in expression of *SVP*-,*SQUAMOSA-* and *SEPALLATA-*genes are important for normal wheat spike and floral development

Based on the results from this and a previous study (Li et al., 2019), we propose the following working model for the roles of SQUAMOSA and SVP MADS-box genes in the regulation of wheat reproductive development (Figure 11). Initially, *VRT2* and *SVP1* contribute to the acceleration of the transition of the vegetative SAM to an IM, which in wheat is driven mainly by the induction of *VRN1* (Yan et al., 2003; Loukoianov et al., 2005). Then, genes from the SQUAMOSA- and SVP-clades share a common role in accelerating the transition of the IM to a terminal spikelet, with mutants in both groups of genes resulting in increased SNS.

**Figure 11.**
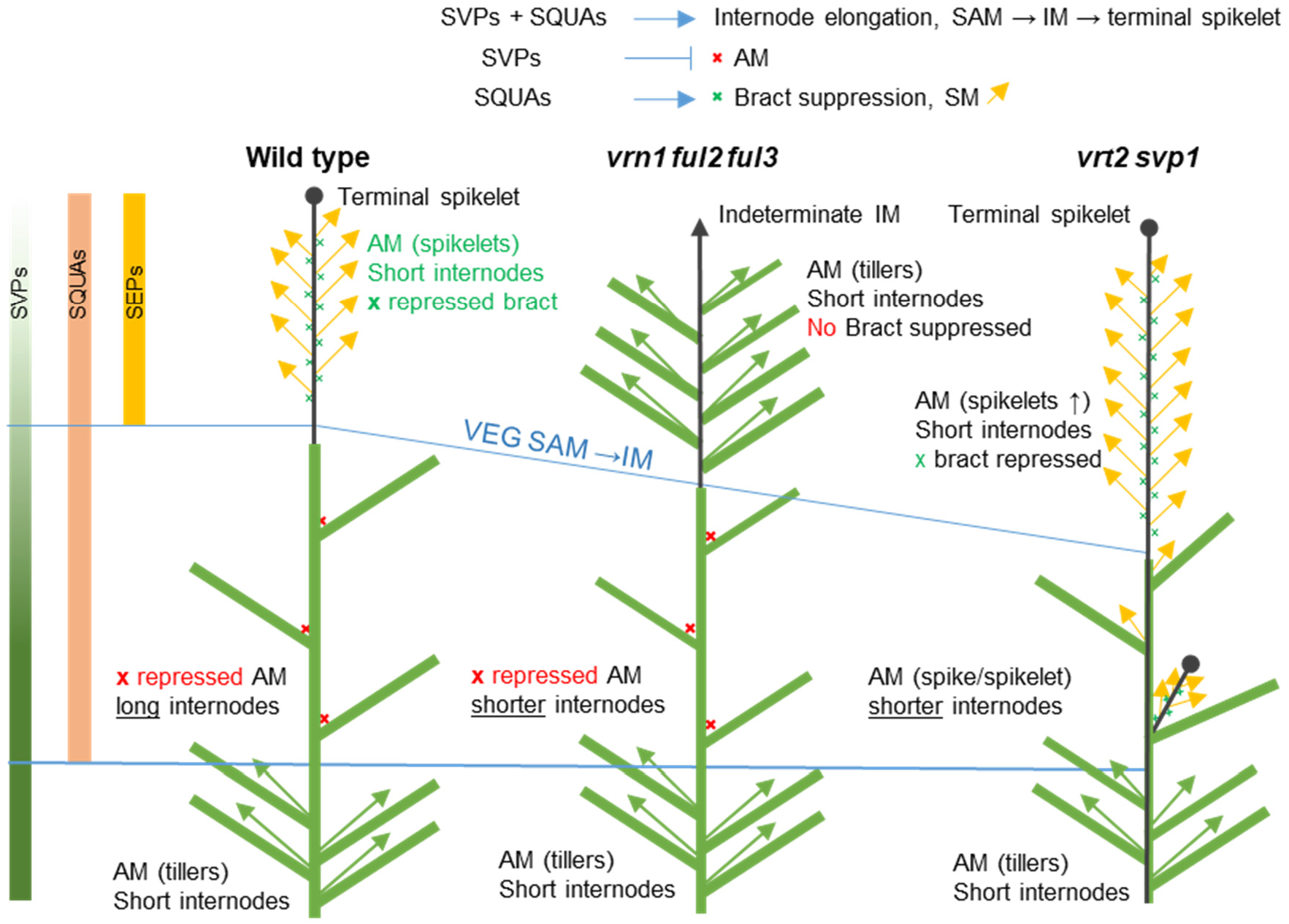
Working model of the role of *SVP* (*VRT2* and *SVP1*), *SQUAMOSA* (*VRN1*, *FUL2* and *FUL3*) and *SEPALLATA* genes on the regulation of wheat plant architecture. Bars on the left represent transcript levels of genes from the three MADS-box clades during wheat development. The three plant models represent the architecture of WT, *vrn1 ful2 ful3* triple and *vrt2 svp1* double mutants. Green rectangles represent leaves and sheaths, black lines the spike rachis (circle end = determinate, arrow end = indeterminate). The green Xs represent repressed bracts in the spike and the red Xs repressed buds in the elongating nodes. The lower part of the plants represents vegetative growth (*SVP* genes only), the region between the two blue lines the elongation zone (central region, *SVP* + *SQUAMOSA* genes) and the region above the blue lines the developing spikes (*SQUAMOSA* + *SEPALLATA* genes). In *vrn1 ful2 ful3*, the lateral spikelet meristems regress to vegetative meristems and the bracts are not suppressed. In *vrt2 svp1*, the axillary buds in the elongation zone are no longer repressed and develop into axillary spikes or spikelets.

Genes from both groups also contribute to normal stem elongation, a process that also requires the induction of *FT1* in the leaves (Pearce et al., 2013). In the nodes of the elongating stem, *VRT2* and *SVP1* play a previously unreported role in the repression of axillary meristems (Figure 11, right), which contributes to the formation of a single terminal inflorescence.

In the developing spike, the down-regulation of the *SVP-*genes by SQUAMOSA proteins at the PDR and subsequent stages is critical for SM specification and normal spikelet development. The repression of *SVP*-genes before floral development is also conserved in Arabidopsis (Yu et al., 2004; Liu et al., 2007), where ChIP experiments have demonstrated that AP1 and SEP3 act as direct repressors of *AGL24* and *SVP* (Gregis et al., 2008).

MADS-box proteins can form dimeric and tetrameric complexes providing flexibility for the formation of different complexes as the abundance of different MADS-box proteins changes through development (Theissen et al., 2016). Our Y3H results in wheat suggest that the down-regulation of the *SVP* genes has the potential to reduce the competition of their encoded proteins with the formation of LOFSEP - SQUAMOSA protein complexes required for normal spikelet and floret development. Through interactions with both the SVP and SEPALLATA proteins, the SQUAMOSA proteins play a pivotal role in the sequential transition between MADS-box protein complexes favoring vegetative characteristics and those favoring the development of floral organs. A similar function as a protein interaction hub between the flower induction pathway (e.g. SVP, AGL24, and SOC1) and the floral organ identity proteins was proposed in Arabidopsis for the SQUAMOSA proteins AP1 and FUL2 (de Folter et al., 2005).

In summary, this study shows that the wheat *SVP* and *SQUAMOSA* genes have synergistic effects on the acceleration of the transitions of apical meristems (SAM → IM → terminal spikelet) and stem elongation, but antagonistic effects on the regulation of lateral meristems in spikes and spikelets, with *SQUAMOSA* genes promoting the transition to floral organs and *SVP* genes having a regressive effect on this transition. In addition, this study reveals a previously unknown role of the *SVP* genes in the repression of axillary meristems in the elongating stem that has drastic effects on wheat inflorescence architecture, and that has interesting implications in the evolution of grass inflorescences. This study and the companion paper (Adamski et al., 2020), show that *SVP* genes affect important agronomic traits such as grain test weight and number of spikelets per spike, which are valuable targets for wheat improvement. However, since these genes have multiple pleiotropic effects, the correct dosage for optimum productivity will have to be adjusted in different environments. Our results for the individual *vrt2* and *svp1*, and their individual homeolog mutants, show that these effects can be readily fine-tuned in a polyploid species like wheat.

## MATERIALS AND METHODS

### Quant-Seq of *vrn1* and *vrn1 ful2* developing spikes

We collected shoot apical meristems from *vrn1* and *vrn1 ful2* mutants at four developmental stages: vegetative (VEG), double-ridge (DR), post-double-ridge (PDR) and terminal spikelet (TS) (Figure 1A). We performed Quant-Seq analysis using four biological replicates for each of the four developmental stages, with each replicate including pools of 6 apices for PDR and TS stages, 9 apices for DR and 12 apices for vegetative apices. Sequencing of the 32 samples (2 genotypes × 4 developmental stages × 4 biological replicates) using Hi-seq (100 bp reads not paired) yielded an average of 7,335,215 unique reads per sample after filtering for duplicates, with an average read length of 74.3 bp (after trimming) and an average quality of 36.4 (Supplemental Table 2). BioProject numbers are available in the Material and Methods section Accession Numbers.

We processed the raw reads using DOE JGI BBTools (https://sourceforge.net/projects/bbmap/) program bbduk.sh to remove Illumina adapter contamination and low-quality reads (forcetrimleft=21 qtrim=r trimq=10). Processed reads were mapped to the IWGSC RefSeq v1.0 genome assembly, using the STAR aligner (Dobin et al., 2013). We used parameters -- outSAMtype BAM SortedByCoordinate --outSAMunmapped Within --outSAMattributes Standard --quantMode TranscriptomeSAM GeneCounts to generate Binary Sequence Alignment/Map (BAM) files for each sample. We used the high confidence gene models from IWGSC Refseq v1.0 (IWGSC_v1.1_HC_20170706.gff) in combination with the BAM files in the R program (featureCounts.R) which uses the Rsubread package (Liao et al., 2019) to calculate the overlap between reads and features. We used the “readExtension5” option that allows a read to be counted as belonging to a gene when the gene was a defined number of bases 5’ of the read (we used 500 bp).

The raw *t*-test values between read counts of *vrn1* and *vrn1 ful2* were corrected for false discovery rate (FDR) using the R function p.adjust (method = ‘BH’, aka ‘FDR”) (Benjamini and Hochberg, 1995; R Core Team, 2018). Differentially expressed genes (DEGs) between *vrn1* and *vrn1 ful2* mutants for each stage were defined as those with a fold change in transcript levels ≥ 2 and FDR ≤ 0.05. We then generated a list of non-redundant down-regulated and up-regulated DEGs across the four stages and performed a cluster analysis based on their expression profiles using the MultiExperiment Viewer (MeV) software (www.tm4.org). For this analysis, the expression levels of each DEG was normalized to the average expression value of each DEG across genotypes and stages (mean normalized expression), and then clustered using K-means with a minimum limit of 10% of total genes per cluster. We then blasted the clustered lists against a rice gene database available from Phytozome (Osativa_323_V7.0.cds_primaryTranscriptOnly.fa) to obtain a functional annotation for the DEGs (Supplemental File 1). Finally, the lists containing the best rice blast hits were used to perform GO enrichment analysis using AgriGO web tool (http://bioinfo.cau.edu.cn/agriGO/).

### Identification of loss-of-function mutations in *VRT2*

The sequenced ethyl methane sulphonate (EMS) mutagenized populations of the tetraploid wheat variety Kronos and hexaploid variety Cadenza were screened for mutations using BLASTN with the sequences for *VRT2* sequence (*TraesCS7A02G175200* and *TraesCS7B02G080300*) and *SVP1* (*TraesCS6A02G313800* and *TraesCS6B02G343900*) as queries (Krasileva et al.,2017). For *VRT-A2*, we detected 46 mutations that generated amino acid changes but we found no truncation mutations in the Kronos mutant population. Therefore, we screened the mutant population of the hexaploid wheat Cadenza, where we identified a mutation that generated a Q125* premature stop codon in mutant line Ca0424, which is predicted to eliminate 47% of the VRT2 protein, including part of the K-box and C terminal domains (Figure 2A, above gene model). Since Kronos × Cadenza crosses result in hybrid necrosis, we crossed Ca0424 to an F_2_ plant from the cross between the hexaploid Insignia and the tetraploid Kronos as a bridge cross. We then intercross the F_1_ with the *vrt-B2* mutant K3404 to combine both mutations (Figure 2C).

The Kronos mutant line K3404 carries a mutation in the donor splice site of the fourth intron of the *VRT-B2* gene, designated hereafter as *vrt-B2* (Figure 2A, below gene model). Sequencing of the RT-PCR products from K3404 revealed three *vrt-B2* alternative splice forms, all resulting in severe truncations. The first alternative splicing form showed a five bp insertion as a result of the utilization of the next available GT splicing site in intron 4. This resulted in a reading frame shift and a premature stop codon that is predicted to eliminate half of the protein including 30% of the conserved K domain. The second alternative splicing form had an insertion of the last 4 bp of intron four between exons four and five, which generated a reading frame shift and an early stop codon. Similar to the first alternative splice form, this change is also predicted to eliminate half of the protein and 30% of the K domain. Finally, the third alternative splicing form was missing exons 3, 4 and 5. Although exons 6, 7 and 8 retained the correct reading frame, the deletion resulted in the elimination of 80% of the K domain (Supplemental Figure 2).

To reduce the background mutations, we backcrossed the F_1_ plant from the cross between *vrt-A2* and *vrt-B2* three times to Kronos, and from the segregating BC_2_F_2_ plants we selected a double homozygous mutant *vrt-A2 vrt-B2*, which was designated as *vrt2*. To test the genetic interaction between *vrt2* and members of the MADS-box genes from the *SQUAMOSA*-clade (*VRN1* and *FUL2*), we combined *vrt2* with loss-of-function mutations at *vrn1 ful2* in the same Kronos background (Li et al., 2019). Since the *vrn1 ful2* line is sterile, we used a line heterozygous for *Vrn-A1* and homozygous for all the other truncation mutations to make the cross (*Vrn-A1*/*vrn-A1*, *vrn-B1*, *ful-A2*, and *ful-B2*, designated as *<u>Vrn1</u> ful2*). All these mutant lines were developed in a Kronos background with no functional copies of *VRN2* to avoid the extremely late heading of the *vrn1* mutants in the presence of *VRN2,* which is a strong flowering repressor in wheat (Distelfeld et al., 2009). We self-pollinated the F_1_ plant and from the F_2_ plants we selected lines homozygous for *vrn1 ful2* and either homozygous for *vrt2* or for the WT alleles (Figure 2C). Since the *vrt2 vrn1 ful2* mutants failed to form spikelets, we also selected lines *<u>Vrn1</u> ful2* with and without *vrt2* to study its effect on spike morphology.

### Identification of loss-of-function mutations in *SVP1*

For *SVP-A1*, we identified the Kronos line K4488 that carries a mutation in the splice donor-site in the third intron, designated as *svp-A1* (Figure 2B). The sequencing of *SVP-A1* RT-PCR products from K4488 revealed two alternative splicing forms. The first one lacks the third exon, which alters the reading frame and generates a premature stop codon that eliminates >60% of the SVP-A1 protein. Since this deletion includes the complete K domain, the resulting protein is likely not functional. The second alternative splice form lacks both the second and third exons, which results in the loss of 47 amino acids but does not alter the reading frame. Since this predicted deletion includes the end of the MADS domain and the beginning of the K domain, the resulting protein is likely not functional.

For *SVP-B1*, we identified Kronos line K0679 with a mutation that encodes an early stop codon in the third exon (Q99*), designated as *svp-B1* (Figure 2B). We crossed both mutants separately two times to the parental Kronos to reduce background mutations (BC_1_) and then combined them by crossing and selection in BC_1_F_2_, to generate the double mutant designated *svp1*. Finally, we intercrossed *svp1* and *vrt2*, self-pollinated the F_1_, and selected F_2_ plants homozygous for the four mutations (*vrt-A2 vrt-B2 svp-A1 svp-B1*) which were designated as *vrt2 svp1* (Figure 2C).

### Transgenic plants and complementation

Transgenic Kronos plants overexpressing *VRT2* were generated at the UC Davis Plant Transformation Facility (http://ucdptf.ucdavis.edu/) using the Japan Tobacco (JT) vector pLC41 (hygromycin resistance) and transformation technology licensed to UC Davis. The coding region of *VRT-A*^*m*^*2* gene from *Triticum monococcum* accession G3116 (GenBank MW218446) was cloned downstream of the maize *UBIQUITIN* promoter with a C-terminal 4×MYC tag (henceforth *UBI::VRT2*). *Agrobacterium* strain EHA105 was used to infect Kronos immature embryos and all transgenic plants were tested by PCR using primers described in Supplemental Table 4.

To test complementation of the mutant phenotypes, we crossed *UBI::VRT2* plants with *vrt2* mutants. We self-pollinated the F_1_ plants and used molecular markers to select F_2_ plants homozygous for the *vrt-A2* and *vrt-B2* mutations with and without the transgene. These sister lines were evaluated in growth chamber as described in the following section.

### Growth conditions and phenotyping

We phenotyped mutant and transgenic plants for heading time, spikelet number per spike (SNS), stem length, and spike and floral defects in PGR15 CONVIRON growth chambers under long-day photoperiod (16 h of light and 8 h dark) and temperatures of 22 °C during the day and 18 °C during the night. The light intensity of the sodium halide lights was approximately 330 μMm^−2^s^−1^. Plants were germinated in petri dishes at 4 °C for 3 to 5 days. After the first leaf emerged, we transplanted the seedlings into the soil in one-gallon pots, and recorded days to heading from this day until emergency of half of the main spike from the flag leaf. Length measurements were taken at maturity for the complete plants and for each of the internodes and peduncle separately.

We also evaluated the *vrt2* Kronos mutants in a field experiment sown November 22, 2019 at the UC Experimental Field Station in Davis, CA (38° 32′ N, 121° 46′ W). We used one-meter rows with 20 plants each as experimental units, organized in a completely randomized design. The experiment included 20 replications for each of the *vrt2* double mutants and the WT sister lines and 10 replications for each of the single mutants *vrt-A2* and *vrt-B2.* Plants in the field were evaluated from heading time, SNS and total plant height (measured from the soil to the top of the main spike excluding awns).

### Statistical analyses

Effects of individual homeologs (A and B genome) and their interactions or of individual genes and their interactions in double mutants were compared using 2 × 2 factorial ANOVAs with genes or homeologs as factors and alleles as levels. Simple effects were evaluated using orthogonal contrasts. Means of the individual genotypes were compared with the wild type using Dunnett tests. Homogeneity of variances was tested with the Levene’s test and normality of residuals with the Shapiro-Wilks test. When necessary, we transformed data to meet the assumptions of the ANOVA. All statistical analyses were performed using SAS version 9.4. Distribution of the data within each genotype are presented with box-plots including individual data points generated with Excel. The middle line of the box represents the median and the × represents the mean. The bottom line of the box represents the first quartile and the top line the third quartile. The whiskers extend from the ends of the box to the minimum value and maximum value. A data point was considered an outlier if it exceeded a distance of 1.5 times the inter quartile range. The number of plants analyzed is indicated in each graph.

### cDNA Preparation and qRT-PCR Analysis

To quantify transcript levels of different flowering genes in transgenic plants overexpressing VRT2, we extracted total RNA from pools of 6-8 shoot apical meristems (SAM) after the terminal spikelet stage from four biological replicates using the Spectrum Plant Total RNA Kit (Sigma-Aldrich). The cDNA was synthesized using the High-Capacity cDNA Reverse Transcription Kit (Thermo Fisher Scientific, 4368814) from 2 μg RNA treated with RQ1 RNase-free DNase (Promega). The cDNA was then diluted 20 times in water and 5 μl of the dilution was used for the realtime qPCR analysis. The Quantitative PCR was performed using the 7500 Fast Real-Time PCR system (Applied Biosystems) with 2×VeriQuest Fast SYBRGreen qPCRMaster Mix (Affymetrix, 75690). The relative transcript level was determined for each sample and normalized using *ACTIN* as an endogenous control. The normalization was performed as described previously (Livak and Schmittgen, 2001). Melting curve analyses at the end of the process and “no template controls” were performed to ensure product-specific amplification without primer-dimer artifacts. Primer sequences are given in Supplemental Table 4.

### Yeast-Two-Hybrid (Y2H) assay and Bimolecular Fluorescence Complementation (BiFC)

We used the GAL4-based Y2H system to investigate protein interactions. We amplified the full-length cDNAs of the different genes from the *SQUAMOSA-* (*VRN1*, *FUL2* and *FUL3*), *SVP-* (*VRT2*, *SVP1*, *SVP3*) and *SEPALLATA*-clades (*SEP1-2*, *SEP1-4*, and *SEP1-6*) and cloned them into the gateway™ pDONR™/Zeo Vector (Catalog number: 12535035) using primers listed in Supplemental Table 4. We then cloned these genes into Y2H vectors pGADT7 (activation-domain vector) and pGBKT7 (DNA-binding domain vector) by either restriction enzyme-based cloning or In-Fusion HD Cloning method (638910In-Fusion^®^ HD Cloning Plus Takara). Primer sequences used to generate Y2H and Y3H constructs are listed in Supplemental Table 4. Both bait and prey vectors were transformed into yeast AH109 gold strain. The co-transformants were plated on selective solid Synthetic Dropout agar medium without leucine (L) and tryptophan (W) (SD-L-W). Positive transformants were re-plated on Synthetic Dropout medium lacking L, W, histidine (H) and adenine (A) to test for interaction (SD-L-W-H-A). We co-transformed each bait vector with pGADT7 and each prey vector with pGBKT7 to test autoactivation.

For the bi-molecular fluorescent complementation (BiFC or split YFP) assays, we cloned the same genes into modified Gateway-compatible vectors UBI::NYFP-GW and UBI::CYFP-GW by recombination reactions (UBI= *UBIQUITIN* promoter). These vectors generated fusion proteins with YFP-N-terminal fragment or YFP-C-terminal-fragment at the N-terminus and the proteins being tested at the C-terminus. Wheat protoplasts were prepared, transfected and visualized as described in (Shan et al., 2014).

### Yeast Three-Hybrid (Y3H) assays

The pBridge yeast three-hybrid system (Clontech CATALOG No. 630404) was used to test if the wheat VRT2 protein can interfere with the interactions between SEPALLATA and SQUAMOSA proteins. This vector can express two proteins, a DNA-binding domain fusion, and a second protein (Bridge protein) that is controlled by pMET25, an inducible promoter responsive to methionine levels in the medium. The Bridge protein is only expressed in the absence of methionine and inhibited by the addition of 1 mM methionine. For each pBridge vector, one of the *LOFSEPs* (*SEP1-2, SEP1-4* or *SEP1-6*) genes was fused to the DNA-binding domain, and *VRT2* gene was inserted downstream of the MET25 promoter. The same prey vectors generated for *VRN1*, *FUL2* and *FUL3* in Y2H assays were used in Y3H assays. Each pBridge vector was then paired with one prey vector and co-transformed into yeast Gold. Transformants containing both vectors were selected on SD-L-W medium. Protein interactions were quantified using quantitative α-galactosidase assays as described before (Li et al., 2011).

All constructs used in Y2H and Y3H assays have the GAL4 DNA binding (bait) and activation domains (prey) at the N-terminus, and the proteins being tested at the C-terminus of the fusion protein.

### Accession numbers

The *T. monococcum VRT-A*^*m*^*2* sequence used for the constitutive expression construct is deposited in GenBank under accession number MW218446. The Quant-Seq datasets for the *vrn1* and *vrn1ful2* mutants have been deposited in GenBank under the following project numbers (each including four biological replicates): PRJNA681065 (*vrn1*, vegetative samples), PRJNA681067 (*vrn1*, double ridge), PRJNA681097 (*vrn1*, post double ridge) PRJNA681099 (*vrn1*, terminal spikelet), PRJNA681036 (*vrn1 ful2*, vegetative samples), PRJNA680890 (*vrn1 ful2*. double ridge), PRJNA681027 (*vrn1 ful2*, post double ridge), PRJNA681032 (*vrn1 ful2*, terminal spikelet).

## Supporting information

Supplemental Quant-Seq Data

## Supplemental data Files

The Supplemental Information file includes 10 supplemental figures and 9 supplemental tables. Supplemental data file 1: Differentially expressed genes from Quant-Seq data analysis.

## ACKNOWLEDGEMENTS

This project was supported by the Howard Hughes Medical Institute, NRI Competitive Grant 2016-67013-24617 and 2017-67007-25939 from the USDA National Institute of Food and Agriculture (NIFA). We thank Hans Vasquez-Gross, German F. Burguener and Junli Zhang (UC Davis) for the Quant-Seq analyses and deposit of the data into GenBank. We also thank Kevin Childs and Jose Planta from Michigan State University for advice and sharing their pipeline.

## AUTHOR CONTRIBUTIONS

CL, JMD and JD designed the research. KL performed most of the experimental work. JMD, CL, HL and CZ performed research, CL, HL, JMD, KL, and JD analyzed the data. CL, JMD, KL, HL and JD wrote the paper.

## DATA STATEMENT

The Quant-Seq data has been deposited in GenBank under accession number (pending). The *T. monococcum VRT-A*^*m*^*2* sequence used for the constitutive expression construct is deposited in GenBank under accession number MW218446. All other data and genetic materials are available from the authors upon request.

