## Supplemental Figures for "Interactions Between SQUAMOSA and SVP MADS-box Proteins Regulate Meristem Transitions During Wheat Spike Development"

### SUPPLEMENTAL INFORMATION

#### SUPPLEMENTAL FIGURES

**Supplemental Figure 1.** Phylogenetic relationship among SVP proteins in wheat, barley, rice and Arabidopsis.

We inferred the evolutionary history of MADS-box SVP proteins from wheat, barely, rice and Arabidopsis using the Neighbor-Joining method. We used the MUSCLE protein alignment presented in the next page to calculate the optimal tree with branch lengths proportional to the distances used to infer the phylogenetic tree. The evolutionary distances were computed using the Poisson correction method, and are in the units of the number of amino acid substitutions per site. We show the percentage of replicate trees in which the associated taxa clustered together in the bootstrap test (1000 replicates) next to the branches. We removed all ambiguous positions for each sequence pair (pairwise deletion option) and the final dataset included 236 positions. We conducted all the analyses using MEGA X (Kumar et al., 2018, Mol. Biol. Evol. 35**:**1547-1549). Arabidopsis SVP and AGL24 proteins were included as outgroups of the grass SVP proteins.

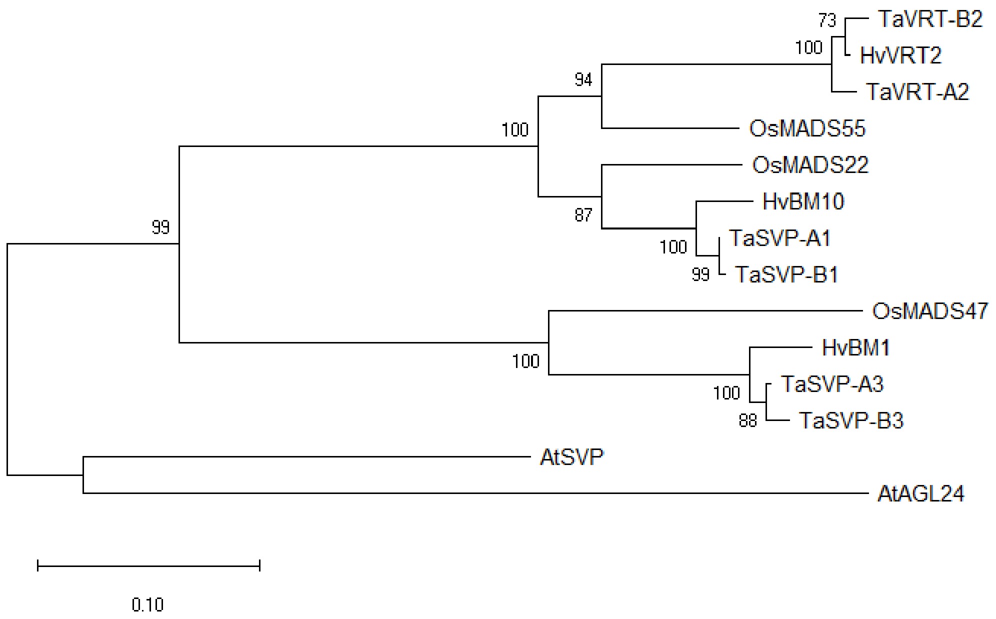


Below we present the tree in Newick machine-readable format:

(((((TaVRT-B2,HvVRT2),TaVRT-A2),OsMADS55),(OsMADS22,(HvBM10,(TaSVP-A1,TaSVP-B1)))),(OsMADS47,(HvBM1,(TaSVP-A3,TaSVP-B3))),(AtSVP,AtAGL24));

Alignment of *Triticum aestivum* (Ta), *Hordeum vulgare* (Hv), *Oryza sativa* (Os) and *Arabidopsis thaliana* (At) MADS-box proteins from the SVP-clade using MUSCLE in MEGA X. Poorly aligned regions at the beginning and end of the proteins were truncated and were not used in the calculation of the phylogenetic tree described in the previous page. We indicate the three different grass orthologous groups with different colors.

```
TaVRT-A2      1 MARERRAIRRIESAAARQVTFSKRRRGLFKKAEELAVLCDADVALVVFSSTGKLSQFASSSMNEIIDKYSTHSKNLGKSD
TaVRT-B2      1 MARERRAIRRIESAAARQVTFSKRRRGLFKKAEELAVLCDADVALVVFSSTGKLSQFASSSMNEIIDKYSTHSKNLGKSD
HvVRT2        1 MARERRAIRRIESAAARQVTFSKRRRGLFKKAEELAVLCDADVALVVFSSTGKLSQFASSSMNEIIDKYSTHSKNLGKSD
OsMADS55      1 MARERREIRRIESAAARQVTFSKRRRGLFKKAEELAVLCDADVALVVFSSTGKLSQFASSNMNEIIDKYTTHSKNLGKTD
TaSVP-A1      1 MARERREIKRIESAAARQVTFSKRRRGLFKKAEELSVLCDADVALIVFSSTGKLSQFASSSMNEIIDKYSTHSKNLGKT-
TaSVP-B1      1 MARERREIKRIESAAARQVTFSKRRRGLFKKAEELSVLCDADVALIVFSSTGKLSQFASSSMNEIIDKYSTHSKNLGKT-
HvBM10        1 MARERREIKRIESAAARQVTFSKRRRGLFKKAEELSVLCDADVALIVFSSTGKLSQFASSSMNEIIDKYSTHSKNLGKT-
OsMADS22      1 MARERREIKRIESAAARQVTFSKRRRGLFKKAEELSVLCDADVALIVFSSTGKLSHFASSSMNEIIDKYNTHSNNLGKA-
HvBM1         1 GKRERIAIRRIENLAARQVTFSKRRRGLFKKAEELSILCDAEVGLAVFSATGKLFQFASSSMNQIIDRYNSHSKILKKVD
OsMADS47      1 GKRERIAIRRIDNLAARQVTFSKRRRGLFKKAEELSILCDAEVGLVVFSATGKLFQFASTSMEQIIDRYNSHSKTLQRA-
TaSVP-A3      1 GRRERIAIRRIENLAARQVTFSKRRRGLFKKAEELSILCDAEVGLAVFSATGKLFQFASSSMNQIIDRYNSHSKILKKAD
TaSVP-B3      1 GKRERIAIRRIENLAARQVTFSKRRRGLFKKAGELSILCDAEVGLAVFSATGKLFQFASSSMNQIIDRYNSHSKILKKAD
AtSVP         1 MAREKIQIRKIDNATARQVTFSKRRRGLFKKAEELSVLCDADVALIIFSSTGKLFEFCSSSMKEVLERHNLQSKNLEKL-
AtAGL24       1 MAREKIRIKKIDNITARQVTFSKRRRGIFKKADELSVLCDADVALIIFSATGKLFEFSSSRMRDILGRYSLHASNINKLM

TaVRT-A2     81 QQPAIDLN-LEHCKYDSLNEQLAEASLRLRHMRGEELDGLSVGELQQMEKNLETGLQRVLCTKDRQFMQQISDLQQKGTQ
TaVRT-B2     81 QQPAIDLN-LEHCKYDSLNEQLAEASLRLRRMRGEELDGLSVGELQQMEKNLETGLQRVLCTKDRQFMQQINDLQQKGTQ
HvVRT2       81 QQPAIDLN-LEHCKYDSLNEQLAEASLRLRHMRGEELDGLSVGELQQMEKNLETGLQRVLCTKDRQFMQQISDLQQKGTQ
OsMADS55     81 KQPSIDLN-LEHSKCSSLNEQLAEASLQLRQMRGEELEGLSVEELQQMEKNLEAGLQRVLCTKDQQFMQEISELQRKGIQ
TaSVP-A1     80 DQPALDLN-LEHSKYANLNDQLAEASLRLRQMRGEELEGLSVDELQQLEKNLETGLHRVLQTKDQQFLEQINELHRKSSQ
TaSVP-B1     80 DQPALDLN-LEHSKYANLNDQLAEASLRLRQMRGEELEGLSVDELQQLEKNLETGLHRVLQTKDQQFLEQINELHRKSSQ
HvBM10       80 DQPTLDLN-LEHSKYANLNDQLAEASLRLRQMRGEELEGLSVDELQQLEKNLETGLHKVLQTKDQQFLEQINELHRKSSQ
OsMADS22     80 EQPSLDLN-LEHSKYAHLNEQLAEASLRLRQMRGEELEGLSIDELQQLEKNLEAGLHRVMLTKDQQFMEQISELQRKSSQ
HvBM1        81 EPSQLDLH--EDSNCARLRDELAEASLWLQQMRGEELQSLNVQQLQALEKSLESGLSSVLKTKSQKIMDQISELEKKRVQ
OsMADS47     80 EPSQLDLQGEDSSTCARLKEELAETSLRLRQMRGEELHRLNVEQLQELEKSLESGLGSVLKTKSKKILDEIDGLERKRMQ
TaSVP-A3     81 EPSQLDLH--EDSNCARLRDELAEASLWLQQMRGEELQSLNVQQLQALEKSLESGLGSVLKTKSQKIMDQISELERKRVQ
TaSVP-B3     81 EPSQLDLH--EDSNCARLRDELAEASLWLQQMRGEELQSLNVQQLQALEKSLESGLGSVLKTKSQKIMDQISELERKRVQ
AtSVP        80 DQPSLELQLVENSDHARMSKEIADKSHRLRQMRGEELQGLDIEELQQLEKALETGLTRVIETKSDKIMSEISELQKKGMQ
AtAGL24      81 DPPSTHLR-LENCNLSRLSKEVEDKTKQLRKLRGEDLDGLNLEELQRLEKLLESGLSRVSEKKGECVMSQIFSLEKRGSE

TaVRT-A2    160 LAEENMRLKNQMHEVPTA-STVAVAE--AENVVPEDAHSSDSVMTAVHSGS--SQDNDDGSDISLKLALP---WK
TaVRT-B2    160 LAEENMRLKNQMHEVPTA-SMVAVADADAENVVPDDVHSSDSVMTAVHSAS--SQDNDDGSDISLKLALP---WK
HvVRT2      160 LAEENMRLKNQMHEVPTA-SMVAVAD-----VVPEDVHSSDSVMTAVHSAS--SQDNDDGSDISLKLALP---WK
OsMADS55    160 LAEENMRLRDQMPQVPT--AGLAVPD--TENVLTEDGQSSESVMTALNSGS--SQDNDDGSDISLKLGLP-----
TaSVP-A1    159 LAEENMKLRNQVGQIPTA-GKLVVAD--TENVVAEDGQSSESVMTALHSGS--SQDNDDGSDVSLKLGLPCLPWK
TaSVP-B1    159 LAEENMKLRNQVGQIPTA-GKLVVAD--TENIVAEDGQSSESVMTALHSGS--SQDNDDGSDVSLKLGLPCLPWK
HvBM10      159 LAEENKKLRNQVAQVPTA-GKLVVVD--TENVIAEDGQSSESVMTALHSGS--SQDNDDGSDVSLKLALP---WK
OsMADS22    159 LAEENMQLRNQVSQISPA-EKQVV-D--TENFVTE-GQSSESVMTALHSGSSQSQDNDDGSDVSLKLGLPCGAWK
HvBM1       159 LIEENARLKEQASKM----EMQVAAD--PLVVVYDEGQSSESVTNTSYPRP--PLDTEDSSDTSLRLGLSLFNSK
OsMADS47    160 LIEENLRLKEQVSRMSRMEEMQPGPD--S-EIVYEEGQSSESVTNASYPRP--PPDNDYSSDTSLKLGLHS----
TaSVP-A3    159 LIEENARLKEQASKM----EMQVAAD--S-PVVYEEGQSSESVTNTSYPRP--PLDTEDSSDTSLRLGLPLYNS-
TaSVP-B3    159 LIEENARLKEQASKM----EMQVAAD--S-PAVYEEGQSSESVTNTSYPRP--PLDTEDSSDTSLRLGLPLYNSK
AtSVP       160 LMDENKRLRQQGTQLTEENERLGMAES-ENAAVYEEGQSSESITNAGNSTG--APVDSESSDTSLRLGLPYGG--
AtAGL24     160 LVDENKRLRDKLETLER-----------AKLTTLKEALETESVTTNVSSYDS-GTPLEDDSDTSLKLGLPS--WE
```

**Supplemental Figure 2.** Effects of *vrt2* and *svp1* mutations on the encoded proteins.

* = stop codon. Rectangles of different colors represent exons. Red and green lines represent MADS and K box domains respectively. A red triangle indicates the position where exons are missing.

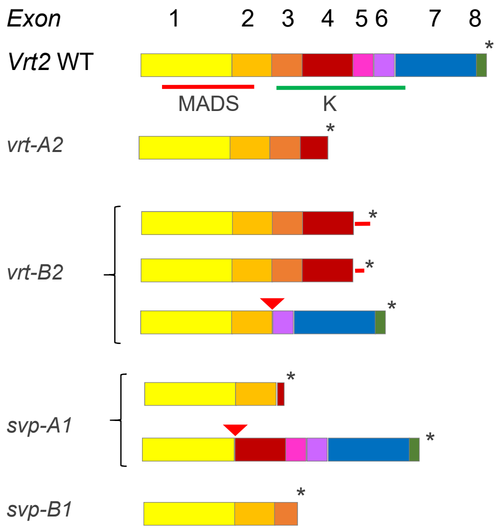

**Supplemental Figure 3** Effects of mutations in the A and B genome homeologs of *VRT2* and *SVP1*.

**(A-F)** Growth chamber experiments. The number of replications is indicated in the X axis below each genotype. **(A, C, E)** Effect of *vrt2*. **(B, D, F)** Effect of *svp1*. **(A-B)** Heading time (HT) **(C-D)** Spikelet number per spike (SNS). **(E-F)** Stem length: each node is in a different color and peduncles are in blue (-1 is the closest node to the peduncle and -4 the most basal node). **(G-I)** Field experiment for *vrt2* mutants only. **(G)** Spikelet number per spike. **(H-I)** Plant height (PH). Number of replications are below the box plots. Significance values are based on Dunnett tests compared to the WT. ns = not significant, * = $P < 0.05$, ** = $P < 0.01$, *** = $P < 0.001$. Box-plot features are explain in Statistical analyses section of Material and Methods

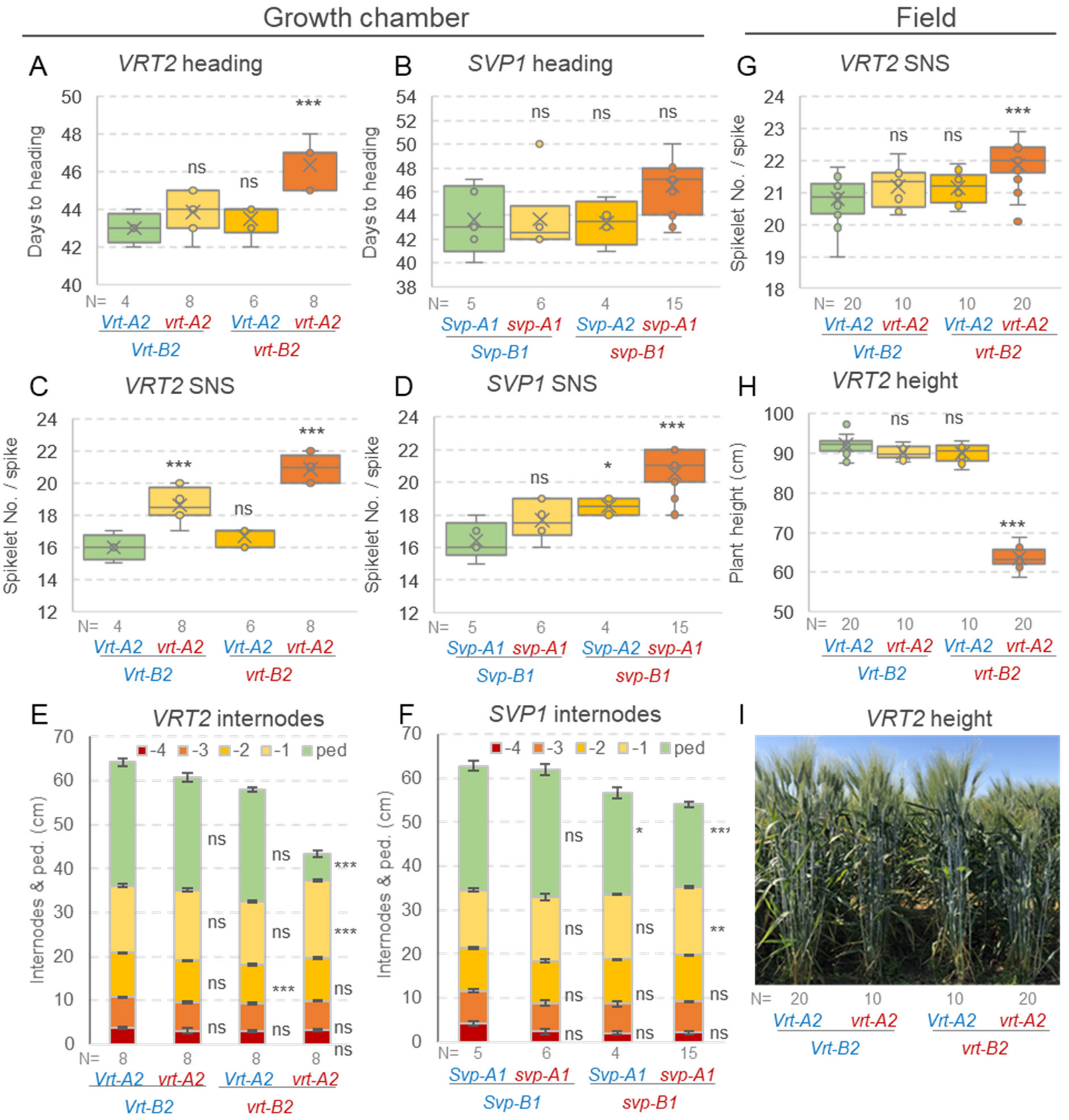

**Supplemental Figure 4** Complementation of the *vrt2* mutation by weak constitutive transgenic *UBI::VRT2* line T#8.

**(A)** Heading time. **(B)** Peduncle length. **(C)** Spikelet number per spike (SNS). WT= homozygous wild type *VRT2* alleles for both homeologs. *vrt2* = homozygous loss-of-function *vrt-A2* and *vrt-B2* alleles. T= *UBI::VRT2* transgene present. NT = Transgene absent. N= 12 biological replicates per genotype (except WT NT where N = 10). Different letters indicate significant differences using a Tukey test $P < 0.05$. Box-plot features are explained in the Statistical analyses section of Material and Methods.

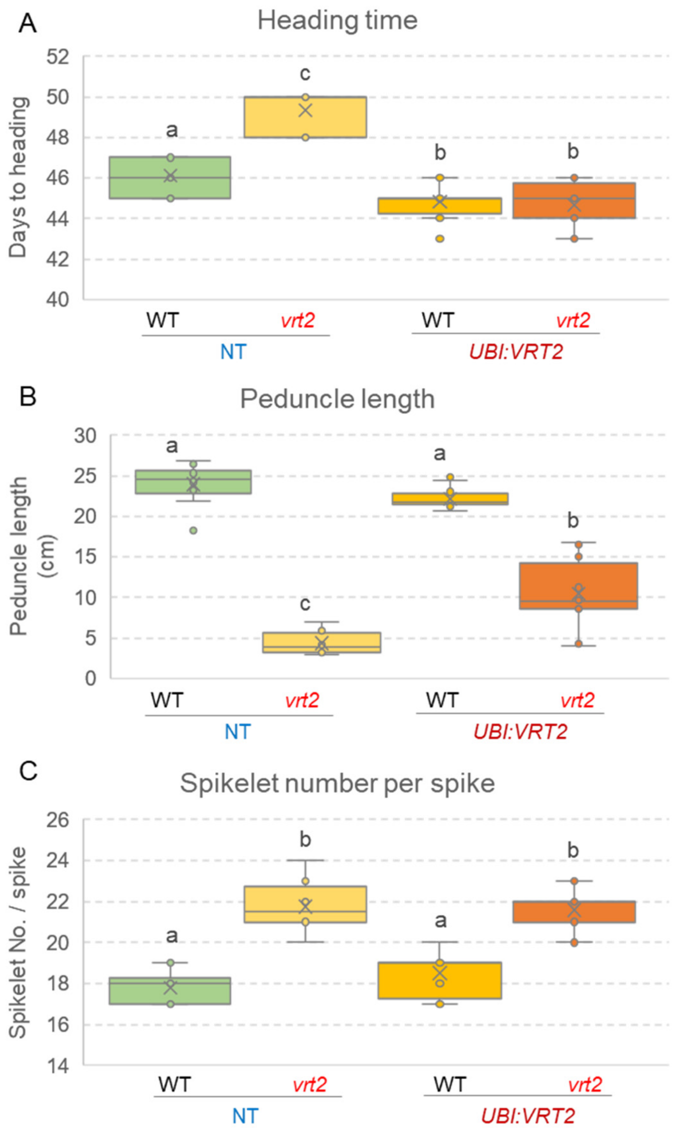

**Supplemental Figure 5.** Phenotypic comparison between *vrn1 ful2* and *vrt2 vrn1 ful2* mutants (in a *vrn2* background).

**(A-B)** Plants were phenotyped at 110 days. **(A** and **C)** *vrn1 ful2* **(B** and **D-G)** *vrt2 vrn1 ful2*. **(C-D)** "Spike of tillers" with lateral spikelets replaced by vegetative tillers. **(D)** "Spike of tillers" emerged from most shoots in *vrn1 ful2* but only from 16% of the shoots in *vrt2 vrn1 ful2*. **(E)** Tiller without an emerging spike. **(F-G)** Dissection of tillers without emerging spikes. **(F)**. Note the very short stem carrying an undeveloped spike (~ 4 mm) with relatively large bracts (~5 mm) subtending the basal lateral meristems. **(G)** Meristems from the non-emerging spikes eventually died inside the sheaths.

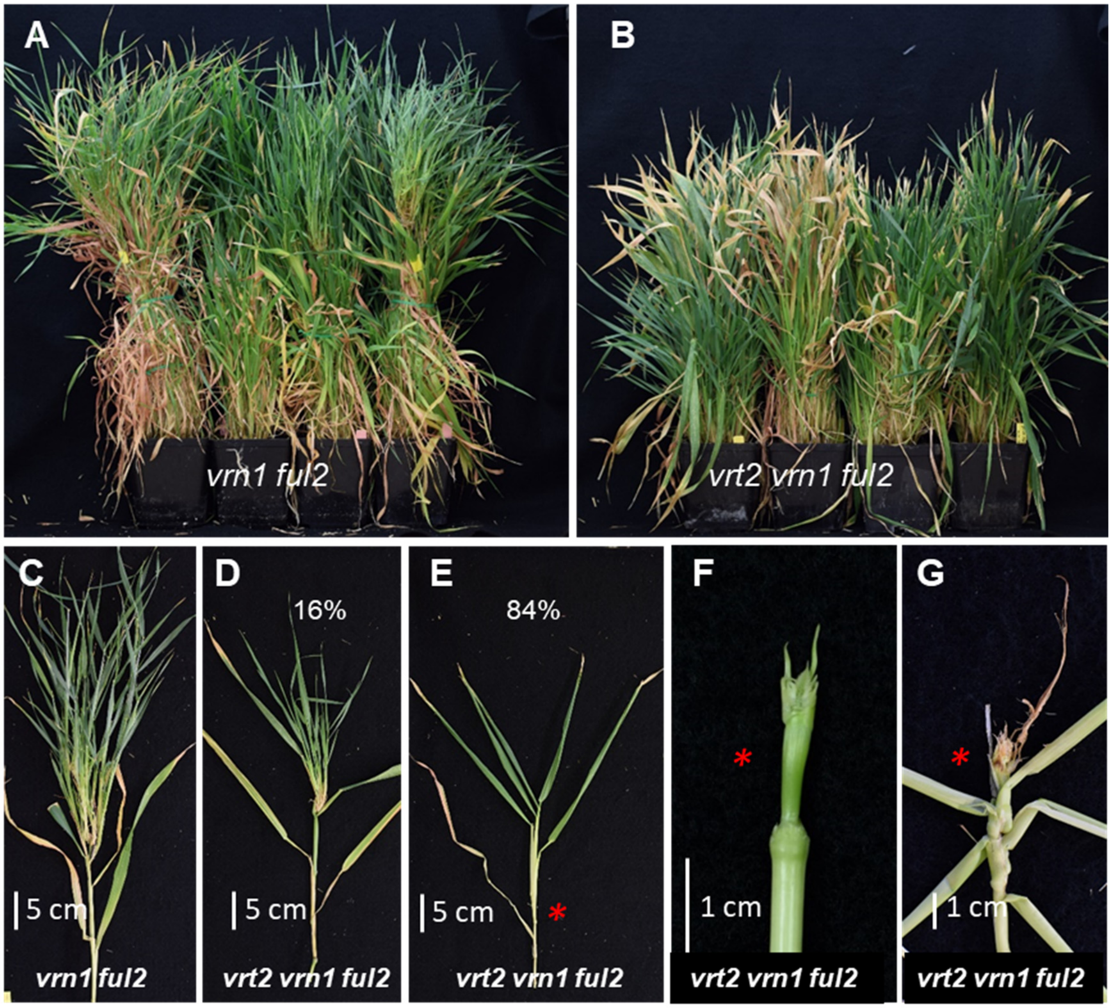

**Supplemental Figure 6.** Effect of the *vrt2* mutation in the partial mutant *Vrn1 ful2* (the underline indicates heterozygous *Vrn-A1 vrn-A1* and homozygous *vrn-B1 vrn-B1*).

**(A)** Days to heading. (**B**) Stem length excluding heads. (**C**) Spikelet number per spike. (**D**) Glume length. (**E**) Lemma length. (**F**) Percent of branched spikelets with florets regressed to spikelets (N = number of plants in A-C & F, spikelets in D and E). ** = $P < 0.01$, *** = $P < 0.001$ (*t* test). Box-plot features are explained in the Statistical analyses section of Material and Methods.

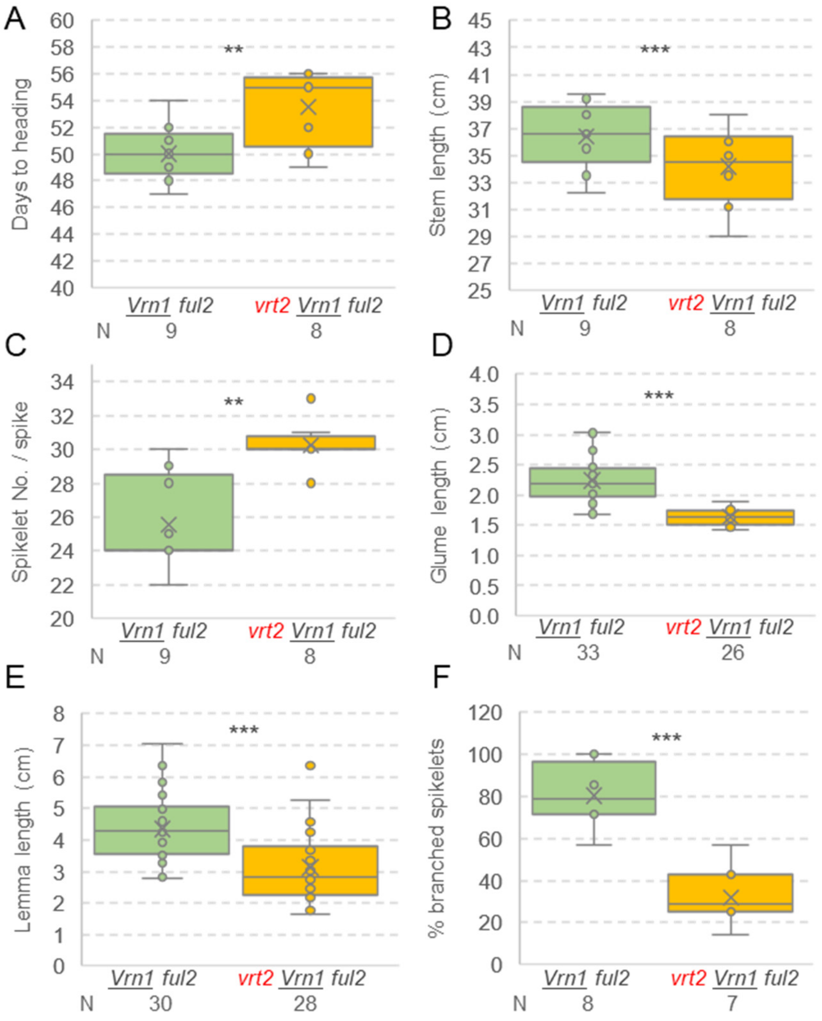

**Supplemental Figure 7.** Auto-activation tests for the bait and prey vectors used in yeast two-hybrid assays (only bait for SEP proteins).

Vector pGBKT7 expressing the GAL4 DNA binding domain is the empty vector used to generate all the bait vectors, whereas pGADT7 expressing the GAL4 activation domain is the empty vector used to generate all the preys. SD medium lacking Leucine and Tryptophan (-L-W) was used to select for yeast transformants containing both bait and prey vectors. We tested the interactions on SD media lacking Leucine, Tryptophan, Histidine and Adenine (-L-W-H-A).

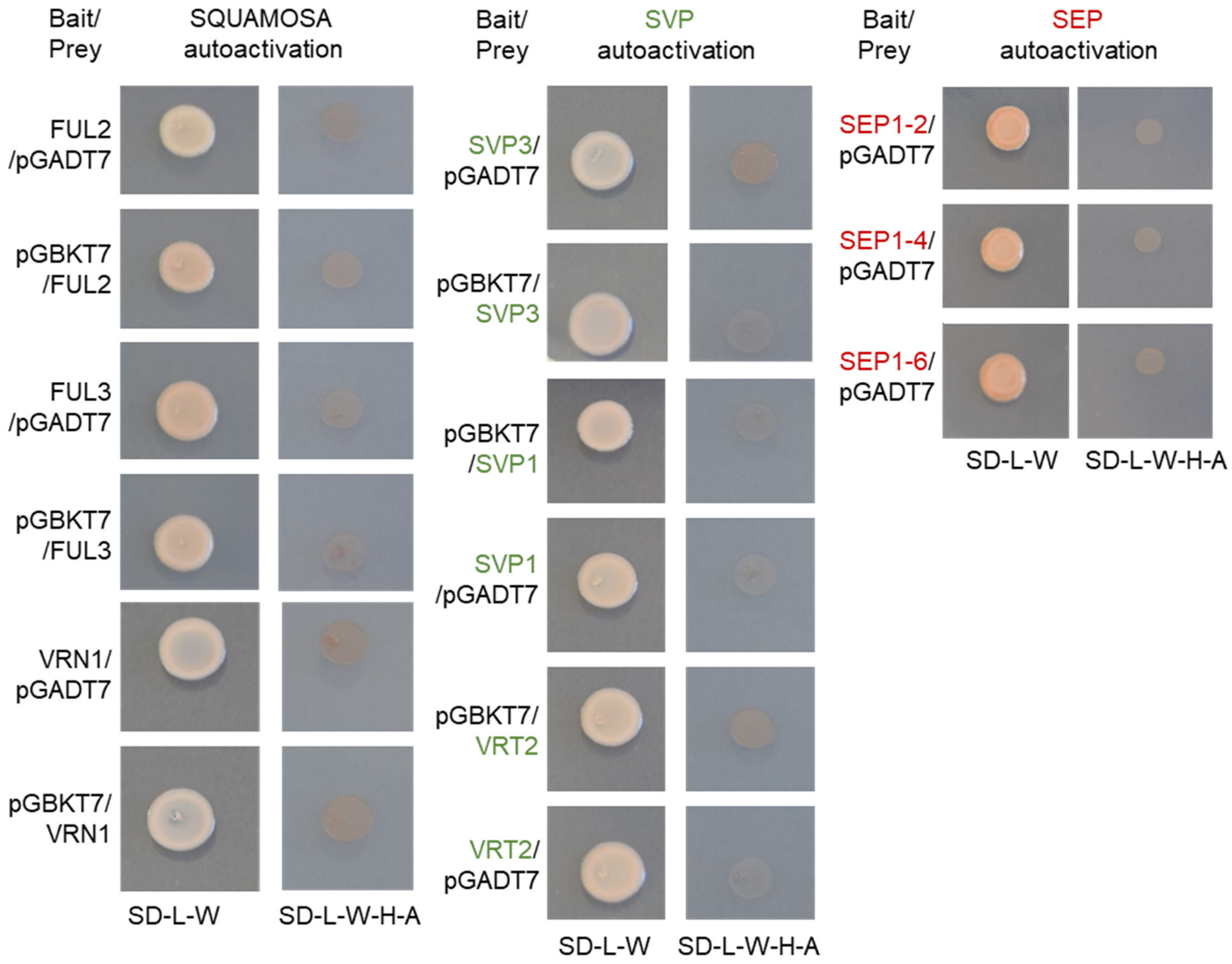

**Supplemental Figure 8.** Yeast-two-hybrid (Y2H) interactions among proteins within the SQUAMOSA- and SVP-clades.

We tested the interactions among the SQUAMOSA MADS-box proteins VRN1, FUL2 and FUL3, and among the SVP proteins VRT2, SVP1 and SVP3. We did not test the interactions among the SEP proteins SEP1-2, SEP1-4, and SEP1-6.

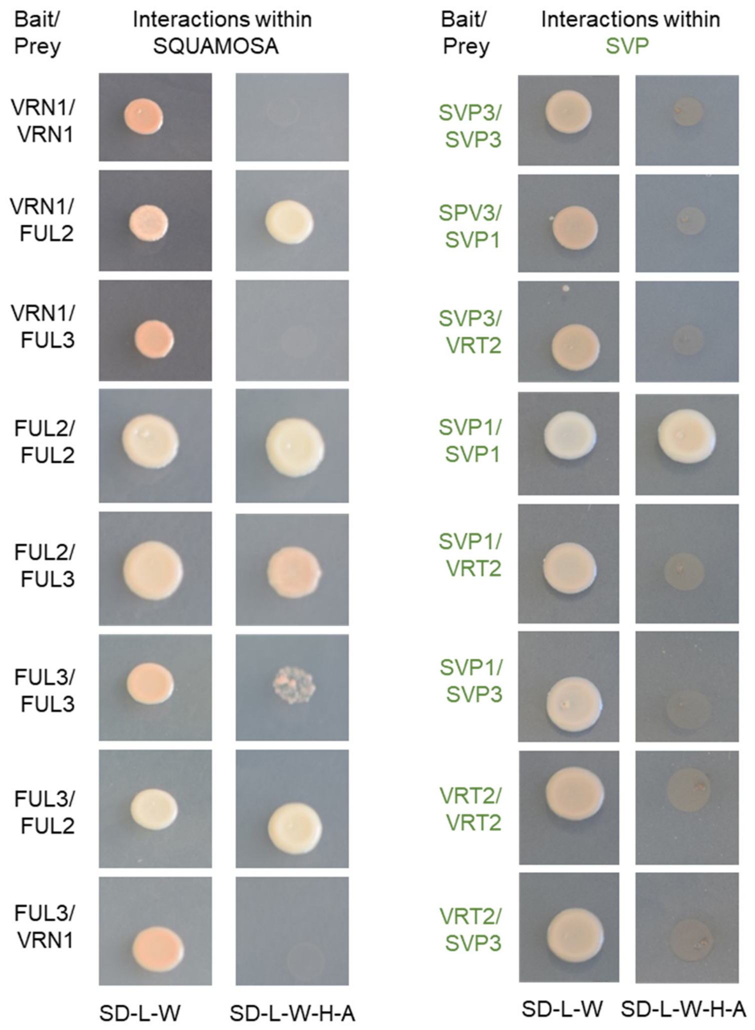

**Supplemental Figure 9.** Yeast-two-hybrid (Y2H) interactions between wheat MADS-box proteins of the SQUAMOSA (VRN1, FUL2 and FUL3), SVP (VRT2, SVP1 and SVP3) and SEP (SEP1-2, SEP1-4, SEP1-6) classes.

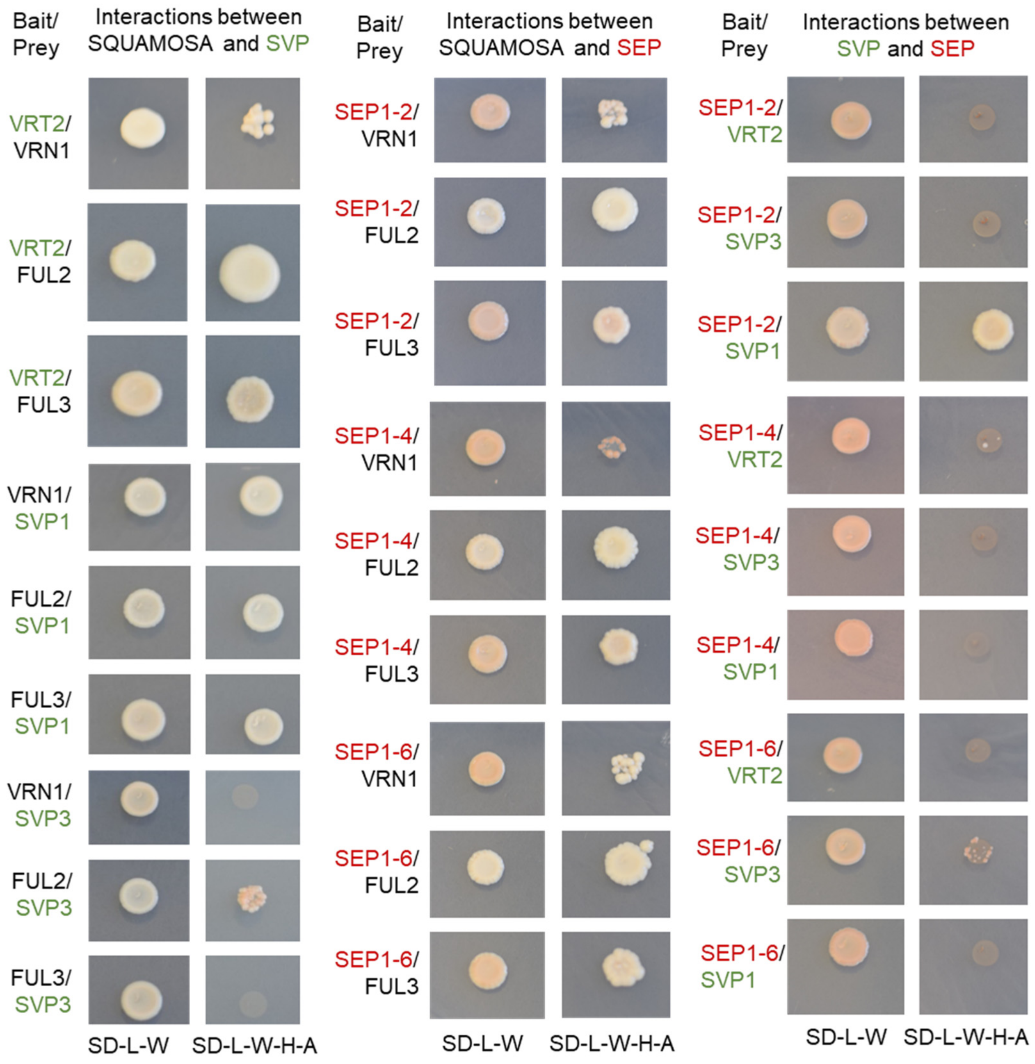

**Supplemental Figure 10.** Bimolecular Fluorescence Complementation (BiFC) between proteins of the SQUAMOSA-clade (VRN1, FUL2 and FUL3) and proteins of both the SVP-clade (VRT2 and SVP1) and SEPALLATA-clade (SEP1-2, SEP1-4 and SEP1-6) in wheat protoplasts. The bar is 200 µm. **(A-I)** Positive nuclear signal. **(J-O)** No nuclear signal. **(P-W)** Negative control.

**A)** UBI::N-YFP:**VRT2** – UBI::C-YFP:**FUL3** (nuclear signal plus fluorescent aggregates)

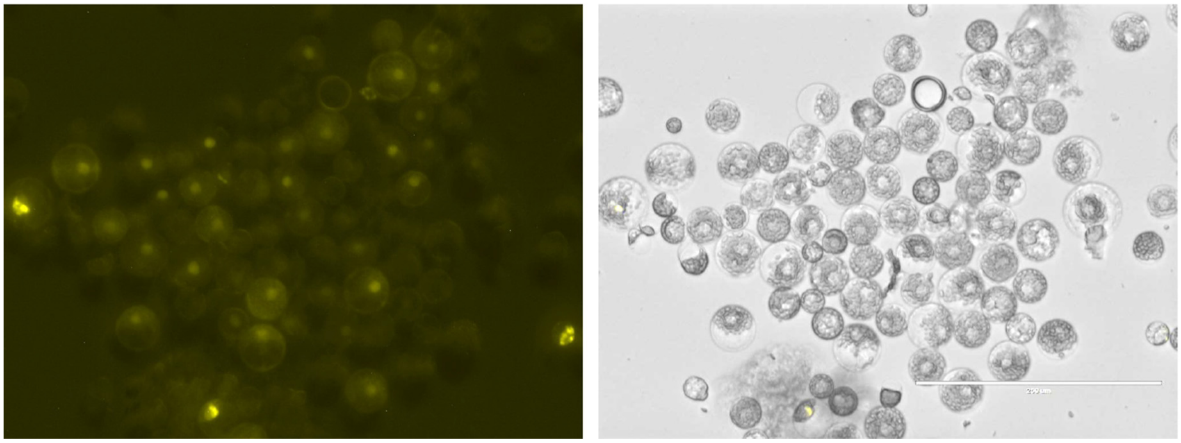

**B)** UBI::N-YFP:**SVP1** – UBI::C-YFP:**FUL3** (nuclear signal plus fluorescent aggregates)

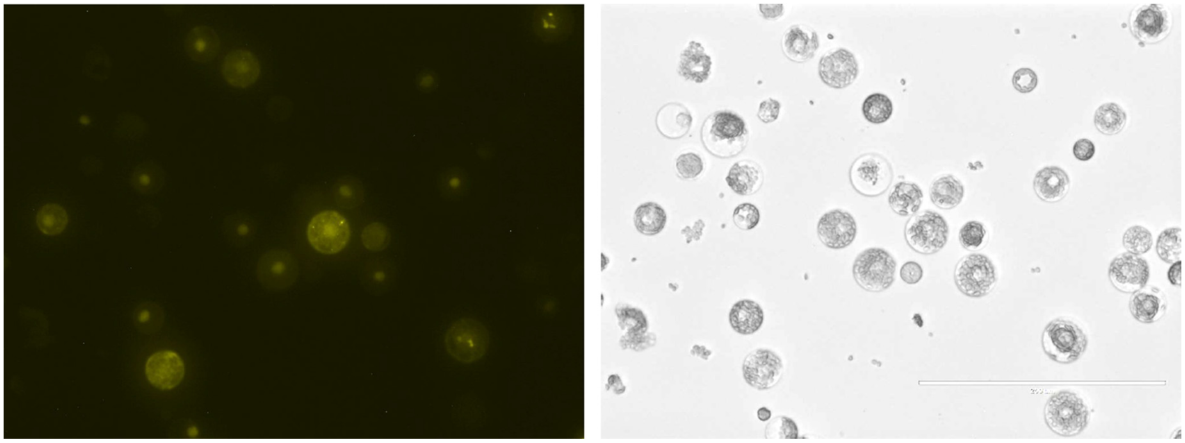

**C)** UBI::N-YFP:**SEP1-2 -** UBI::C-YFP:**FUL3** (nuclear signal plus fluorescent aggregates)

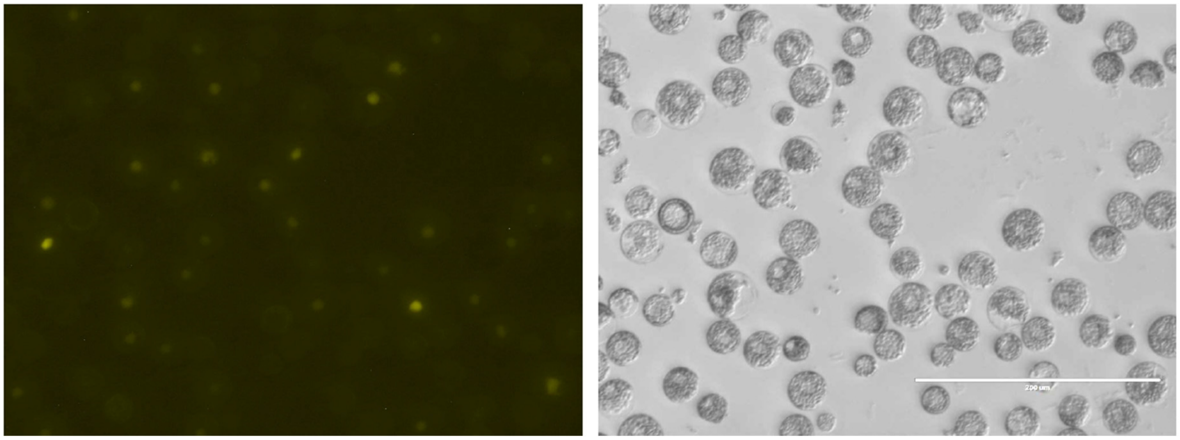

**D)** UBI::N-YFP:**VRT2** – UBI::C-YFP:**VRN1** (nuclear signal plus fluorescent aggregates)

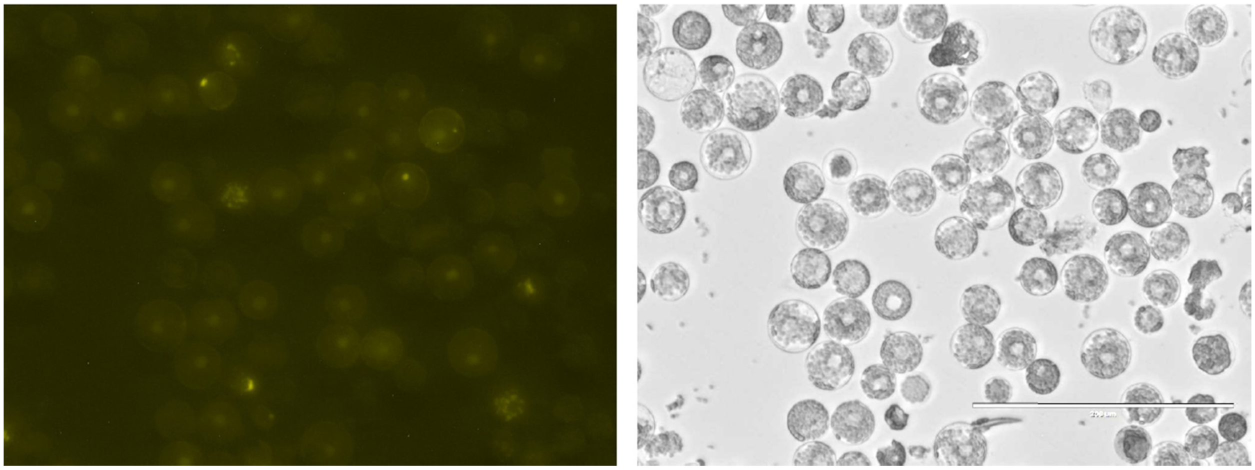

**E)** UBI::N-YFP:**SVP1** – UBI::C-YFP:**VRN1** (nuclear signal plus fluorescent aggregates)

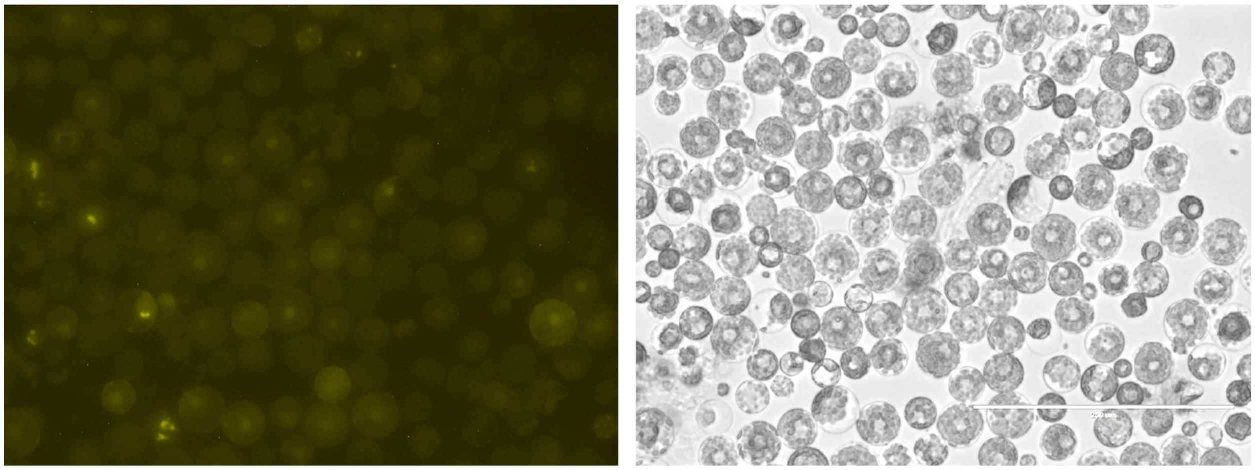

**F)** UBI::N-YFP:**SEP1-2** – UBI::C-YFP:**VRN1** (nuclear signal)

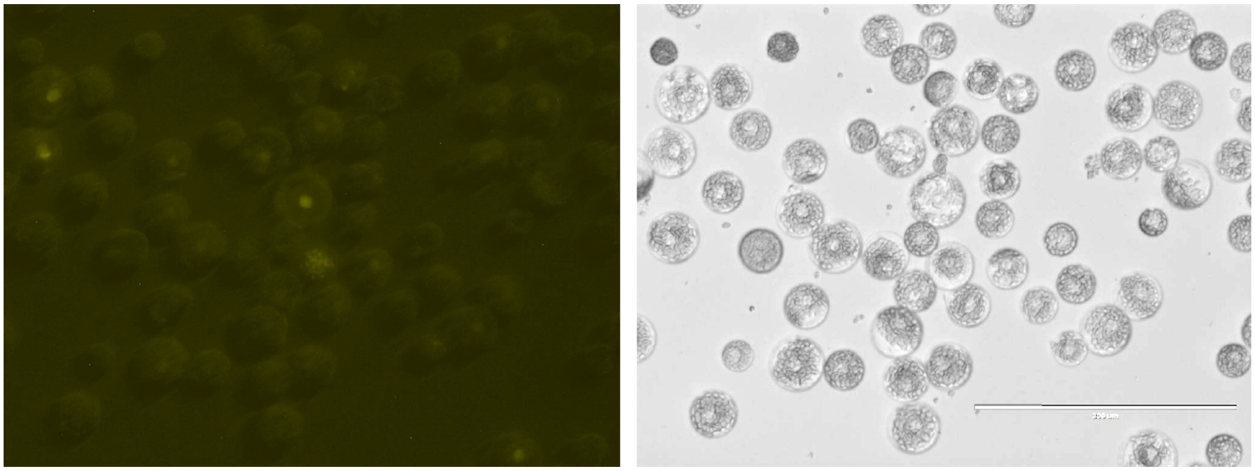

**G)** UBI::N-YFP:**VRT2** – UBI::C-YFP:**FUL2** (nuclear signal plus fluorescent aggregates[1])

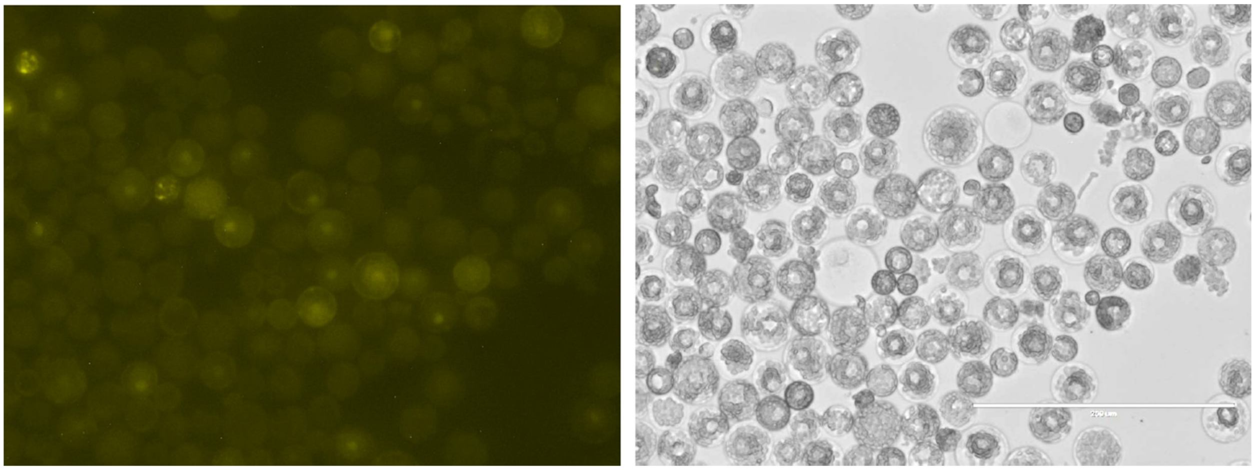

UBI::N-YFP:**VRT2** – UBI::C-YFP:**FUL2** fluorescent aggregates outside nucleus[1].

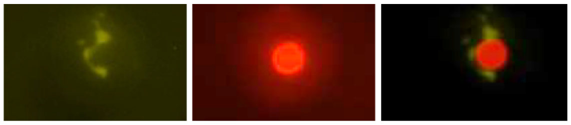

FDL2 mCherry as nuclear marker.

**H)** UBI::N-YFP:**SVP1 –** UBI::C-YFP:**FUL2** (nuclear signal indicated by arrows & fluorescent aggregates[1])

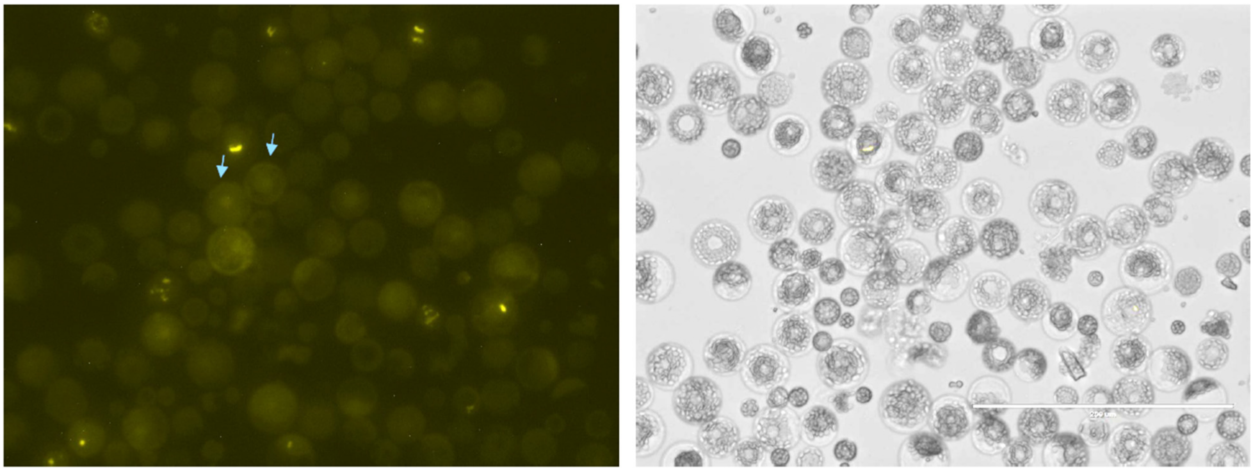

UBI::N-YFP:**SVP1** – UBI::C-YFP:**FUL2** fluorescent aggregates outside the nucleus[1].

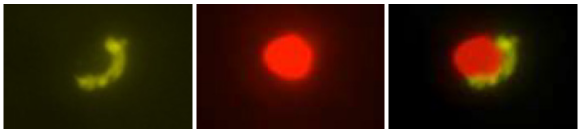

FDL2 mCherry as nuclear marker.

[1] We currently do not know if these bright aggregates are the result of the stabilization of the MADS-box complexes by the YFP reassembly (Robida and Kerppola, 2009, J. Mol. Biol. 394: 391-409), a technical artifact of the over-expression, or a reflection of the real distribution of these MADS-box complexes in the wheat protoplasts.

**I)** UBI::N-YFP:**SEP1-2** – UBI::C-YFP:**FUL2**
Nuclear signal indicated by arrows plus fluorescent aggregates.

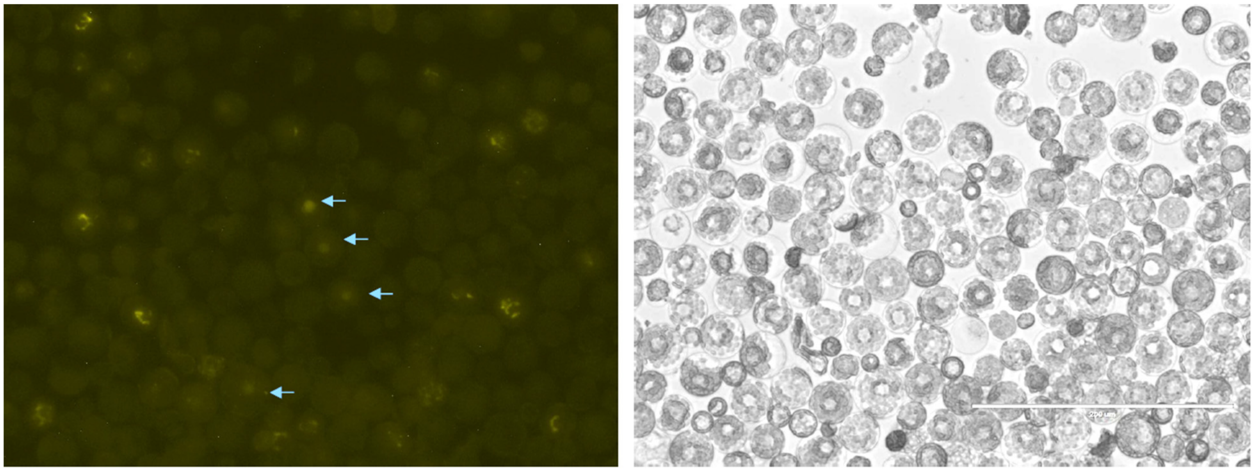

**J)** UBI::N-YFP:**SEP1-4** – UBI::C-YFP:**FUL2** (only fluorescent aggregates)

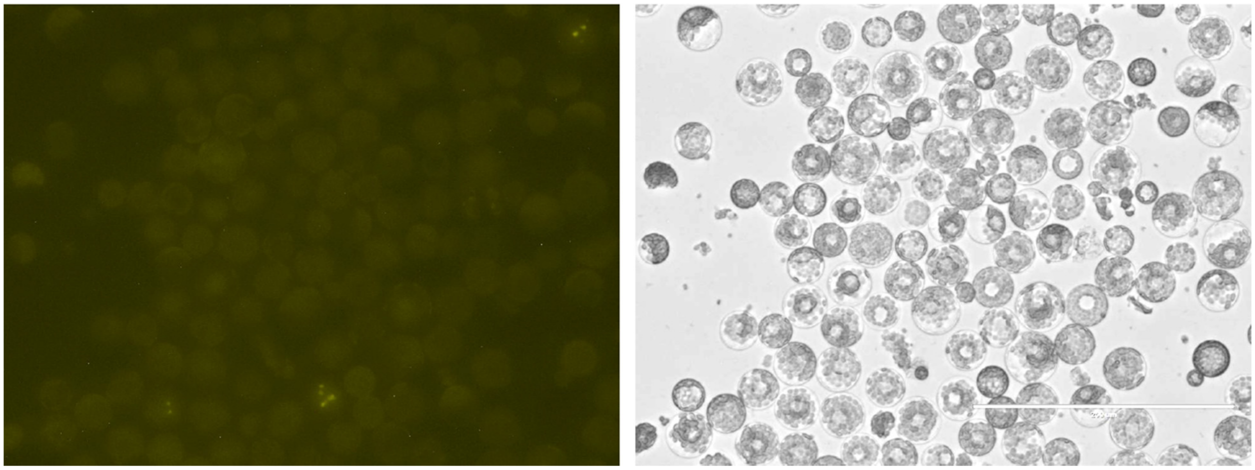

**K)** UBI::N-YFP:**SEP1-6** – UBI::C-YFP:**FUL2** (only fluorescent aggregates)

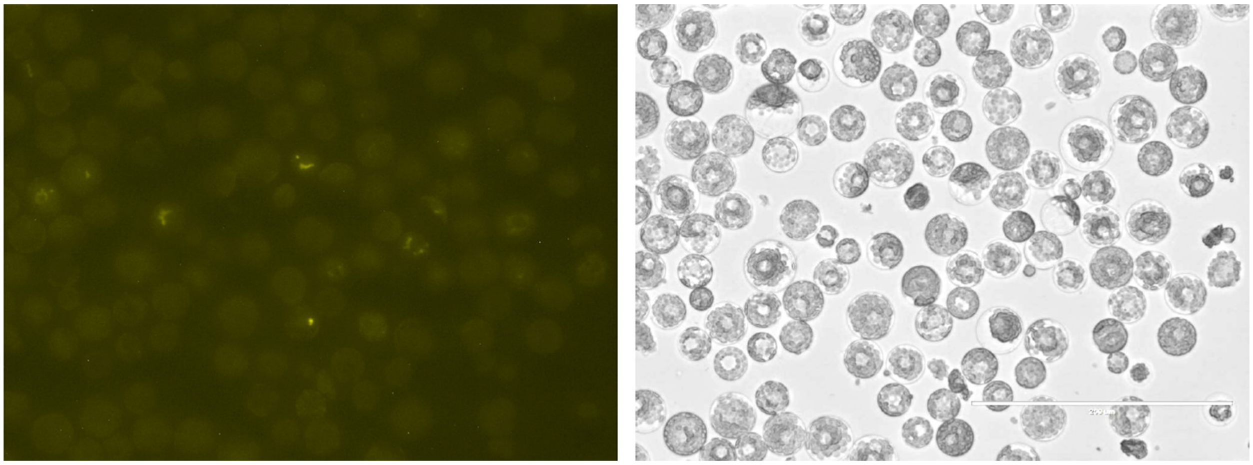

**L)** UBI::N-YFP:**SEP1-6** – UBI::C-YFP:**VRN1** (only fluorescent aggregates)

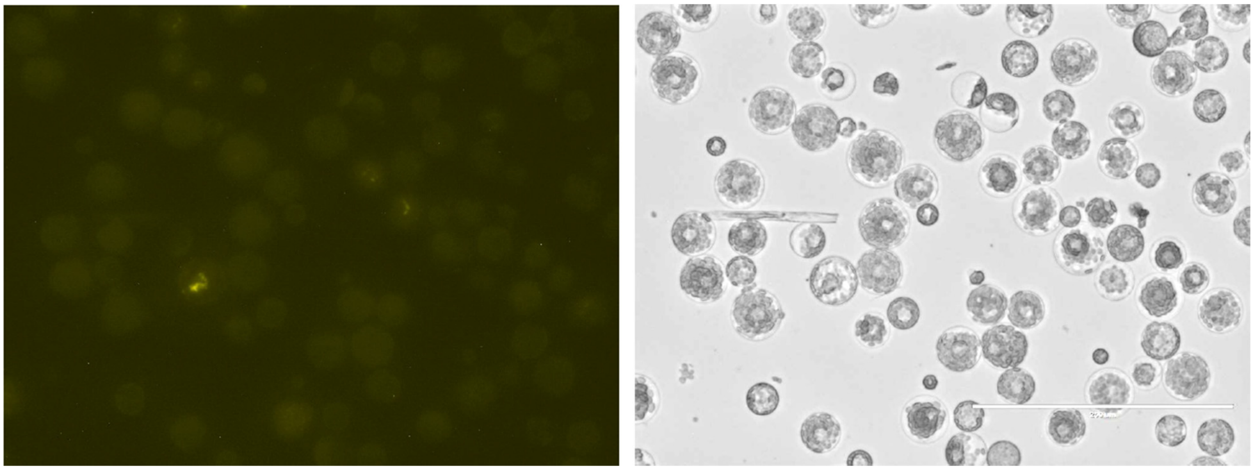

**M)** UBI::N-YFP:**SEP1-4** – UBI::C-YFP:**VRN1** (no signal)

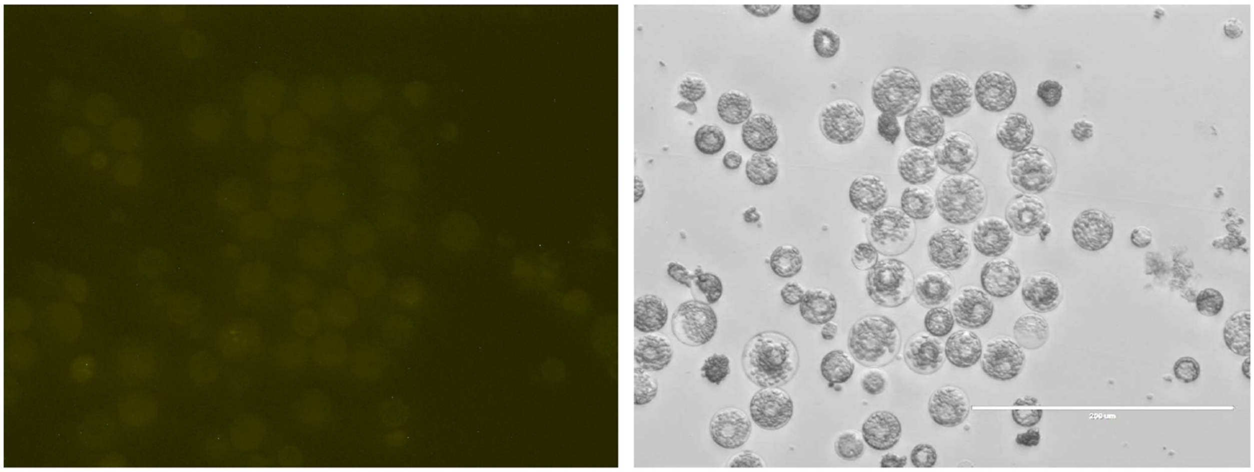

**N)** UBI::N-YFP:**SEP1-4** – UBI::C-YFP:**FUL3** (no signal)

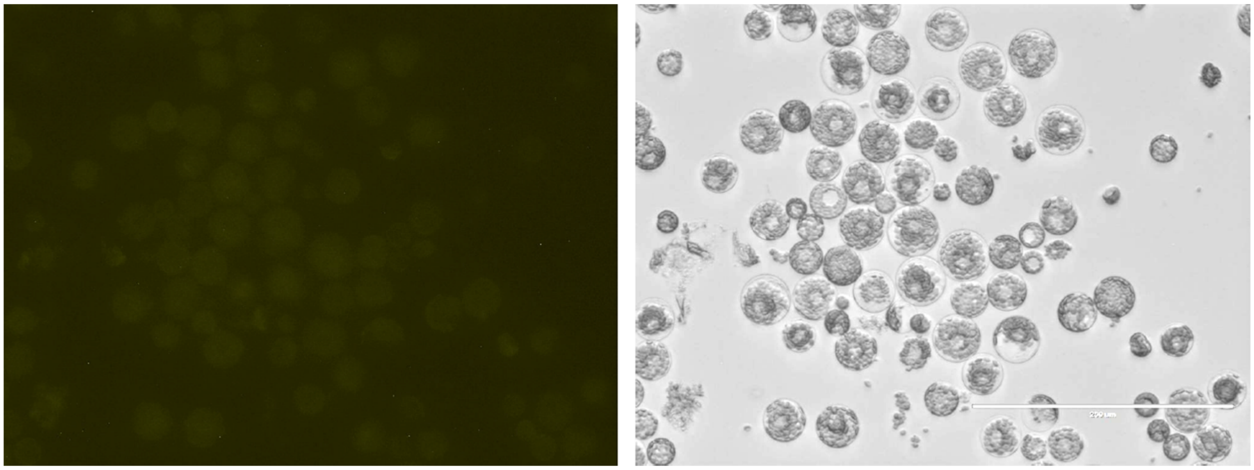

**O)** UBI::N-YFP:**SEP1-6** – UBI::C-YFP:**FUL3** (no signal)

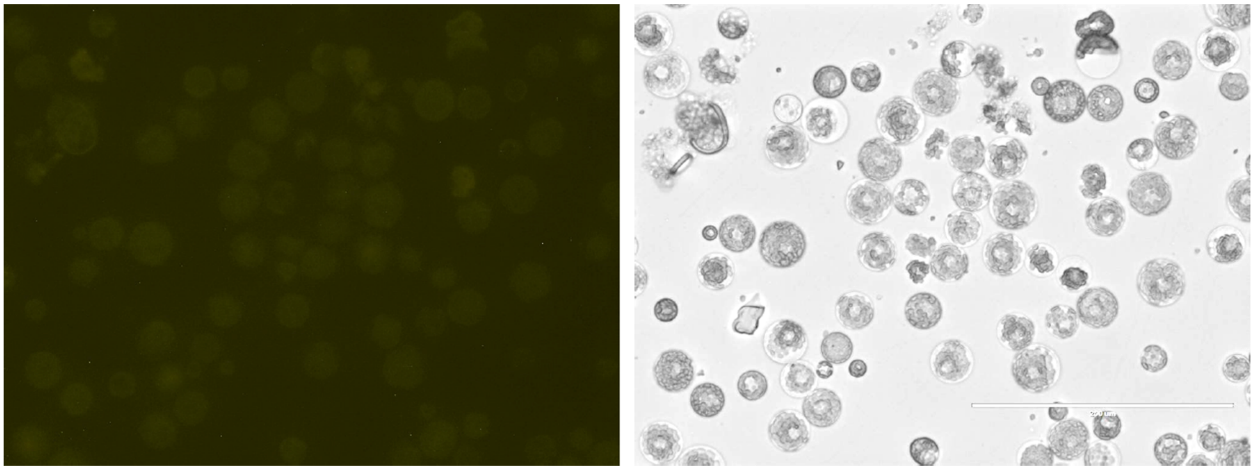

**BiFC negative controls**

**P)** UBI::N-YFP:**VRT2** – C-YFP (negative control, no signal)

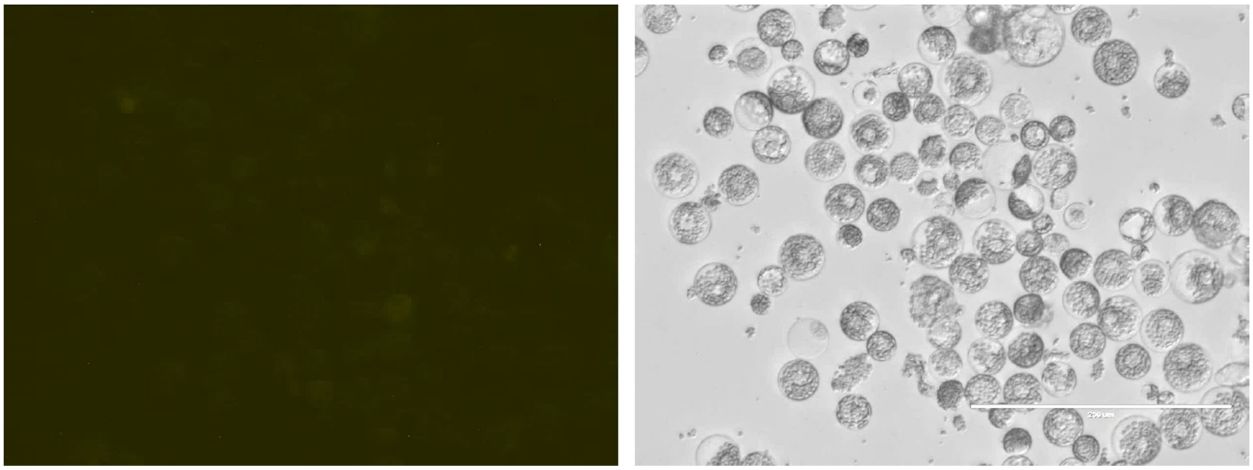

**Q)** UBI::N-YFP:**SVP1** – C-YFP (negative control, no signal)

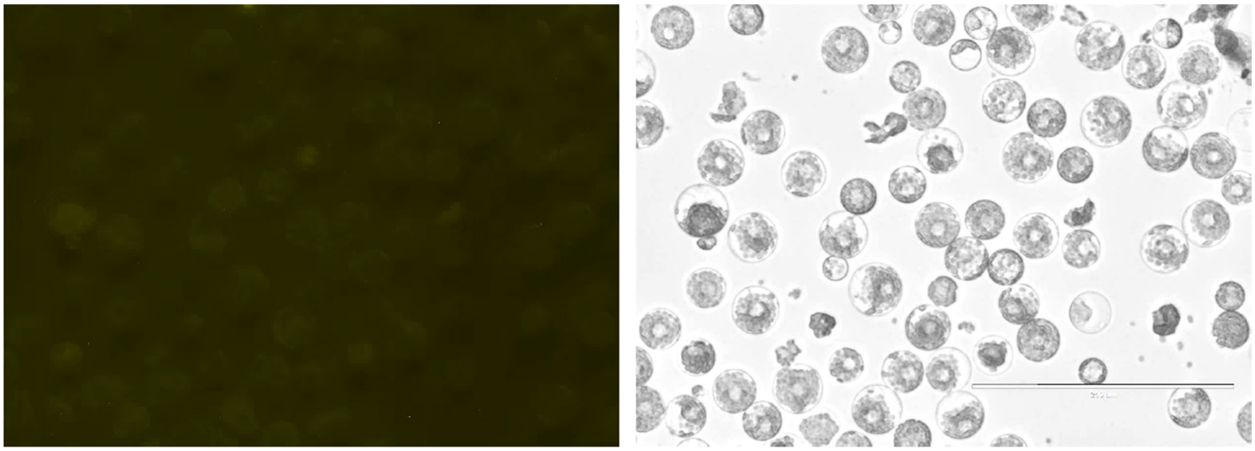

**R)** UBI::N-YFP:**FUL3**- C-YFP (negative control, no signal)

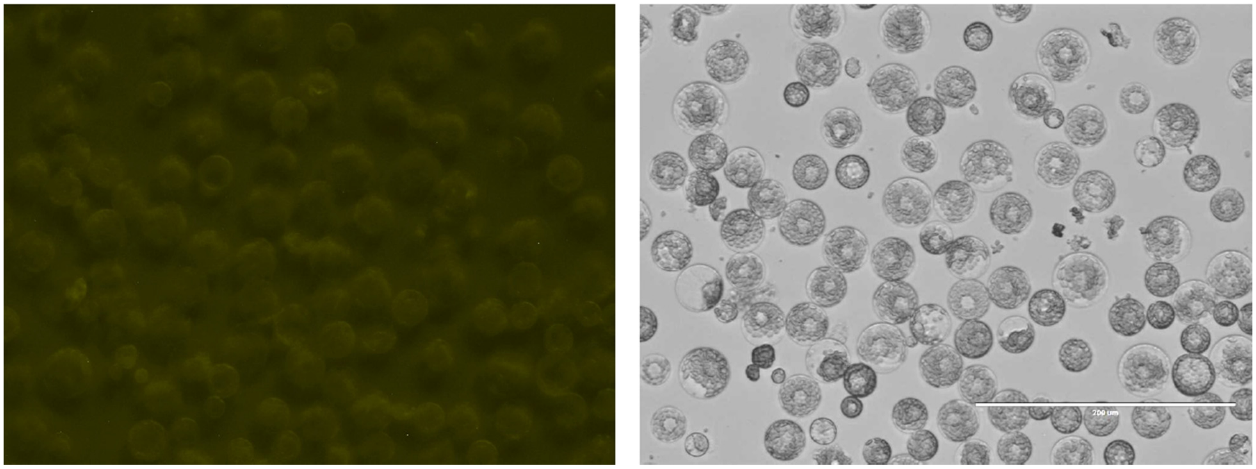

**S)** UBI::N-YFP:**FUL2** – C-YFP (negative control, no signal)

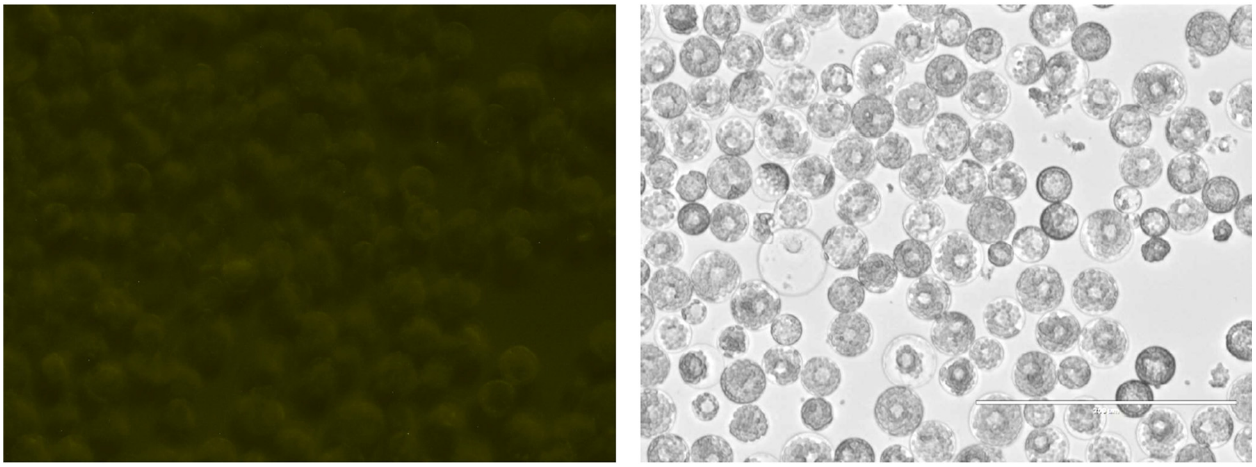

**T)** UBI::N-YFP:**VRN1** – C-YFP (negative control, no signal)

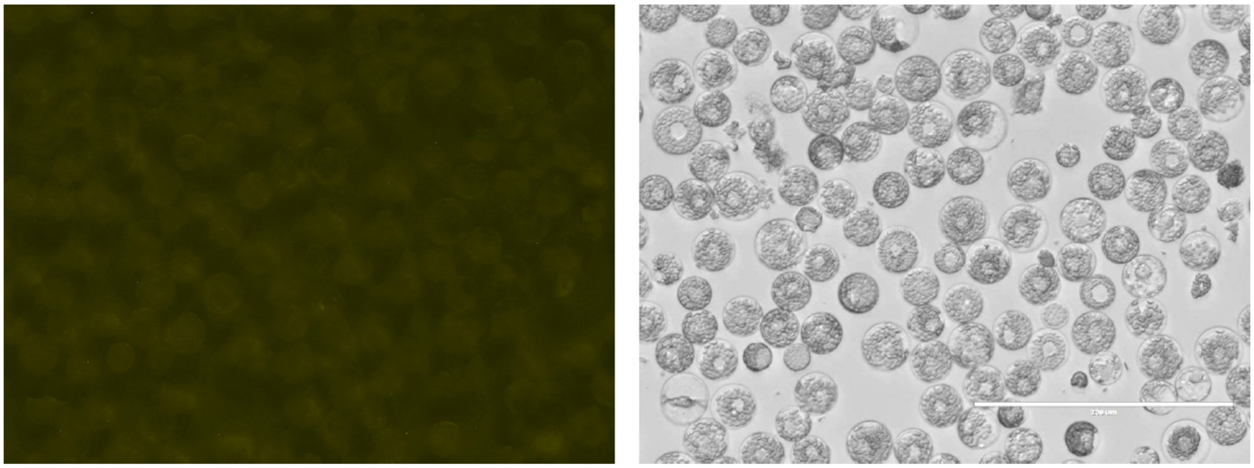

**U)** UBI::N-YFP:**SEP1-2** – C-YFP (negative control, no signal)

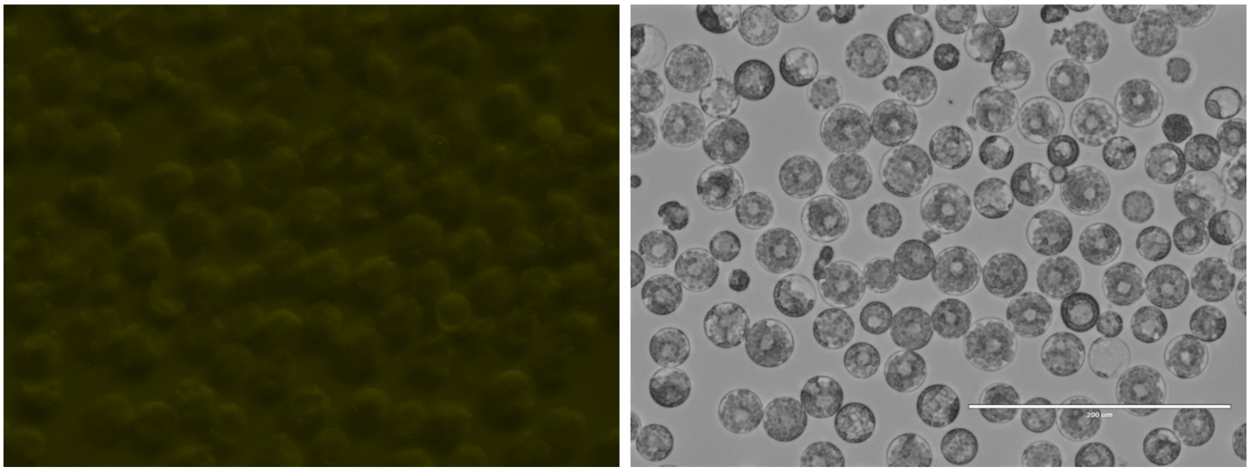

**V)** UBI::N-YFP:**SEP1-4** – C-YFP (negative control, no signal)

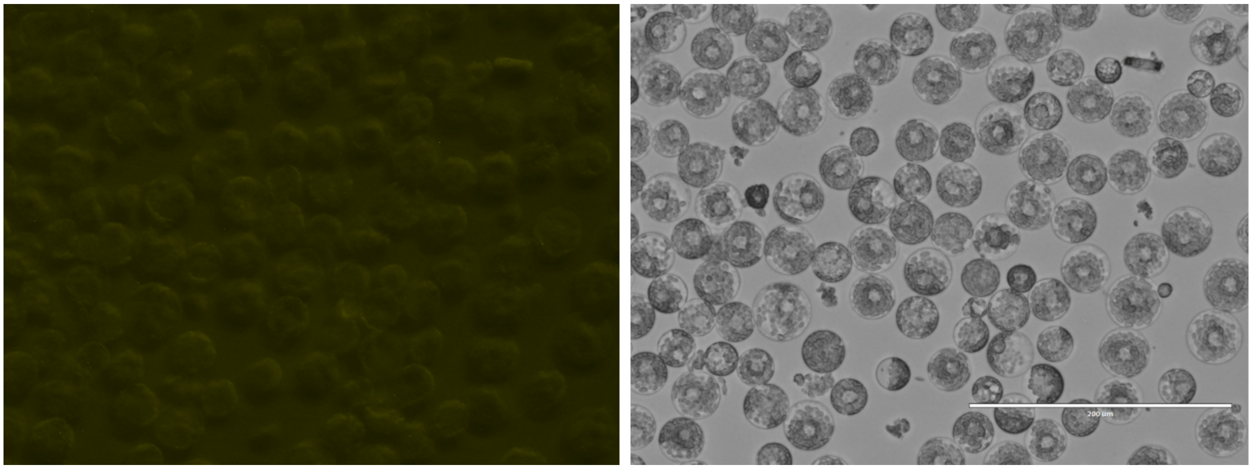

**W)** UBI::N-YFP:**SEP1-6** – C-YFP (negative control, no signal)

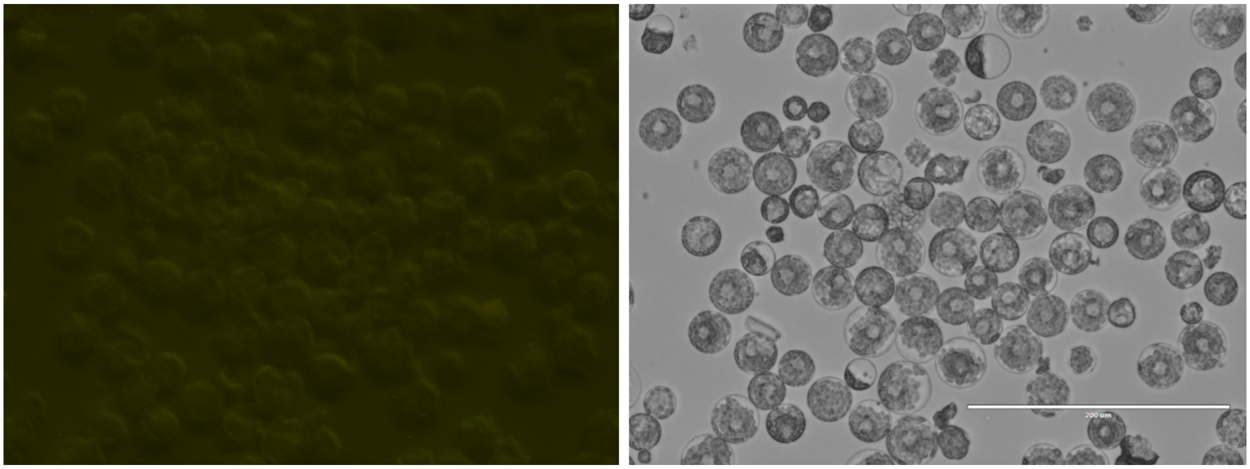
