## Supplemental Tables for "Interactions Between SQUAMOSA and SVP MADS-box Proteins Regulate Meristem Transitions During Wheat Spike Development"

### SUPPLEMENTAL INFORMATION

#### SUPPLEMENTAL TABLES

**Supplemental Table 1.** Nomenclature, accession numbers, synonyms of wheat genes used in this study and their rice orthologs. We adopted the wheat MADS-box nomenclature proposed in a recent and exhaustive review (1) except for the *VRN1* (2) and *VRT2* (3) genes, for which the historical names were used, and *FUL2* and *FUL3*, for which a comprehensive phylogenetic analysis and nomenclature has been proposed before (4).

| MADS | | Wheat | RefSeq v1.1 | | | | |
|---|---|---|---|---|---|---|---|
| Class | Rice | A genome | B genome | **Wheat** | Ref. | Synonyms | Ref. |
| **SVP** | *OsMADS22* | *TraesCS6A02G313800.1* | *TraesCS6B02G343900.1* | *SVP1* | (1) | *BM10* | (5) |
| | *OsMADS47* | *TraesCS4A02G002600.1* | *TraesCS4B02G302600.1* | *SVP3* | (1) | *BM1* | (5) |
| | *OsMADS55* | *TraesCS7A02G175200.1* | *TraesCS7B02G080300.1* | *VRT2* | (3) | *TaSVP-2* | (1) |
| **A-class** | *OsMADS14* | *TraesCS5A02G391700.1* | *TraesCS5B02G396600.1* | *VRN1* | (2) | *TaMADS11*<br>*BM5*<br>*TaFUL1*<br>*TaAP1-1* | (6)<br>(5)<br>(7)<br>(1) |
| | *OsMADS15* | *TraesCS2A02G261200.1* | *TraesCS2B02G281000.1* | *FUL2* | (7) | *BM8*<br>*TaAP1-3* | (5)<br>(1) |
| | *OsMADS18* | *TraesCS2A02G174300.1* | *TraesCS2B02G200800.1* | *FUL3* | (7) | *BM3*<br>*TaAP1-2* | (5)<br>(1) |
| **B-class** | *OsMADS4* | *TraesCS1A02G264300.1* | *TraesCS1B02G275000.1* | *PI1* | (1) | *WPI-1* | (8) |
| | *OsMADS16* | *TraesCS7A02G383800.1* | *TraesCS7B02G286600.1* | *AP3-1* | (1) | *TaMADS51*<br>*WAP3* | (6)<br>(8) |
| **C-class** | *OsMADS3* | *TraesCS3A02G314300.1* | *TraesCS3B02G157500.1* | *AG2* | (1) | *WAG-2* | (9) |
| | *OsMADS58* | *TraesCS1A02G125800.1* | *TraesCS1B02G144800.1* | *AG1* | (1) | *WAG-1* | (9) |
| **E-class** | *OsMADS1* | (*TraesCS4A02G058900.1*)[a] | *TraesCS4B02G245700.1* | *SEP1-2* | (1) | *BM7*<br>*WLHS1*<br>*TaSEP2* | (5)<br>(10)<br>(11) |
| | *OsMADS34* | *TraesCS5A02G391800.1* | *TraesCS5B02G396700.1* | *SEP1-6* | (1) | *TaAGLG1*<br>*PAP2*<br>*TaSEP5* | (2)<br>(12)<br>(11) |
| | *OsMADS5* | *TraesCS7A02G122000.1* | *TraesCS7B02G020800.1* | *SEP1-4* | (1) | *TaSEP6* | (11) |
| | *OsMADS7* | *TraesCS7A02G260600.1* | *TraesCS7B02G158600.1* | *SEP3-1* | (1) | *WSEP*<br>*TaSEP4* | (10)<br>(11) |
| | *OsMADS8* | *TraesCS5A02G286800.1* | *TraesCS5B02G286100.1* | *SEP3-2* | (1) | *TaSEP3* | (11) |

[a] Pseudogene in Chinese Spring but complete and expressed gene in Kronos

#### References Supplemental Table 1

**Supplemental Table 2.** Summary statistics for the Quant-Seq samples. 100 bp not-paired reads were obtained from Hi-Seq.

| Genotype | Stage | Rep. | Raw Seq. | Post filtering & trimming Seq. | Avg. read length | % GC | Avg. quality per read |
|---|---|---|---|---|---|---|---|
| *vrn1 ful2* | Vegetative | 1 | 8,908,554 | 8,133,600 | 74.6 | 44 | 36.4 |
| *vrn1 ful2* | Vegetative | 2 | 8,712,913 | 7,941,270 | 73.9 | 44 | 36.5 |
| *vrn1 ful2* | Vegetative | 3 | 8,058,165 | 7,421,715 | 74.4 | 44 | 36.4 |
| *vrn1 ful2* | Vegetative | 4 | 6,645,853 | 6,095,630 | 74.4 | 43 | 36.3 |
| *vrn1 ful2* | Double Ridge | 1 | 7,410,731 | 6,806,199 | 74.5 | 44 | 36.5 |
| *vrn1 ful2* | Double Ridge | 2 | 7,917,524 | 7,322,349 | 74.7 | 44 | 36.5 |
| *vrn1 ful2* | Double Ridge | 3 | 7,107,594 | 6,531,596 | 74.3 | 44 | 36.4 |
| *vrn1 ful2* | Double Ridge | 4 | 8,690,383 | 8,001,221 | 74.4 | 44 | 36.5 |
| *vrn1 ful2* | Post Double Ridge | 1 | 7,815,528 | 7,129,083 | 74.3 | 44 | 36.5 |
| *vrn1 ful2* | Post Double Ridge | 2 | 8,229,842 | 7,487,536 | 73.8 | 45 | 36.5 |
| *vrn1 ful2* | Post Double Ridge | 3 | 7,924,170 | 7,292,702 | 74.6 | 44 | 36.4 |
| *vrn1 ful2* | Post Double Ridge | 4 | 7,434,355 | 6,776,312 | 74.0 | 44 | 36.6 |
| *vrn1 ful2* | Terminal spikelet | 1 | 7,914,775 | 7,224,431 | 73.8 | 45 | 36.2 |
| *vrn1 ful2* | Terminal spikelet | 2 | 9,227,663 | 8,530,373 | 74.7 | 44 | 36.5 |
| *vrn1 ful2* | Terminal spikelet | 3 | 7,132,618 | 6,540,557 | 74.2 | 44 | 36.4 |
| *vrn1 ful2* | Terminal spikelet | 4 | 8,463,345 | 7,787,485 | 74.5 | 44 | 36.5 |
| *vrn1* | Vegetative | 1 | 8,116,199 | 7,382,112 | 74.0 | 44 | 36.5 |
| *vrn1* | Vegetative | 2 | 8,248,815 | 7,526,889 | 74.4 | 44 | 36.4 |
| *vrn1* | Vegetative | 3 | 8,678,594 | 7,838,265 | 73.7 | 44 | 36.4 |
| *vrn1* | Vegetative | 4 | 7,024,474 | 6,368,904 | 73.6 | 43 | 36.7 |
| *vrn1* | Double Ridge | 1 | 7,629,003 | 6,983,084 | 74.3 | 44 | 36.4 |
| *vrn1* | Double Ridge | 2 | 6,081,080 | 5,525,915 | 73.3 | 44 | 36.1 |
| *vrn1* | Double Ridge | 3 | 7,770,006 | 7,209,184 | 74.7 | 44 | 36.5 |
| *vrn1* | Double Ridge | 4 | 7,224,717 | 6,630,339 | 74.4 | 44 | 36.5 |
| *vrn1* | Post Double Ridge | 1 | 6,348,081 | 5,874,810 | 76.3 | 44 | 36.3 |
| *vrn1* | Post Double Ridge | 2 | 8,990,467 | 8,144,957 | 73.6 | 45 | 36.5 |
| *vrn1* | Post Double Ridge | 3 | 8,151,534 | 7,518,656 | 74.6 | 44 | 36.5 |
| *vrn1* | Post Double Ridge | 4 | 8,287,973 | 7,593,081 | 74.2 | 45 | 36.3 |
| *vrn1* | Terminal spikelet | 1 | 7,635,487 | 6,996,867 | 74.3 | 43 | 36.5 |
| *vrn1* | Terminal spikelet | 2 | 9,248,399 | 8,497,189 | 74.1 | 44 | 36.6 |
| *vrn1* | Terminal spikelet | 3 | 10,025,819 | 9,043,389 | 73.6 | 44 | 36.6 |
| *vrn1* | Terminal spikelet | 4 | 9,251,175 | 8,571,176 | 74.9 | 43 | 36.5 |
| | **Average** | | **8,009,557** | **7,335,215** | **74.3** | **44.0** | **36.4** |

**Supplemental Table 3**. Differentially Expressed Genes (DEGs) between *vrn1* and *vrn1 ful2* at four different developmental stages of spike development: VEG= vegetative, DR= double ridge, PDR= post double ridge, TS= terminal spikelets. Supplementary File 1 has the complete list of DEGs at FDR 0.05. This table includes DEGs with known functions in inflorescence or flower development in grasses, with their corresponding references and short functional description. DEGs are grouped by expression clusters as described in Figure 1. The X indicates effect on spikelet number per spike or panicle (SNS) or spikelet and floret organ identity (O. ID).

| Gene | Name/Symbol | Sp. | Ref. | Description | Function | |
|---|---|---|---|---|---|---|
| | | | | | SNS | O. ID |
| **Cluster 4** | | | | | | |
| TraesCS2A02G116900<br>TraesCS2B02G136100 | *BRANCHED HEAD1*<br>*COMPOSITUM2*<br>*BRANCHED SILKLESS*<br>*FRIZZY PANICLE* | *Ta*<br>*Hv*<br>*Zm*<br>*Os* | (1)<br>(2)<br>(3) | Mutations in this gene are associated with branched spikes in wheat and with the replacement of spikelets by branches in rice. | x | |
| TraesCS1A02G314200<br>TraesCS1B02G326500 | *MULTI-FLORET SPIKELET1* | *Os* | (4) | Involved in the regulation of spikelet meristem determinacy and floral organ identity in rice. | x | x |
| TraesCS3A02G350600<br>TraesCS3B02G383000 | *LAX PANICLE1*<br>*BARREN STALK1* | *Os*<br>Zm | (5)<br>(6) | Required for the initiation/maintenance of axillary meristems in rice and maize inflorescence (branches and flowers). | | |
| TraesCS2A02G232400<br>TraesCS2B02G250900 | *SPL13* | *Os* | (7) | Involved in the regulation of panicle length, number of primary branches and grain size in rice. | x | |
| TraesCS5A02G319200 | *SIX-ROWED SPIKE 2* | *Hv* | (8) | The barley *vrs2* mutants develops supernumerary spikelets at its base. | x | |
| TraesCS5B02G319600 | *SHORT INTERNODES1* | *Os* | (9) | The rice *shi1* mutant shows reduced tiller number, enhanced culm strength, and increased panicle branch number. | x | |
| TraesCS3A02G247200 | *DWARF TILLER1* | *Os* | (10) | The rice *dtw1* mutant has short internodes. | | |
| **Cluster 5** | | | | | | |
| TraesCS4A02G058900<br>TraesCS4B02G245700 | *OsMADS1*<br>*LEAFY HULL STERILE1* | *Os* | (11) | The rice *lhs1* mutant contains leafy palea and lemmas. | | x |
| TraesCS7A02G122000<br>TraesCS7B02G020800 | *OsMADS5* | *Os* | (12) | LOFSEP-clade genes *OsMADS1*, *OsMADS5*, and *OsMADS34* together regulate floral meristem determinacy and specify spikelet organs identity. | | x |
| TraesCS5A02G286800<br>TraesCS5B02G286100 | *OsMADS8* | *Os* | (13) | Plants affected in both *OsMADS7* and *OsMADS8* show late flowering, homeotic changes of lodicules, stamens and carpels into palea/lemma-like organs, and a loss of floral determinacy. | | x |
| TraesCS7A02G260600<br>TraesCS7B02G158600 | *OsMADS7* | *Os* | (13) | | | |
| TraesCS1A02G264300 | *OsMADS4 / PI1* | *Os* | (14) | These genes control lodicule and stamen development. | | x |
| TraesCS3B02G440200 | *OsMADS2 / PI2* | *OS* | (15) | | | x |
| TraesCS6A02G259000<br>TraesCS6B02G286400 | *OsMADS6*<br>*MOSAIC FLORAL ORGANS1* | *Os* | (16)<br>(17) | Rice *osmads6* mutant has altered palea identity, extra glume-like or mosaic organs, abnormal carpels and loss of floral meristem determinacy.<br>In *mfo1* mutants, the identities of palea and lodicule are disturbed, and mosaic organs were observed and the determinacy of the floral meristem was lost. | | x |
| TraesCS3B02G157500 | *OsMADS3* | *Os* | (18) | A knockout line shows homeotic transformation of stamens into lodicules and ectopic development of lodicules in the second whorl near the palea, carpels develop almost normally. | | x |
| TraesCS1A02G125800 | *OsMADS58* | *Os* | | RNA-silenced lines of *OSMADS58* develop flowers that reiterate a set of floral organs, including lodicules, stamens, and carpel-like organs. | | x |

| | | | | | | |
|---|---|---|---|---|---|---|
| TraesCS3B02G318300 | *OsMADS32* | *Os* | (19) | Rice *OsMADS32* (*CHIMERIC FLORAL ORGANS1*) plays a role in maintaining floral organ identity. | | x |
| TraesCS2A02G312200<br>TraesCS2B02G329000 | *NONSTOP GLUMES1* | *Os* | (20) | Rice *NSG1* plays a pivotal role in maintaining organ identities in the spikelet by repressing the expression of *LHS1*, *DL*, and *MFO1*. | | x |
| TraesCS5A02G134700<br>TraesCS5B02G133700 | *LAX-A* | *Hv* | (21) | Barley mutant *lax-a* phenotype includes extended rachis internodes, broadened base of the lemma awns, thinner grains and homeotic conversion of lodicules into two stamenoid structures. Homolog of *BLADE-ON-PETIOLE1* (*BOP1*) and *BOP2* | | |
| **Cluster 7** (includes *SVP1*, *VRT2* and *SVP3* from this study) | | | | | | |
| TraesCS1A02G154900<br>TraesCS1B02G172100 | *TAW1* | Os | (22) | Gain-of-function mutant prolongs branch formation and increased numbers of spikelets. Upstream SVP-like genes. | x | |
| TraesCS2A02G420900<br>TraesCS2B02G440000 | *NAL1 / SPIKE* | Os | (23) | Natural variant with increased expression are associated with increases in spikelet number, leaf size, root system, and number of vascular bundles. | x | |
| TraesCS1A02G106200<br>TraesCS1B02G123500 | *GA2OX1* | Os | (24) | Ectopic expression of the OsGA2ox1 cDNA in transgenic rice inhibited stem elongation and the development of reproductive organs. | | |
| **Cluster 8** | | | | | | |
| TraesCS5A02G391800<br>TraesCS5B02G396700 | *OsMADS34 / PAP2* | Os | (25) | Rice *pap2-1* mutant shows transformation of early arising spikelets into rachis branches. Rudimentary glumes and sterile lemmas, the outermost organs of the spikelet, elongate into a leafy morphology. | x | x |
| TraesCS5A02G265900 | *SPL17* | Os | (26) | Panicle branching and spikelets are reduced in *SPL17* RNAi plants. | x | |
| TraesCS1B02G448200 | *NECK LEAF1 (NL1)* | Os | (27) | Rice *nl1* mutant shows delay in flowering time and smaller panicles. | x | |
| TraesCS7B02G384000 | *APO1 / WAPO1* | Os | (28)<br>(29) | Rice *APO1* and its wheat ortholog *WAPO1* positively control spikelet number. | x | |
| TraesCS5B02G560300 | *GA20OX1 / GNP1* | Os | (30) | Natural variants with increased expression are associated with increases in grain number. | x | |
| **Cluster 9** | | | | | | |
| TraesCS4B02G042700 | *TB1-1* | Ta | (31)<br>(32) | Increased dosage of TB1 promotes paired spikelet production and delays inflorescence growth.<br>TCP family in wheat. | x | |
| TraesCS5B02G127600 | *RICE CENTRODIALIS1* | Os | (33) | *35S::RCN1* shows delay of transition to the reproductive phase and more branched, denser panicle morphology. | x | |
| TraesCS7A02G229400 | *OsMFT1* | Os | (34) | Overexpressing *OsMFT1* delays heading time and increased spikelets and branches per panicle. | | |
| TraesCS2A02G188500 | *vrs1*<br>*TaVRS1* | Hv<br>Ta | (35)<br>(36) | Loss-of-function mutations result in conversion of two-rowed barley rudimentary lateral spikelets into fertile spikelets in six-rowed barley.<br>Over-expression of *TaVRS1-2B* reduces SNS. | x | |
| **Cluster 10** | | | | | | |
| TraesCS7B02G195200 | OsMFT1 | Os | (34) | Overexpressing *OsMFT1* delays heading time and increases spikelets and branches per panicle. | x | |
| TraesCS4A02G409200<br>TraesCSU02G202000<br>TraesCS2B02G310700 | *RICE CENTRODIALIS2* | Os | (33) | *35S::RCN2* shows delay of transition to the reproductive phase and more branched, denser panicle morphology. | x | |
| TraesCS5A02G001900<br>TraesCS5B02G002300 | *TB1-2* | Ta | (32) | TCP family in wheat. The wheat *TB1* locus is duplicated. | | |
| TraesCS2A02G514000 | *WOX4* | | (37) | *OsWOX4* is as a key regulator at the early stages of leaf development. | | |

#### References Supplemental Table 3

**Supplemental Table 4**. Primers used for gene cloning, genotyping of mutations and transgenic plants, qRT-PCR, and generation of Y2H and Y3H constructs.

| Use & gene | Orientation | Primer sequence 5' to 3' | Res. Enzyme |
|---|---|---|---|
| **Gene cloning** (into Gateway™ pDONR™/Zeo Vector**)** | | | |
| *VRT2* | VRT2-ATTB1 | ggggacaagtttgtacaaaaaagcaggcttcATGGCGCGGGAGAGGCGGGC | |
| | VRT2-ATTB2 | ggggaccactttgtacaagaaagctgggtcTTACTTCCAAGGTAACGCTAG | |
| *SVP1* | SVP1-ATTB1 | ggggacaagtttgtacaaaaaagcaggctatATGGCGCGGGAGCGGAGGGAGA | |
| | SVP1-ATTB2 | ggggaccactttgtacaagaaagctgggtaTTACTTCCACGGAAGGCAGG | |
| *SEP1-2* | SEP1-2-ATTB1 | ggggacaagtttgtacaaaaaagcaggctatATGGGTCGGGGGAAGGTGGAG | |
| (*MADS1*) | SEP1-2-ATTB2 | ggggaccactttgtacaagaaagctgggtaTCATATCCAACCTGCAGATG | |
| *SEP1-4* | SEP1-4-ATTB1 | ggggacaagtttgtacaaaaaagcaggctatATGGGGCGCGGCAAGGTGGAGCT | |
| (*MADS5*) | SEP1-4-ATTB2 | ggggaccactttgtacaagaaagctgggttTCATTCTTTGTTCAACTGATCC | |
| *SEP1-6* | SEP1-6-ATTB1 | ggggacaagtttgtacaaaaaagcaggctatATGGGTCGCGGCAAGGTGGT | |
| (*MADS34*) | SEP1-6-ATTB2 | ggggaccactttgtacaacaaagctgggttCTATGCCATCCATGCAGGCGG | |
| **Genotyping of mutants** | | | |
| *vrt-A2* | Forward | TGAAGTATATACCTGCCTCGG | CAPS(*Bbv*I) |
| | Reverse | ACCTTGTGTTGGAGGTCAGTA | |
| *vrt-B2* | Forward | GGATTGCAGAGGGTGCTTTGTACGAAG | dCAPS(*Mbo*II) |
| | Reverse | CAGAGTCTAAACAAACAATTG | |
| *svp-A1* | Forward | TCAGCTTGCAGAAGCTAGTCTTCGACGAAG | dCAPS(*Mbo*II) |
| | Reverse | TTGAAAAATGACATGTTCACTG | |
| *svp-B1* | Forward | GTTCATGATAGCTGTAAAAAAAT | dCAPS(*Pvu*II) |
| | Reverse | AACCTTAGTCGAAGACTAGCTTCTGCCAGCT | |
| **Transgenic *VRT2* plants** | | | |
| 297 | Forward | TCCCTGAAACTTGCGTTACC | |
| 1064 | Reverse | TCGCTTATTTAAAGGGCGAAT | |
| **qRT-PCR** | | | |
| *VRT2* | Forward | GAGGTGAGGAACTTGACGGA | |
| | Reverse | GAATTGCCGGTCCTTTGTAC | |
| *TaSVP1* [1] | Forward | CACAGGGTGCTTCAGACAAA | |
| | Reverse | TTGGAATCTGGCCTACTTGG | |
| *TaSVP3* [1] | Forward | CCAGCAGATGAGAGGAGAGG | |
| | Reverse | GGCTCTTGGTTTTCAGAACG | |
| *TaSEP1-2* | MADS1-Fw | GGAGCAAGAATTGCAGGATG | |
| (*MADS1*) [2] | MADS1-Rev | GCTASACTGCCCTCCGTCTT | |
| *TaSEP1-4* | MADS5-Fw | GGCGACAAAGAGCCAACAGT | |
| (*MADS5*) [2] | MADS5-Rev | TCCAACATCCTGGCAAGACA | |
| *TaSEP3-1* | MADS7-Fw | CAGTTGGAGGAGAGCAACCA | |
| (*MADS7*) [2] | MADS7-Rev | AAGGGGGTGGAAGAATCCAT | |
| *TaSEP3-2* | MADS8-Fw | CCAACTTGCTCGGCTACGAC | |
| (*MADS8*) [2] | MADS8-Rev | TGCGTTGTTTATCTGCTCCTG | |
| *TaPI-1* | MADS4-Fw | AGATGCTGGAGGAGGAGCAC | |
| (*MADS4*) [2] | MADS4-Rev | CGGCATCTGGGAAGTGAAAT | |
| *TaAP3* | MADS16Fw3 | AAAATGTCGATGCCGCTCTC | |
| (*MADS16*) [2] | MADS16Rev3 | CTCCTGGGAGTGCTTCACCT | |
| *TaSEP1-6* | MADS34-Fw | GCAGCCAGAGCACTTCTTCC | |
| (*MADS34*) [2] | MADS34-Rev | GGCTGGTTCACATCCATGC | |
| *FUL2* [3] | FUL2-FW | CCATACAAAAATGTCACAAGC | |
| | FUL2-Rev | TTCTGCCTCTCCACCAGTTC | |
| *VRN1* [4] | VRN1-Ex5-6-F2 | AAGAAGGAGAGGTCACTGCAGG | |
| | VRN1-Ex8-R37 | GGCTGCACTGCCGCA | |
| *ACTIN* | Actin F | ACCTTCAGTTGCCCAGCAAT | |
| | Actin R | CAGAGTCGAGCACAATACCAGTTG | |

#### Continuation of Supplemental Table 4

| | | |
|---|---|---|
| **Y2H and Y3H constructs** [5] | | |
| *VRN1* | VRN1-NdeI-F | CTGCATATGGGGCGGGGGAAGGTGCAG |
| | VRN1-EcoRI-R | GAATTCCCCGTTGATGTGGCTCACC |
| *FUL2* | FUL2-EcoRI-F | GAATTCATGGGTCGCGGCAAGGTGCAG |
| | FUL2-BamHI-R | GGGATCCGCGTTGAGGTGGCTCAGCATC |
| *FUL3* | FUL3-NdeI-F | CTGCATATGGGGCGCGGGCCGGTGCAG |
| | FUL3-EcoRI-R | GAATTCTCTGTTGCTGATGGTGGAGAG |
| *VRT2* | VRT2-NdeI-F | CCCCATATGGCGCGGGAGAGGCGGGC |
| | VRT2-EcoRI-R | CCCGAATTCCTTCCAAGGTAACGCTAG |
| *SVP1* | SVP1-NedI-F | CCCCATATGGCGCGGGAGCGGAGG |
| | SVP1-EcoRI-R | CCCGAATTCCTTCCACGGGAGGCAGGGCA |
| *SVP3* | SVP3-NdeI-F | AATCATATGGCGGGGAAGAGGGAGAGG |
| | SVP3-EcoRI-R | AAGAATTCCTTCGAGTTGTAGAGTGGTAATC |
| *SEP1-2* | SEP1-2-EcoRI-F | TGTATCGCCGGAATTCATGGGTCGGGGGAAG |
| | SEP1-2-BamHI-R | GCAGGTCGACGGATCCTCATATCCAACCTGCAG |
| *SEP1-4* | SEP1-4-EcoRI-F | TGTATCGCCGGAATTCATGGGGCGCGGCAAGG |
| | SEP1-4-BamHI-R | GCAGGTCGACGGATCCTCATTCTTTGTTCAACT |
| *SEP1-6* | SEP1-6-EcoRI-F | TGTATCGCCGGAATTCATGGGTCGCGGCAAGGTG |
| | SEP1-6-BamHI-R | GCAGGTCGACGGATCCCTATGCCATCCATGCAGG |
| *VRT2* | VRT2-M25-NotI-F | GAAAGGTGGCGGCCGCATGGCGCGGGAGAGGCGG |
| | VRT2-M25-BglII-R | ATCAGCCCGAAGATCTTTACTTCCAAGGTAACGC |

[1] (Li et al., 2019), [2] (Debernardi et al., 2020), [3] (Chen and Dubcovsky, 2012), and [4] (Yan et al., 2006).

[5] Restriction sites for enzyme-based cloning and overlapping vector sequences required for in-fusion cloning are underlined.

#### References Supplemental Table 4


**Chen, A., and Dubcovsky, J.** (2012). Wheat TILLING mutants show that the vernalization gene *VRN1* down-regulates the flowering repressor *VRN2* in leaves but is not essential for flowering. PLoS Genet. **8**: e1003134.

**Debernardi, J.M., Greenwood, J.R., Jean Finnegan, E., Jernstedt, and J., Dubcovsky, J.** (2020). *APETALA 2*-like genes *AP2L2* and *Q* specify lemma identity and axillary floral meristem development in wheat. Plant J. **101**: 171-187.

**Li, C.X., Lin, H.Q., Chen, A., Lau, M., Jernstedt, J., and Dubcovsky, J.** (2019). Wheat *VRN1*, *FUL2* and *FUL3* play critical and redundant roles in spikelet development and spike determinacy. Development. **146**: dev175398.

**Yan, L., Fu, D., Li, C., Blechl, A., Tranquilli, G., Bonafede, M., Sanchez, A., Valarik, M., Yasuda, S., and Dubcovsky, J.** (2006). The wheat and barley vernalization gene *VRN3* is an orthologue of *FT*. Proc. Natl. Acad. Sci. U.S.A. **103**: 19581-19586.

**Supplemental Table 5**. ANOVA tables for Figure 3, 2x2 factorial design with *VRT2* and *SVP1* as factors and WT and mutant alleles as levels.

**Figure 3C. Heading time.**

| Source | DF | Sum of Squares | Mean Square | F Value | Pr > F |
|---|---|---|---|---|---|
| Genotype | 3 | 6192.14 | 2064.05 | 233.48 | <.0001 |
| Error | 36 | 318.26 | 8.84 | | |
| Corrected Total | 39 | 6510.40 | | | |
| R-Square: 0.951116 | | | | | |

| | DF | SS | Mean Square | F Value | Pr > F |
|---|---|---|---|---|---|
| *VRT2* | 1 | 1816.53 | 1816.53 | 205.48 | <.0001 |
| *SVP1* | 1 | 2287.23 | 2287.23 | 258.72 | <.0001 |
| Int. *VRT2* x *SVP1* | 1 | 1082.83 | 1082.83 | 122.49 | <.0001 |

**Figure 3D. Leaf number.**

| Source | DF | Sum of Squares | Mean Square | F Value | Pr > F |
|---|---|---|---|---|---|
| Genotype | 3 | 87.35 | 29.12 | 94.42 | <.0001 |
| Error | 26 | 8.02 | 0.31 | | |
| Corrected Total | 29 | 95.37 | | | |
| R-Square: 0.915926 | | | | | |

| | DF | SS | Mean Square | F Value | Pr > F |
|---|---|---|---|---|---|
| *VRT2* | 1 | 37.50 | 37.50 | 121.61 | <.0001 |
| *SVP1* | 1 | 35.15 | 35.15 | 113.98 | <.0001 |
| Int. *VRT2* x *SVP1* | 1 | 17.81 | 17.81 | 57.77 | <.0001 |

**Figure 3E. Spikelet number per spike (SNS).**

| Source | DF | Sum of Squares | Mean Square | F Value | Pr > F |
|---|---|---|---|---|---|
| Genotype | 3 | 1097.99 | 366.00 | 135.68 | <.0001 |
| Error | 35 | 97.11 | 2.70 | | |
| Corrected Total | 39 | 1195.10 | | | |
| R-Square: 0.918742 | | | | | |

| | DF | SS | Mean Square | F Value | Pr > F |
|---|---|---|---|---|---|
| *VRT2* | 1 | 493.04 | 493.04 | 182.77 | <.0001 |
| *SVP1* | 1 | 313.08 | 313.08 | 116.06 | <.0001 |
| Int. *VRT2* x *SVP1* | 1 | 123.26 | 123.26 | 45.69 | <.0001 |

**Figure 3F. Plant height.**

| Source | DF | Sum of Squares | Mean Square | F Value | Pr > F |
|---|---|---|---|---|---|
| Genotype | 3 | 5897.36 | 1965.79 | 195.01 | <.0001 |
| Error | 36 | 362.90 | 10.08 | | |
| Corrected Total | 39 | 6260.26 | | | |
| R-Square: 0.942032 | | | | | |

| | DF | SS | Mean Square | F Value | Pr > F |
|---|---|---|---|---|---|
| *VRT2* | 1 | 4193.32 | 4193.33 | 415.98 | <.0001 |
| *SVP1* | 1 | 971.67 | 971.67 | 96.39 | <.0001 |
| Int. *VRT2* x *SVP1* | 1 | 100.86 | 100.86 | 10.01 | 0.0032 |

**Supplemental Table 6**. ANOVA tables for Supplemental Figure 3, 2x2 factorial design with *VRT2* or *SVP1* homeologs as factors WT and mutant alleles as levels

**Supplemental Figure 3A. *VRT-A2* x *VRT-B2*. Heading time.**

| Source | DF | Sum of Squares | Mean Square | F Value | Pr > F |
|---|---|---|---|---|---|
| Genotype | 3 | 46.10 | 15.37 | 13.94 | <.0001 |
| Error | 22 | 24.25 | 1.10 | | |
| Corrected Total | 25 | 70.35 | | | |
| R-Square: 0.655276 | | | | | |

| | DF | SS | Mean Square | F Value | Pr > F |
|---|---|---|---|---|---|
| *VRT-A2* | 1 | 21.09 | 21.09 | 19.14 | 0.0002 |
| *VRT-B2* | 1 | 13.50 | 13.50 | 12.25 | 0.0020 |
| Int. *VRT-A2* x *VRT-B2* | 1 | 6.00 | 6.00 | 5.44 | 0.0292 |

**Supplemental Figure 3B. *SVP-A1* x *SVP-B1*. Heading time.**

| Source | DF | Sum of Squares | Mean Square | F Value | Pr > F |
|---|---|---|---|---|---|
| Model | 3 | 64.75 | 21.58 | 3.54 | 0.0285 |
| Error | 26 | 158.72 | 6.10 | | |
| Corrected Total | 29 | 223.47 | | | |
| R-Square: 0.289734 | | | | | |

| | DF | SS | Mean Square | F Value | Pr > F |
|---|---|---|---|---|---|
| *SVP-A1* | 1 | 14.91 | 14.91 | 2.44 | 0.1302 |
| *SVP-B1* | 1 | 9.96 | 9.96 | 1.63 | 0.2129 |
| *SVP-A1* x *SVP-B1* | 1 | 13.69 | 13.69 | 2.24 | 0.1463 |

**Supplemental Figure 3C. *VRT-A2* x *VRT-B2*. Spikelet number per spike (SNS).**

| Source | DF | Sum of Squares | Mean Square | F Value | Pr > F |
|---|---|---|---|---|---|
| Genotype | 3 | 90.38 | 30.13 | 41.21 | <.0001 |
| Error | 22 | 16.08 | 0.73 | | |
| Corrected Total | 25 | 106.46 | | | |
| R-Square: 0.848928 | | | | | |

| | DF | SS | Mean Square | F Value | Pr > F |
|---|---|---|---|---|---|
| *VRT-A2* | 1 | 70.04 | 70.04 | 95.81 | <.0001 |
| *VRT-B2* | 1 | 12.76 | 12.76 | 17.45 | 0.0004 |
| Int. *VRT-A2* x *VRT-B2* | 1 | 3.76 | 3.76 | 5.14 | 0.0335 |

**Supplemental Figure 3D. *SVP-A1* x *SVP-B1*. Spikelet number per spike (SNS).**

| Source | DF | Sum of Squares | Mean Square | F Value | Pr > F |
|---|---|---|---|---|---|
| Model | 3 | 80.73 | 26.91 | 17.82 | <.0001 |
| Error | 26 | 39.27 | 1.51 | | |
| Corrected Total | 29 | 120.00 | | | |
| R-Square: 0.672778 | | | | | |

| | DF | SS | Mean Square | F Value | Pr > F |
|---|---|---|---|---|---|
| *SVP-A1* | 1 | 15.94 | 15.94 | 10.55 | 0.0032 |
| *SVP-B1* | 1 | 36.10 | 36.10 | 23.90 | <.0001 |
| *SVP-A1* x *SVP-B1* | 1 | 0.86 | 0.86 | 0.57 | 0.4572 |

**Supplemental Figure 3E *VRT-A2* x *VRT-B2*. Peduncle length.**

| Source | DF | Sum of Squares | Mean Square | F Value | Pr > F |
|---|---|---|---|---|---|
| Model | 3 | 2541.34 | 847.11 | 158.16 | <.0001 |
| Error | 28 | 149.97 | 5.36 | | |
| Corrected Total | 31 | 2691.31 | | | |

R-Square: 0.944277

| | DF | SS | Mean Square | F Value | Pr > F |
|---|---|---|---|---|---|
| *VRT-A2* | 1 | 980.14 | 980.14 | 183.00 | <.0001 |
| *VRT-B2* | 1 | 982.35 | 982.35 | 183.41 | <.0001 |
| Int. *VRT-A2* x *VRT-B2* | 1 | 578.85 | 578.85 | 108.07 | <.0001 |

**Supplemental Figure 3F *SVP-A1* x *SVP-B1*. Peduncle length.**

| Source | DF | Sum of Squares | Mean Square | F Value | Pr > F |
|---|---|---|---|---|---|
| Model | 3 | 615.63 | 205.21 | 33.84 | <.0001 |
| Error | 26 | 157.66 | 6.06 | | |
| Corrected Total | 29 | 773.29 | | | |

R-Square: 0.796117

| | DF | SS | Mean Square | F Value | Pr > F |
|---|---|---|---|---|---|
| *SVP-A1* | 1 | 19.48 | 19.48 | 3.21 | 0.0847 |
| *SVP-B1* | 1 | 342.50 | 342.50 | 56.48 | <.0001 |
| Int. *SVP-A1* x *SVP-B1* | 1 | 36.41 | 36.41 | 6.01 | 0.0213 |

**Supplemental Figure 3G *VRT-A2* x *VRT-B2*. Spikelet number per spike (SNS). Field.**

| Source | DF | Sum of Squares | Mean Square | F Value | Pr > F |
|---|---|---|---|---|---|
| Genotype | 3 | 11.63 | 3.88 | 8.97 | <.0001 |
| Error | 56 | 24.20 | 0.43 | | |
| Corrected Total | 59 | 35.82 | | | |

R-Square: 0.324546

| | DF | SS | Mean Square | F Value | Pr > F |
|---|---|---|---|---|---|
| *VRT-A2* | 1 | 3.96 | 3.96 | 9.17 | 0.0037 |
| *VRT-B2* | 1 | 3.54 | 3.54 | 8.18 | 0.0059 |
| Int. *VRT-A2* x *VRT-B2* | 1 | 0.39 | 0.39 | 0.89 | 0.3490 |

**Supplemental Figure 3H *VRT-A2* x *VRT-B2*. Plant height. Field.**

| Source | DF | Sum of Squares | Mean Square | F Value | Pr > F |
|---|---|---|---|---|---|
| Genotype | 3 | 10065.21 | 3355.07 | 698.44 | <.0001 |
| Error | 56 | 269.01 | 4.80 | | |
| Corrected Total | 59 | 10334.22 | | | |

R-Square: 0.973969

| | DF | SS | Mean Square | F Value | Pr > F |
|---|---|---|---|---|---|
| *VRT-A2* | 1 | 2672.46 | 2672.46 | 556.34 | <.0001 |
| *VRT-B2* | 1 | 2714.15 | 2714.15 | 565.01 | <.0001 |
| Int. *VRT-A2* x *VRT-B2* | 1 | 1985.35 | 1985.35 | 413.30 | <.0001 |

**Supplemental Table 7**. Effects of *vrt2* and *svp1* mutations on heading time, spikelet number per spike, stem length, and leaf number in growth chambers under LD conditions. Experiment 1 compared *vrt-A2* (n = 8), *vrt-B2* (n = 6), *vrt2* (n = 8) and wild type sister lines (n = 4). Experiment 2 compared *svp-A1* (n = 6), *svp-B1* (n = 4), *svp1* (n = 15) and wild type sister lines (n = 5). Experiment 3 compared *svp1* (n = 9), *vrt2* (n = 9), *vrt2 svp1* (n = 13) and wild type sister lines (n = 9). In Experiments 1 and 2 main effects are the differences between least square means for the mutant allele minus the WT allele (*P* values are from the 2 x 2 factorial ANOVA). Significant differences between the mutants and the WT are in blue if the mutant increases the value of the trait relative to the WT and in red if the mutants decreased the values (*P* values are from the Dunnett tests). Stem nodes are numbered from the peduncle (-1 is the closest internode to the peduncle and -4 the most basal node).

| Exp. | Effect | HD days | SNS | Peduncle cm | -1 inter-node cm | -2 inter-node cm | -3 inter-node cm | -4 inter-node cm | Leaf No. |
|---|---|---|---|---|---|---|---|---|---|
| 1 | *VRT-A2* Main | 1.9 *** | 3.4 *** | -11.1 *** | 2.1 *** | 0.2 ns | -0.1 ns | -0.2 ns | NA |
| 1 | *VRT-B2* Main | 1.5 ** | 1.5 ** | -11.1 *** | 0.3 ns | -0.5 * | -0.3 ns | -0.3 ns | NA |
| 1 | *vrt2* - WT | 3.4 *** | 4.9 *** | -22.1 *** | 2.4 *** | -0.3 ns | -0.4 ns | -0.5 ns | NA |
| 2 | *SVP-A1* Main | 1.2 ns | 1.7 ** | -1.8 ns | 1.1 * | 0.1 ns | -0.2 ns | -0.9 ns | NA |
| 2 | *SVP-B1* Main | 1.3 ns | 2.5 *** | -7.7 *** | 1.3 * | 0.7 * | -0.1 ns | -1.3 * | NA |
| 2 | *svp1* - WT | 2.9 ns | 2.9 *** | -9.5 *** | 2.4 ** | 0.8 ns | -0.3 ns | -2.2 ** | NA |
| 3 | *vrt2* - WT | 3.1 * | 3.6 *** | -20.1 *** | 1.9 * | 0.1 ns | 0.2 ns | 0.5 ns | 0.7 ns |
| 3 | *svp1* - WT | 4.8 *** | 2.1 * | -8.5 * | -0.6 ns | -0.2 ns | -0.7 ns | -0.7 ns | 0.6 ns |
| 3 | *vrt2 svp1* - WT | 29.0 *** | 13.0 *** | -25.0 *** | -4.0 *** | -0.8 ns | -1.5 * | 0.1 ns | 4.4 *** |

**Supplemental Table 8**. ANOVA tables for Figure 7, 2 x 2 factorial design with *FUL2* and *UBI::VRT2* weak transgenic T#8 as factors and alleles as levels.

**Figure 7A. Stem length**

| Source | DF | Sum of Squares | Mean Square | F Value | Pr > F |
|---|---|---|---|---|---|
| Genotype | 3 | 829.03 | 276.34 | 55.17 | <.0001 |
| Error | 20 | 100.17 | 5.01 | | |
| Corrected Total | 23 | 929.20 | | | |
| R-Square: 0.892 | | | | | |

| | DF | SS | Mean Square | F Value | Pr > F |
|---|---|---|---|---|---|
| T#8 | 1 | 3.36 | 3.36 | 0.67 | 0.4224 |
| *FUL2* | 1 | 793.00 | 793.00 | 158.33 | <.0001 |
| Int. T#8 x *FUL2* | 1 | 15.99 | 15.99 | 3.19 | 0.0892 |

**Figure 7B. Spikelet number per spike (SNS)**

| Source | DF | Sum of Squares | Mean Square | F Value | Pr > F |
|---|---|---|---|---|---|
| Genotype | 3 | 189.77 | 63.26 | 159.51 | <.0001 |
| Error | 22 | 8.72 | 0.40 | | |
| Corrected Total | 25 | 198.50 | | | |
| R-Square: 0.956 | | | | | |

| | DF | SS | Mean Square | F Value | Pr > F |
|---|---|---|---|---|---|
| T#8 | 1 | 76.04 | 76.04 | 191.74 | <.0001 |
| *FUL2* | 1 | 74.01 | 74.01 | 186.61 | <.0001 |
| Int. T#8 x *FUL2* | 1 | 11.01 | 11.01 | 27.76 | <.0001 |

**Figure 7C. Glume length**

| Source | DF | Sum of Squares | Mean Square | F Value | Pr > F |
|---|---|---|---|---|---|
| Genotype | 3 | 2.92 | 0.97 | 53.36 | <.0001 |
| Error | 23 | 0.42 | 0.02 | | |
| Corrected Total | 26 | 3.34 | | | |
| R-Square: 0.874 | | | | | |

| | DF | SS | Mean Square | F Value | Pr > F |
|---|---|---|---|---|---|
| T#8 | 1 | 1.66 | 1.66 | 91.12 | <.0001 |
| *FUL2* | 1 | 0.76 | 0.76 | 41.62 | <.0001 |
| Int. T#8 x *FUL2* | 1 | 0.26 | 0.26 | 14.27 | 0.0010 |

**Figure 7D. Lemma length**

| Source | DF | Sum of Squares | Mean Square | F Value | Pr > F |
|---|---|---|---|---|---|
| Genotype | 3 | 2.97 | 0.99 | 61.02 | <.0001 |
| Error | 23 | 0.37 | 0.02 | | |
| Corrected Total | 26 | 3.34 | | | |
| R-Square: 0.888 | | | | | |

| | DF | SS | Mean Square | F Value | Pr > F |
|---|---|---|---|---|---|
| T#8 | 1 | 1.24 | 1.24 | 76.26 | <.0001 |
| *FUL2* | 1 | 0.92 | 0.92 | 56.45 | <.0001 |
| Int. T#8 x *FUL2* | 1 | 0.41 | 0.41 | 25.06 | <.0001 |

**Supplemental Table 9**. Summary of Y2H interactions (lower triangular table) and BiFC interactions (upper triangular table) tested among the SQUAMOSA, SVP and SEP MADS-box proteins.

| | VRN1 | FUL2 | FUL3 | VRT2 | SVP1 | SVP3 | SEP1-2 | SEP1-4 | SEP1-6 |
|---|---|---|---|---|---|---|---|---|---|
| VRN1 | Y2H- | NA | NA | BiFC+[1] | BiFC+ | NA | BiFC+ | BiFC- | (BiFC-)[2] |
| FUL2 | Y2H+ | Y2H+ | NA | BiFC+ | BiFC+ | NA | BiFC+ | (BiFC-)[2] | (BiFC-)[2] |
| FUL3 | Y2H- | Y2H+ | Y2H+[3] | BiFC+ | BiFC+ | NA | BiFC+ | BiFC- | BiFC- |
| VRT2 | Y2H+ | Y2H+[2] | Y2H+ | Y2H- | NA | NA | NA | NA | NA |
| SVP1 | Y2H+ | Y2H+ | Y2H+ | Y2H- | Y2H+ | NA | NA | NA | NA |
| SVP3 | Y2H- | Y2H+ | Y2H- | Y2H- | Y2H- | Y2H- | NA | NA | NA |
| SEP1-2 | Y2H+ | Y2H+ | Y2H+ | Y2H- | Y2H+ | Y2H- | NA | NA | NA |
| SEP1-4 | Y2H+ | Y2H+ | Y2H+ | Y2H- | Y2H- | Y2H- | NA | NA | NA |
| SEP1-6 | Y2H+ | Y2H+ | Y2H+ | Y2H- | Y2H- | Y2H+[3] | NA | NA | NA |

[1] The interaction VRT2-VRN1 was confirmed by CoIP and luciferase assays in previous study (Xie et al., 2019).

[2] (BiFC-) in parentheses indicates no clear nuclear signal but bright fluorescent aggregates outside the nucleus.

[3] Weak interaction.

#### Reference Supplemental Table 9

**Xie, L., Zhang, Y., Wang, K., Luo, X., Xu, D., Tian, X., Li, L., Ye, X., Xia, X., Li, W., Yan, L., and Cao, S.** (2019). *TaVrt2*, an SVP-like gene, cooperates with *TaVrn1* to regulate vernalization-induced flowering in wheat. *New Phytol*. doi: 10.1111/nph.16339
